# An Expanded Registry of Candidate cis-Regulatory Elements for Studying Transcriptional Regulation

**DOI:** 10.1101/2024.12.26.629296

**Authors:** Jill E. Moore, Henry E. Pratt, Kaili Fan, Nishigandha Phalke, Jonathan Fisher, Shaimae I. Elhajjajy, Gregory Andrews, Mingshi Gao, Nicole Shedd, Yu Fu, Matthew C Lacadie, Jair Meza, Mohit Ganna, Eva Choudhury, Ross Swofford, Nina P. Farrell, Anusri Pampari, Vivekanandan Ramalingam, Fairlie Reese, Beatrice Borsari, Michelle Yu, Eve Wattenberg, Marina Ruiz-Romero, Milad Razavi-Mohseni, Jinrui Xu, Timur Galeev, Michael A. Beer, Roderic Guigó, Mark Gerstein, Jesse Engreitz, Mats Ljungman, Timothy E. Reddy, Michael P. Snyder, Charles B Epstein, Elizabeth Gaskell, Bradley E Bernstein, Diane E. Dickel, Axel Visel, Len A. Pennacchio, Ali Mortazavi, Anshul Kundaje, Zhiping Weng

## Abstract

Mammalian genomes contain millions of regulatory elements that control the complex patterns of gene expression. Previously, The ENCODE consortium mapped biochemical signals across many cell types and tissues and integrated these data to develop a Registry of 0.9 million human and 300 thousand mouse candidate cis-Regulatory Elements (cCREs) annotated with potential functions^1^. We have expanded the Registry to include 2.35 million human and 927 thousand mouse cCREs, leveraging new ENCODE datasets and enhanced computational methods. This expanded Registry covers hundreds of unique cell and tissue types, providing a comprehensive understanding of gene regulation. Functional characterization data from assays like STARR-seq, MPRA, CRISPR perturbation, and transgenic mouse assays now cover over 90% of human cCREs, revealing complex regulatory functions. We identified thousands of novel silencer cCREs and demonstrated their dual enhancer/silencer roles in different cellular contexts. Integrating the Registry with other ENCODE annotations facilitates genetic variation interpretation and trait-associated gene identification, exemplified by discovering *KLF1* as a novel causal gene for red blood cell traits. This expanded Registry is a valuable resource for studying the regulatory genome and its impact on health and disease.

## Introduction

Mammalian genomes are extensive repositories of DNA-encoded instructions that control cellular functions through complex regulatory mechanisms. Central to this regulation are cis-regulatory elements (CREs), which are non-coding DNA sequences that control the transcription of nearby genes. Usually associated with open chromatin and specific histone modifications, CREs contain binding sites for transcription factors and other chromatin-associated proteins that interact with one another and transcriptional machinery to regulate gene expression^2,3^. Understanding the biological contexts and functions of CREs is essential for deciphering genome function and its impact on human health and disease^4–7^.

Over the years, the ENCODE project has made major contributions towards our understanding of gene regulation by systematically identifying and annotating functional elements across the human and mouse genomes^8–11^. Performing tens of thousands of high-throughput assays such as DNase-seq^12,13^, ATAC-seq^14^, RNA-seq^15,16^, and ChIP-seq for histone modifications^17^ and transcription factors^18,19^, the ENCODE consortium has comprehensively mapped a diverse set of biochemical signatures and used these signatures to annotate CREs. One major result of ENCODE Phase 3 was the creation of the ENCODE Registry of candidate cis-Regulatory Elements (cCREs), a resource that provides insights into the roles of cCREs in gene regulation and that serves as a powerful aid for researchers to study the genetic basis of complex traits and diseases.^1^

In this work, we have significantly expanded the Registry of cCREs to include 2.35 million human and 927 thousand mouse elements. This expansion leverages new datasets produced during ENCODE Phase 4^20^ and improved computational methods, making the Registry one of the most extensive repositories of CREs available. The updated Registry spans hundreds of unique cell and tissue types, enhancing our understanding of gene regulation across a broad range of biological contexts.

In addition to an increase in the number of cCREs, the Registry now includes functional characterization data^21^ for over 90% of human cCREs from assays such as genome-wide STARR-seq^22,23^, MPRA^24–26^, CRISPR perturbation^27,28^, and transgenic mouse assays. These functional assays provide deeper insights into how cCRE sequences impact their function, highlighting the regulatory complexity within the genome. We have also identified thousands of candidate silencer cCREs, many of which can function as enhancers in other cellular contexts, with the majority being novel to this study.

Finally, integrating the updated Registry with other ENCODE annotations facilitates the interpretation of genetic variation and the identification of trait-associated genes. This integration has practical implications, as demonstrated by our discovery of *KLF1* as a novel causal gene for red blood cell traits. These advancements underscore the utility of the ENCODE Registry as an invaluable resource for genomic research, enhancing our ability to study the regulatory landscape of the genome and its impact on human health and disease.

## Results

### The expanded ENCODE Registry comprises 2.35 million human and 927 thousand mouse cCREs

During ENCODE4, we leveraged new datasets and improved computational methods to broaden the scope and scale of the Registry of cCREs. The current Registry now encompasses 2,348,854 human cCREs and 926,843 mouse cCREs—a threefold increase compared to the ENCODE3 Registry, establishing it as one of the most extensive repositories of CREs currently available (**Figure 1, Supplementary Note 1, Supplementary Data 1, 2**).

**Figure 1.**
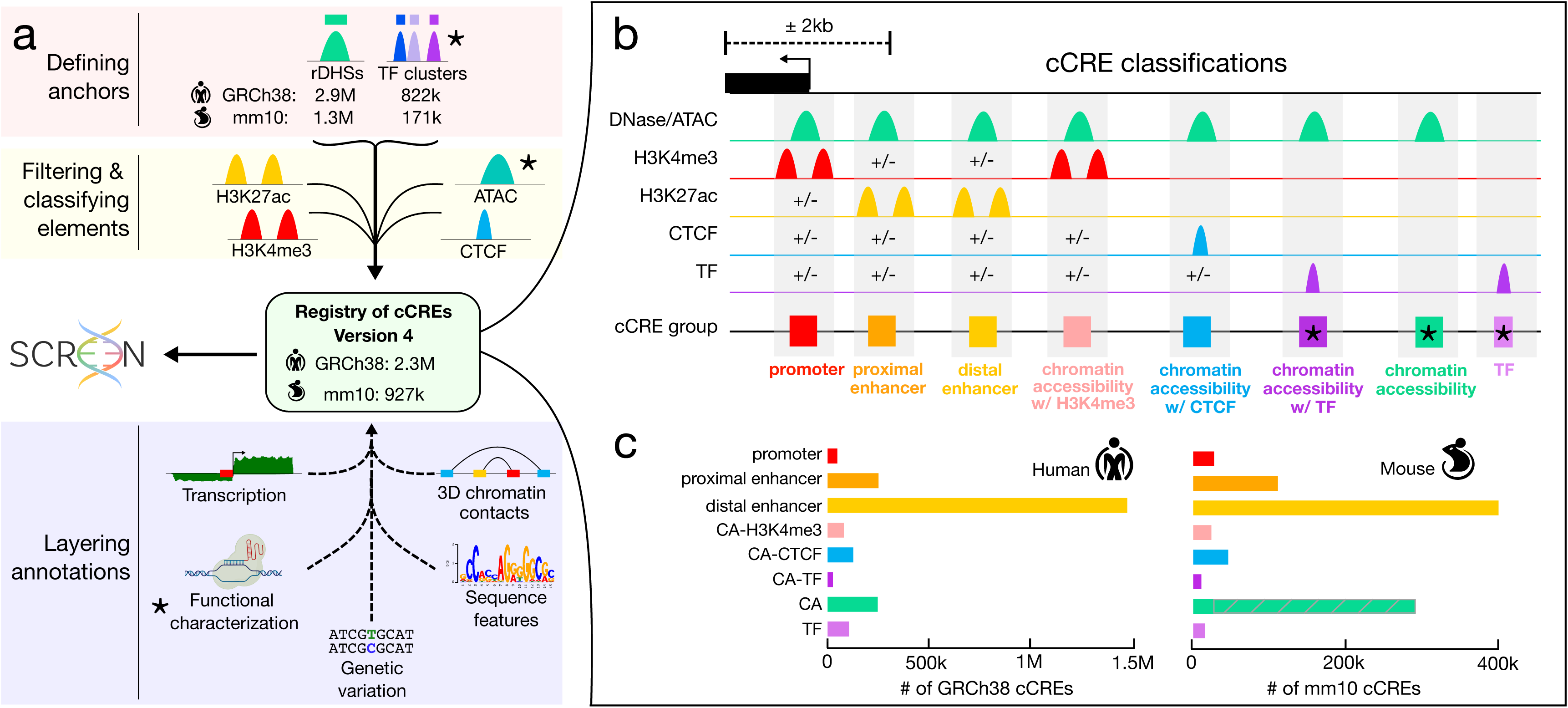
The updated Registry of candidate cis-Regulatory elements. **a**, Schematic of the pipeline used to make Version 4 of the Registry of cCREs. We define element anchors by generating representative DHSs (rDHSs) and transcription factor clusters. Element anchors are scored with H3K4me3, H3K27ac, and CTCF ChIP-seq and ATAC-seq signals (yellow box) and classified according to the scheme in b. This results in 2.3 million cCREs in the human genome and 927 thousand in the mouse genome. We supplement the Registry with additional ENCODE Encyclopedia annotations including transcription quantifications, 3D chromatin contacts, functional characterization measurements, sequence features, and genetic variation (blue box). The Registry of cCREs and all layered annotations are housed in our web portal SCREEN. New components of the pipeline are denoted by stars. **b**, Overview of our cCRE classification scheme. cCREs are classified based on their patterns of biochemical signals (chromatin accessibility in green, H3K4me3 in red, H3K27ac in yellow, CTCF in blue, transcription factor in purple) and distance from annotated TSSs. High signals are denoted by peaks. A +/- symbol indicates that the corresponding signal may or may not be present and its presence does not impact classification. New categories of elements are denoted by stars. **c**, Bar graphs depicting the number of cCREs annotated in each class for human (left) and mouse (right). The gray hatched bar indicates an upper bound for the number of CA cCREs in mouse that would be classified as enhancers if H3K27ac data were available.

This expansion reflects our incorporation of newly generated ENCODE4 data as well as data from new types of assays—in total, 5,712 human and 758 mouse experiments (**Supplementary Table 1**). In this version of the Registry, we continued to anchor cCREs on nucleosome-sized representative DNase Hypersensitivity Sites (rDHSs)^1^ derived from DNase-seq data^29,30^ (**Supplementary Note 1.1**). To annotate the plausible biochemical function of a cCRE, we used ChIP-seq data of two histone modifications (H3K4me3 and H3K27ac, marking active promoters and enhancers, respectively) and the insulator-binding protein CTCF (*Supplementary Methods*). New to this version of the Registry, we also incorporated 2,509 transcription factor ChIP-seq experiments from human^31^ and 167 from mouse to define 105,286 human and 15,238 mouse cCREs in regions with low chromatin accessibility (**Supplementary Note 1.1, Supplementary Figure 1, Supplementary Table 1 c, d, Supplementary Table 2**). We also used ATAC-seq data to annotate chromatin accessibility in biosamples lacking DNase-seq data. Finally, we made technical improvements to our pipeline to define 24,160 human and 40,537 mouse cCREs in recently duplicated genomic regions (**Supplementary Note 1.2, Supplementary Figure 2, Supplementary Table 3, Supplementary Data 3, 4**).

The number of human DNase-seq and H3K4me3, H3K27ac, and CTCF ChIP-seq experiments increased by 2.3 fold from ENCODE3 to ENCODE4. This doubled the number of biosamples—unique tissues, cell types, and cellular states—represented in the Registry from 839 to 1,679. These 1,679 biosamples were from a diverse array of biological contexts, covering 42 different human organs and tissues (**Extended Data Figure 1a**). The majority of these biosamples comprise primary cells and tissues, but also include *in vitro* differentiated cells, organoids, and cell lines (**Extended Data Figure 1b**). Annotation of cCREs in such a wide variety of biosample types is useful for designing and interpreting experiments in more amenable systems—such as cell lines and organoids—and then evaluating their biological relevance in the primary biosamples.

Coordinated data production during ENCODE4 led to a nearly seven-fold increase of human biosamples with all four experiments (DNase, H3K4me3, H3K27ac and CTCF), from 25 in ENCODE3 to 170 in ENCODE4 (**Extended Data Figure 1c**). This *Core Collection* of 170 biosamples (**Supplementary Table 1**) allows us to thoroughly annotate cCREs across diverse cell and tissue types. The remaining human biosamples have chromatin accessibility (DNase or ATAC) data but are only partially covered by ChIP-seq of the other three marks (*Partial Data Collection*, with 1154 biosamples) or have ChIP-seq data but lack chromatin accessibility (*Ancillary Data Collection*, with 354 biosamples), permitting partial cCRE classification (**Supplementary Note 1.3, Supplementary Table 1).**

### cCREs are classified into putative functional categories based on biochemical signatures and genomic context

We previously classified cCREs into putative functional categories based on their genomic distance from annotated transcription start sites (TSSs) and combinations of biochemical signals^1^. Here, we expanded this classification scheme to include eight classes of cCREs (**Figure 1b, c, Supplementary Note 1.4**):

- <u>Promoter-like signatures (promoter)</u> must fall within 200 bp of a TSS and have high chromatin accessibility and H3K4me3 signals.
- <u>TSS-proximal enhancer-like signatures (proximal enhancer)</u> have high chromatin accessibility and H3K27ac signals and are within 2 kb of an annotated TSS. If they are within 200 bp of a TSS, they must also have low H3K4me3 signal.
- <u>TSS-distal enhancer-like signatures (distal enhancer)</u> have high chromatin accessibility and H3K27ac signals and are farther than 2 kb from an annotated TSS.
- <u>Chromatin accessibility + H3K4me3 (CA-H3K4me3)</u> have high chromatin accessibility and H3K4me3 signals but low H3K27ac signals and do not fall within 200 bp of a TSS.
- <u>Chromatin accessibility + CTCF (CA-CTCF)</u> have high chromatin accessibility and CTCF signals but low H3K4me3 and H3K27ac signals.
- <u>Chromatin accessibility + transcription factor (CA-TF)</u> have high chromatin accessibility, low H3K4me3, H3K27ac, and CTCF signals, and are bound by a transcription factor.
- <u>Chromatin accessibility (CA)</u> have high chromatin accessibility and low H3K4me3, H3K27ac, and CTCF signals.
- <u>Transcription factor (TF)</u> have low chromatin accessibility, low H3K4me3, H3K27ac, and CTCF signals and are bound by a transcription factor.

Like the ENCODE3 version of the Registry^1^, we performed the above cCRE classification across all cell types (cell type-agnostic) and in specific cell types (**Supplementary Note 1.4**).

Specifically, our expanded classification scheme resulted in 1,121,741 new cCREs for human and 478,302 for mouse. These newly included cCREs were enriched for evolutionary conservation, functional activity, and chromatin accessibility in previously underrepresented cell types (**Supplementary Note 1.5, Supplementary Figure 3a-c, Supplementary Table 4a**). We also compared ENCODE4 cCREs with two recent studies that reported a moderate overlap between their brain-specific regulatory elements and an earlier version of our Registry^32,33^. We observed a large increase in the percentage of overlapping elements, specifically due to cCREs with high chromatin accessibility in neural cell types and brain tissues (**Supplementary Note 1.5, Supplementary Figure 3d,e, Supplementary Table 4b,c**). These results demonstrate that the ENCODE4 Registry is comprehensive while still maintaining biological specificity.

To better understand how our cCRE classification scheme related to DNA sequence, we trained cell type-specific variational autoencoders on the sequences of the cCREs annotated in three cell types (**Extended Data Figure 2a, Supplementary Note 1.6, Supplementary Figure 4**). In each cell type, promoters and distal enhancers segregated across the first dimension (**Extended Data Figure 2b, Supplementary Figure 4**); this dimension strongly correlated with the percentage of guanine and cytosine (GC) nucleotides of the sequences (R = 0.92, **Extended Data Figure 2c**) and differed significantly between promoters of protein-coding versus lncRNA genes and among distal enhancers overlapping novel TSSs or expressing eRNAs (**Extended Data Figure 2d**). These results suggest that GC content is one of the primary sequence features differentiating transcribed cCREs and that our current classification scheme is able to detect such differences.

It is worth noting that the smaller number of mouse cCREs reflects ENCODE’s emphasis on generating data from human samples. Additionally, because ENCODE4 primarily focused on measuring chromatin accessibility and gene expression in mouse biosamples, rather than histone modifications, the mouse Registry contains a higher percentage of CA cCREs (**Figure 1c**). We estimate that up to 65% of these elements could have been classified as enhancer cCREs had H3K27ac data been available (**Supplementary Note 1.7**).

### Testing cCRE activity using functional assays

Having defined and annotated the putative functions of cCREs using chromatin signatures, we aimed to evaluate their functional activities. ENCODE4 tested the activities of millions of regions in the human genome using four types of functional assays^21^—genome-wide STARR-seq assays^23,31^, massively parallel reporter assays (MPRA)^24–26^, CRISPR perturbation assays^27,28^, and transgenic mouse assays (**Fig. 2a**). Nearly all of the human cCREs (93%) were tested by at least one assay in at least one cellular context (**Figure 2a, Supplementary Note 2, Supplementary Table 5**). Across all assays, cCREs were much more likely to be functionally active than non-cCRE genomic regions, and cCREs with active chromatin signatures in a cell type more frequently tested positive in the same cell type than other cCREs (**Supplementary Figure 5a-f**). This cell type-specificity was most dramatic for CRISPR perturbation experiments that disrupt elements in their native chromatin context (37-fold enrichment, Fisher’s Exact Test, p < 2.2 × 10^−16^**, Supplementary Figure 5f**) compared to methods such as MPRAs that evaluate element activity episomally (2-fold enrichment, Fisher’s Exact Test, p < 2.2 × 10^−16^, **Supplementary Figure 5d**).

**Figure 2.**
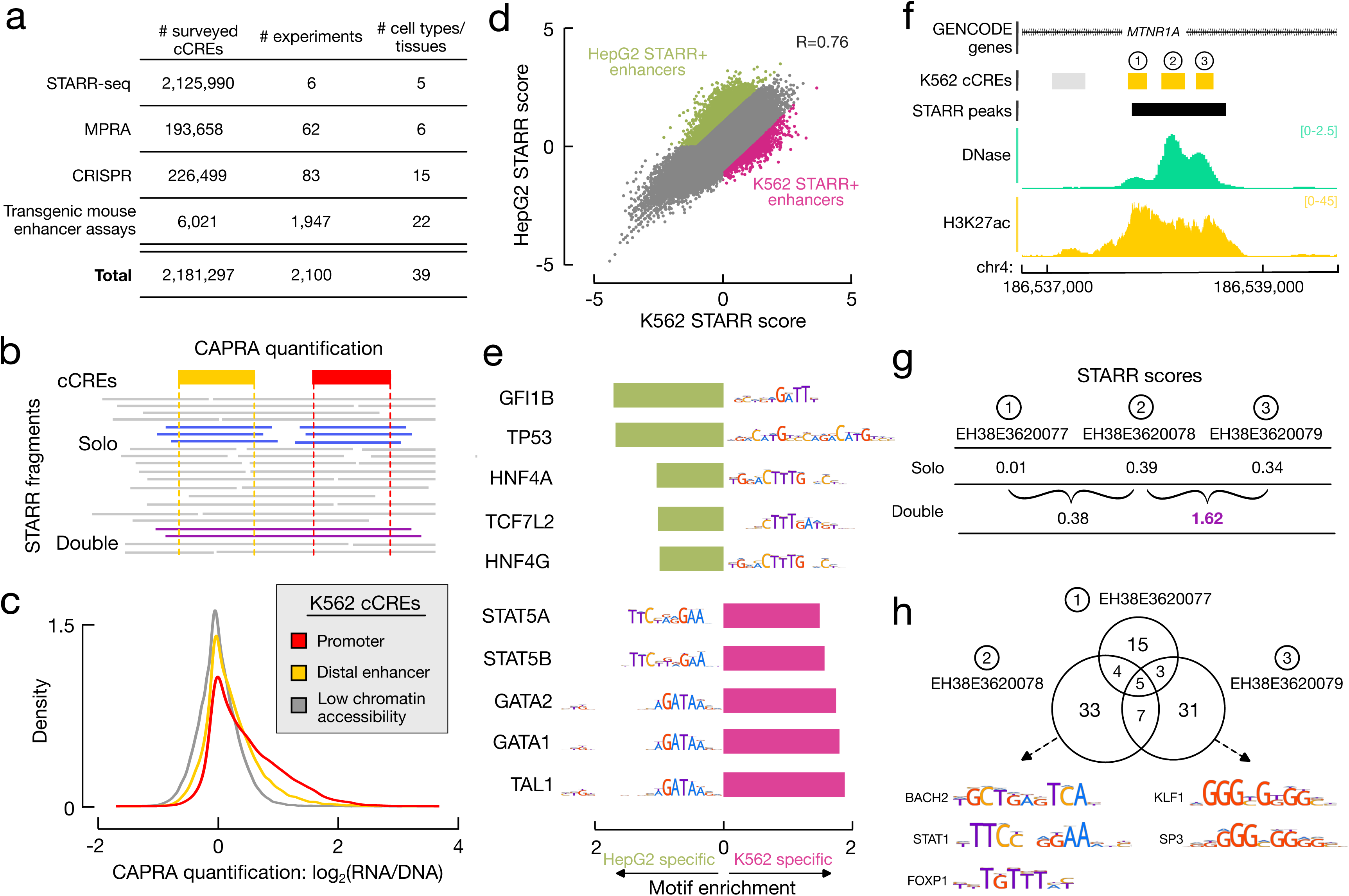
Functional characterization of the Registry of cCREs. **a**, Summary of cCREs tested by ENCODE4 functional characterization assays. **b**, Schematic of the CAPRA quantification method which utilizes solo fragments (overlapping single cCREs in their entirety, blue) and double fragments (overlapping two cCREs in their entirety, purple). **c**, Density plot showing the distributions of CAPRA quantifications in K562 cells for K562 promoter (red), distal enhancer (yellow) or low chromatin accessibility (gray) cCREs. **d**, Scatterplot of CAPRA quantifications for distal enhancer cCREs in K562 (x-axis) and HepG2 (y-axis). Color of points indicates cCREs with enriched activity (STARR+) in K562 (pink) or HepG2 (green). **e**, Barplots of motif enrichment for HepG2 (green) or K562 (pink) STARR+ distal enhancers (as defined in **d**). Top five motifs are shown for each group of cCREs along with their corresponding logo. **f**, Genome browser view of three distal enhancer cCREs (denoted by 1-3) in the *MTNR1A* intron with DNase (green) and H3K27ac (yellow) signals in K562. A STARR-seq peak call is shown in black. **g**, CAPRA quantifications for the three enhancers shown in **f**: EH38E3620077 (1), EH38E3620078 (2) and EH38E3620079 (3) using solo fragments (top) and double fragments (bottom). High quantifications are denoted in purple (*p* = 0.03). **h**, Overlap of common K562 transcription factor motifs at the three enhancers in **f** and **g.** Representative motif logos for EH38E3620078 and EH38E3620079 are shown.

Among the four types of assays, whole genome STARR-seq had the highest throughput, testing 2.2 million cCREs; thus, we focused subsequent analyses on this assay. Because the genomic regions tested by STARR-seq were generated by random shearing, these regions had varying sizes (up to several thousand base pairs) and could contain multiple cCREs, which are 150-350 base pairs long. To delineate the STARR-seq activity scores (STARR scores) of individual cCREs, we developed a novel method called CRE-centric Analysis and Prediction of Reporter Assays (CAPRA). To isolate the impact of neighboring cCREs, we first quantified the STARR score of a cCRE using the “solo” STARR-seq fragments, which overlap the cCRE in its entirety and do not overlap any other cCREs (**Figure 2b**). We then use “double” STARR-seq fragments, which overlap two neighboring cCREs in their entirety, to assess the interaction between the cCREs. By first determining fragment counts across the input DNA and the output RNA libraries, CAPRA calculates the *p*-value of RNA to DNA ratio for each cCRE using DESeq2^34^ (see **Methods**). Due to the strict overlap requirements, solo and double fragments only compose 8-12% of a STARR-seq library; however, our approach still characterizes 75-87% of the 2.35 million human cCREs in each STARR-seq experiment (**Supplementary Figure 5g, Supplementary Table 5a, Supplementary Data 5**).

We first studied cell type-specific functional activity of cCREs by comparing STARR scores for cCREs in K562 cells vs. HepG2 cells. When we stratified cCREs by their class, we found that promoter cCREs were more likely to have consistent STARR scores across cell types compared to distal enhancer cCREs (**Supplementary Figure 5h, i)**. These findings were concordant with our previous results^1^ and the general understanding that enhancers tend to be more cell type-specific than promoters^3,35^.

To determine what sequence features were responsible for the cell type-specific activity of enhancer cCREs, we investigated which transcription factor motifs were enriched in the distal enhancer cCREs with differentially high STARR scores in K562 cells (K562 STARR+) versus HepG2 cells (HepG2 STARR+) (**Figure 2d**). Both the K562 and HepG2 STARR+ cCREs were enriched for cell type-relevant transcription factor motifs and were more likely to be annotated as enhancers in their respective cell types (**Figure 2e, Supplementary Figure 6, Supplementary Table 6a**). K562 STARR+ cCREs were enriched for transcription factors related to hematopoiesis, such as STAT5A/B^36^, TAL1^37^, and GATA family factors GATA1 and GATA2^38^. HepG2 STARR+ cCREs were enriched for hepatocyte nuclear factors HNF4A and HNF4G along with TCF7L2, which is known to co-localize with these transcription factors in HepG2^39^. When we considered two other cell types with STARR-seq data, HCT116 and MCF-7, STARR+ enhancer cCREs enriched for the respective hematopoietic or hepatocyte transcription factor motifs were also much more likely to have cell type-specific STARR activity (**Extended Data Figure 3a**). This cell type-specific activity was concordant with the expression of these transcription factors in the corresponding cell types (**Extended Data Figure 3b**).

We also observed enrichment for two transcription factor motifs in HepG2 STARR+ enhancers that were initially unexpected, TP53 and GFI1B. HepG2 STARR+ enhancers with TP53 motifs had low STARR scores in K562, but high STARR scores in HCT116 and MCF-7 (**Extended Data Figure 3a**). This is because K562 does not have functional TP53,^40^ unlike the other three assayed cell lines^41–44^. These results underscore the importance of biosample selection, as the disruption of regulatory mechanisms in a cancer cell line can impact the interpretation of biological data obtained using that cell line. Additionally, HepG2 STARR+ enhancers with GFI1B motifs had moderate to high STARR scores in HCT116 and MCF-7 but lower than baseline activity in K562 (**Extended Data Figure 3a**). GFI1B is a transcriptional repressor expressed in erythrocyte progenitors^45^ and K562 cells (**Extended Data Figure 3b**). These results suggest that these genomic regions are repressing transcription in the STARR-seq assay in K562 cells and that our method can identify elements with repressive or silencing activities.

In addition to characterizing the activity of individual cCREs, our CAPRA method quantified the activity of 335,909 pairs of cCREs using whole-genome K562 STARR-seq data (**Supplementary Data 6**). Combined activity levels were generally correlated with the averaged activity of the individual cCREs, and this correlation increased as more stringent filters were applied to the data (**Supplementary Figure 7a-d**). Nevertheless, there were notable exceptions, including cCRE pairs with lower or higher than expected effects. cCRE pairs with lower than expected activities had lower chromatin accessibility and H3K27ac signals in K562, compared to other pairs of cCREs (Fisher’s Exact Tests, *p* < 1.7 × 10^−5^, **Supplementary Table 6b**), suggesting that these regions are less likely to be active in their native chromatin context (**Supplementary Note 2.3, Supplementary Figure 7e, f**). cCREs that had a dominant repressive effect on the activity of their partner cCREs were enriched in motifs for transcriptional repressors, such as GFI1B (Fisher’s Exact Test, *p* = 6.5 × 10^−3^, **Supplementary Table 6c**). A small fraction of cCRE pairs (6-15%, depending on stringency of filtering) exhibited higher than expected STARR activity. For example, three neighboring enhancer cCREs in an intron of the *MTNR1A* gene have moderate to low STARR scores when tested separately in K562 cells (0.01, 0.39, and 0.34 for EH38E3620077, EH38E3620078, and EH38E3620079, respectively, **Figure 2f, g**). When tested together, the first two maintained moderate activity (0.38) while the last two had very high activity (1.62, *p* = 0.03, **Figure 2g**). To investigate this cooperativity, we analyzed the cCRE sequences and determined that EH38E3620078 and EH38E3620079 overlapped motifs for more K562 transcription factors compared to EH38E3620077 (49 and 46, respectively, compared to 27, **Figure 2h, Supplementary Table 6d**). Additionally, EH38E3620078 and EH38E3620079 overlap distinct motifs from one another, with EH38E3620078 overlapping AT-rich motifs (STAT, FOX, and BACH families) and EH38E3620079 overlapping GC-rich motifs (KLF and SP families, **Figure 2h**). These findings also hold in native chromatin context as supported by transcription factor ChIP-seq data (**Supplementary Table 6e**). We hypothesize that the higher than expected cooperativity between EH38E3620078 and EH38E3620079 could be attributed to the increased diversity of transcription factor binding. While our power to detect such events is currently limited by the STARR-seq library construction—as there are far fewer double than single fragments—in the future, we can design experiments to include a wider range of fragment lengths to further study combinatorial effects on a larger number of elements.

### Silencers and dual-function elements among REST-Bound cCREs

Silencers are cis-regulatory elements that repress the transcription of target genes^46,47^. Several studies have identified human silencers using chromatin and functional data^48–53^; however, there is little overlap among these collections (**Supplementary Table 7a, b**). We aimed to identify cCREs that act as silencers by integrating ENCODE data. We started with the well-studied repressor transcription factor REST (Restrictive Element-1 Silencing Transcription factor), which is only expressed in non-neuronal cell types and binds to a sequence motif (**Figure 3a**) in neuron-restrictive silencer elements (NRSEs) to repress neuronal genes in non-neuronal cells^54–56^. We defined candidate NRSEs, which we refer to as REST+ cCREs, by selecting all non-promoter cCREs that contained a REST motif and overlapped the summits of at least five REST ChIP-seq peaks (**Supplementary Note 3.1**, **Supplementary Table 7c**). This resulted in 5,850 REST+ cCREs, spanning all cCRE classes, including 2,516 distal enhancer and 789 CA-TF cCREs (1.9-fold depletion and 9.1-fold enrichment, respectively, chi-square test, *p* < 2.2 × 10^−16^, **Figure 3b, c, Supplementary Table 7d**). We then characterized the functional activity of REST+ cCREs using two complementary functional characterization assays—transgenic mouse enhancer assays (**Supplementary Table 7e**) and whole genome STARR-seq—to assess their enhancer and silencer activities, respectively (**Figure 3d**).

**Figure 3.**
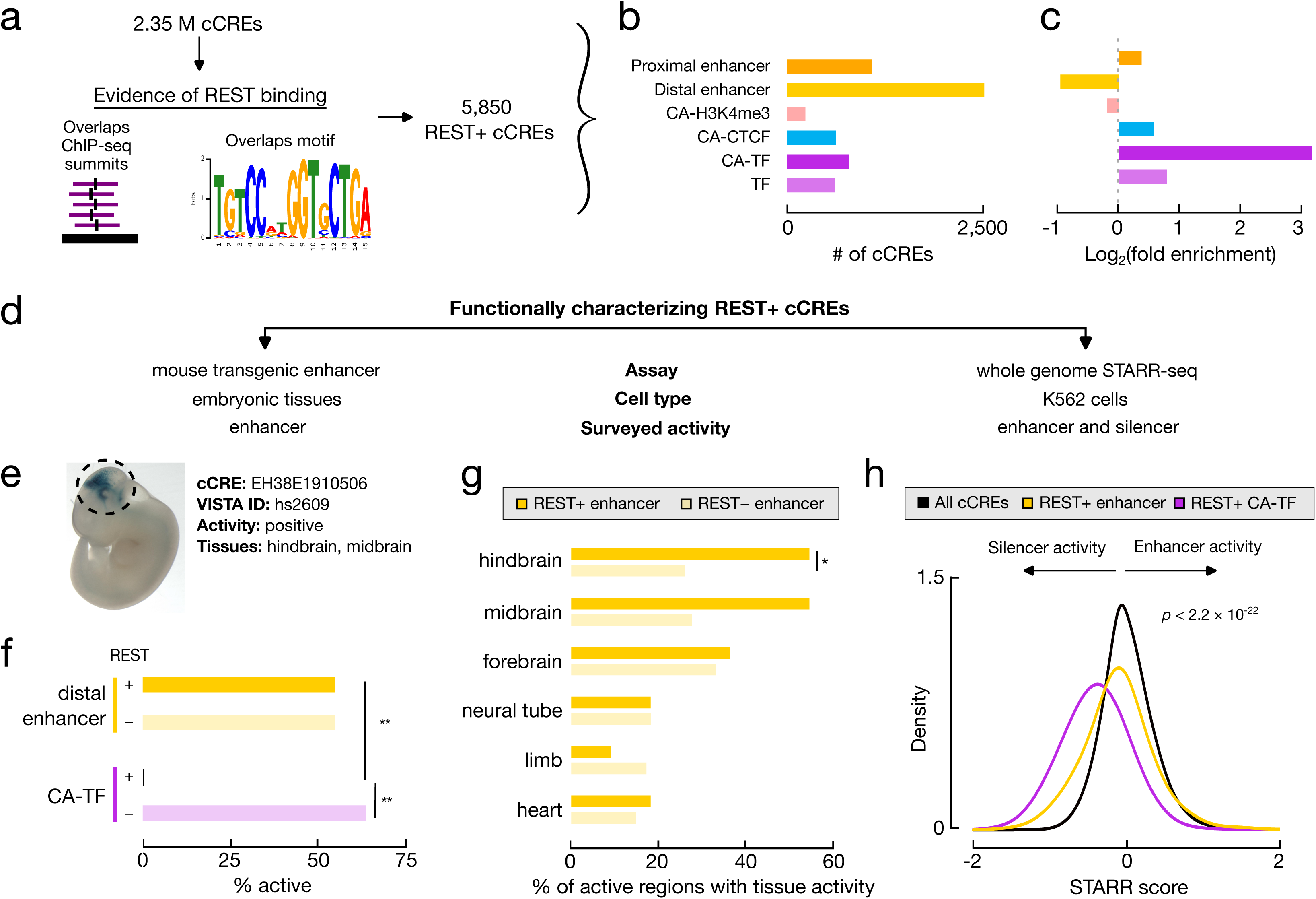
Identification of distinct functional categories of REST-bound cCREs. **a**, Computational pipeline for identifying REST-bound cCREs (REST+ cCREs). We overlapped cCREs with REST ChIP-seq peaks and selected all cCREs that overlap at least five peak summits and an annotated REST motif instance, resulting in 5,850 REST+ cCREs. **b**, Barplots depicting the number of REST+ cCREs stratified by cCRE class. **c**, Barplots depicting the enrichment for cCRE classes of REST+ cCREs compared to the entire Registry. **d**, Workflow for functionally characterizing REST+ cCREs. **e**, Representative result from the mouse transgenic enhancer assay showing activity of REST+ enhancer cCRE EH38E1910506 in mouse brain regions. **f**, Bar graphs denoting the percentage of regions tested in transgenic mouse enhancer assays with positive activity. Regions are stratified into four groups based on cCRE classification (distal enhancer in yellow, CA-TF in purple) and REST binding (+ and dark bars indicate REST+, - and light bars indicate REST-). ** denotes a Fisher Exact Test *p-*value less than 0.01. **g**, Bar graphs showing the percentage of distal enhancer cCREs with transgenic mouse enhancer assay activity in specific tissues stratified by REST binding (as defined in **f**). * denotes a Fisher Exact Test *p-*value less than 0.05. **h**, Density plot of the distributions of STARR scores calculated by CAPRA for all cCREs (black), REST+ distal enhancer cCREs (yellow), and REST+ CA-TF cCREs (purple). Both groups of REST+ cCREs have median STARR scores less than zero, suggesting silencer activity for both groups. *P*-value is calculated using a Wilcoxon test.

Over half of the regions overlapping REST+ distal enhancer cCREs were active in the transgenic mouse enhancer assay—a validation rate as high as regions overlapping distal enhancer cCREs lacking REST binding (REST–, 55% for both sets, **Figure 3e, f, Supplementary Figure 8a**). Compared with the REST– subset, the REST+ subset of distal enhancer cCREs were more likely to be active in brain tissues (2.1-fold enrichment, Fisher’s Exact Test *p* = 0.04, **Figure 3g**, **Supplementary Table 7f**), which was further supported by enrichment for H3K27ac signals in neuron-related biosamples (**Supplementary Note 3.2, Supplementary Table 7g, h**). In K562 cells, these REST+ distal enhancer cCREs had lower than expected STARR scores (median = –0.10, compared with –0.02 for all cCREs, Wilcoxon test, *p* < 2.2 × 10^−16^, **Figure 3h**, **Supplementary Figure 8b**), suggesting they have moderate silencer activities in this cell line. Additionally, genes near REST+ distal enhancer cCREs had lower expression levels than genes near cCREs inactive in K562 cells (median TPM of 0.7 vs 2.4, Wilcoxon test, *p* = 3.0 × 10^−6^, **Supplementary Figure 8c**). We also observed similar activity patterns for a subset of proximal enhancer cCREs—enhancer activity in the brain tissues of transgenic mice and lower than expected STARR scores and gene expression in K562 cells (**Supplementary Figure 8a-c**). Collectively, these results indicate that REST+ enhancer cCREs have dual functions, acting as enhancers when not bound by REST in neurons and acting as silencers when bound by REST in other cell types.

Of the 11 regions overlapping REST+ CA-TF cCREs that were tested in the transgenic mouse enhancer assay, none were active, in sharp contrast with the 64% validation rate for REST– CA-TF cCREs (Fisher’s Exact Test, *p* = 5.5 × 10^−4^, **Figure 3f, Supplementary Figure 8a**). In K562 cells, REST+ CA-TF cCREs had very low STARR scores, even lower than the REST+ distal enhancer cCREs (median = –0.40, Wilcoxon test, *p* < 2.2 × 10^−16^, **Figure 3h, Supplementary Figure 8b**). Genes near REST+ CA-TF cCREs had low expression in K562 cells with a median TPM of 0.1 (Wilcoxon test, *p* < 2.2 × 10^−16^, **Supplementary Figure 8c**). These results suggest that REST+ CA-TF cCREs act as strong silencers in REST-bound cell types such as K562. We observed similar activity patterns for the CA-H3K4me3, CA-CTCF, and TF classes of REST+ cCREs—i.e., low STARR scores and low expression of nearby genes—suggesting that these elements also primarily function as silencers (**Supplementary Figure 8a-c**).

We illustrate the two types of REST+ cCREs using the enhancer/silencer cCRE EH38E2130108 (**Extended Data Figure 4a**) and the silencer cCRE EH38E4127779 (**Extended Data Figure 4b**), both with high REST ChIP-seq signals, moderate DNase signals, and low STARR signals (−0.64 and -0.94, respectively) in K562 cells (**Extended Data Figure 4c,d**). Only EH38E2130108 had enhancer activity, specifically in forebrain, midbrain, and hindbrain mouse tissues of transgenic mice (**Extended Data Figure 4e,f**), which was further supported by high H3K27ac and DNase signals in human *in vitro*-derived neurons. On the other hand, EH38E4127779 lacked enhancer activity in transgenic mice and enhancer signatures in neurons. Neither these example REST+ cCREs nor REST+ cCREs as a whole were enriched for any specific repressive chromatin signatures such as H3K9me3 or H3K27me3 in K562 cells (**Supplementary Table 7i**).

### Annotation of silencer elements using STARR-seq data

Building on our findings with REST+ cCREs, we extended our analysis to more broadly annotate silencer elements using STARR-seq data. This approach enabled us to identify additional silencer cCREs beyond NRSEs that have significant repression activity in K562 cells. We identified 545 stringent (*p* < 0.01) and 5,396 robust (*p* < 0.05) STARR-silencer cCREs in K562, 9% and 5% of which were also REST+ cCREs, respectively (**Extended Data Figure 5a, b, Supplementary Table 8a**). Like REST+ cCREs, STARR-silencer cCREs were enriched for CA-TF, TF, CA-CTCF and CA-H3K4me3 cCREs (**Extended Data Figure 5c**), again highlighting the importance of including these new categories in our expanded cCRE classification scheme.

Similar to genes near REST+ cCREs, genes located near STARR-silencer cCREs had lower expression levels than genes near cCREs with active chromatin signatures in K562 or inactive cCREs (median TPMs of 0.5 and 1.4 for stringent and robust STARR-silencer cCREs respectively, pairwise Wilcoxon test with FDR correction, *p* < 6.6 × 10^−4^, **Supplementary Figure 8c**). STARR-silencer cCREs were also enriched for motifs for transcription factor GFI1B (> 2.6-fold enrichment, *p* < 2.2 × 10^−16^, **Supplementary Table 8b**), the same repressor we identified in anti-cooperative cCRE pairs described above. In native chromatin context, STARR-silencer cCREs were enriched for ChIP-seq peaks of six transcription factors and chromatin remodelers including SETDB1, TRIM28, MIER1, ZNF146, and ZNF280A, in addition to REST (**Supplementary Table 8c**). Distinct groups of STARR-silencer cCREs emerged when clustered by the binding of these six proteins, suggesting sub-classes of silencer elements (**Supplementary Figure 8d**). Despite the enrichment for chromatin remodeling factors, none of the STARR-silencer cCRE groups were enriched for repressive chromatin signatures in K562 cells; they were only depleted for active histone marks (**Supplementary Table 8d**).

To assess STARR-silencer cCRE activity in native chromatin context, we integrated results from K562 CRISPRi-FlowFISH experiments targeting 18 genes across four genomic loci^57^. Of the 734 perturbed cCREs, two were STARR-silencer cCREs (**Supplementary Table 8e**), with EH38E4193243 being one of them. This cCRE is both a STARR-silencer (STARR score of −1.0, *p* = 0.04) and a REST+ enhancer/silencer. It is a proximal enhancer for retbindin (*RTBDN*), a gene preferentially expressed in the retina^58^. It has active enhancer signatures exclusively in WERI-Rb-1, a retinoblastoma cell line, suggesting that this cCRE has highly cell type-specific enhancer activity. In other biosamples (including K562, where CRISPRi-FlowFISH was performed), the cCRE is bound by REST and *RTBDN* is not expressed.

CRISPRi perturbation of EH38E4193243 resulted in increased expression of *PRDX2* (**Supplementary Note 4.1**). Though this cCRE lies 30 kb upstream of *PRDX2*, RNAPII and CTCF ChIA-PET interactions demonstrate that this cCRE is proximal to *PRDX2* in three dimensional space (**Extended Data Figure 5d**). Perturbing the anchors of these interactions also resulted in a similar increase in *PRDX2* expression (**Supplementary Note 4.1**). These results suggest that silencers can impact the expression of multiple genes. We can use our annotated STARR-silencer and REST+ silencer cCREs to aid in future CRISPRi-FlowFISH study design to further investigate such long-range silencer interactions and validate additional silencer cCREs in their native chromatin contexts.

### Novel silencer cCREs have unique chromatin accessibility patterns

Our collection of 9,972 silencer cCREs—comprising both types of REST+ cCREs and STARR-silencer cCREs—only overlap a subset of previously-annotated silencers^48–53^, with the highest concordance with silencers identified by Jayavelu *et al.*, who also used a STARR-based method (**Supplementary Note 4.2, Supplementary Table 7a**). For regions identified by both studies, 93% of silencer cCREs were classified as silencers by Jayavelu *et al*. (2.3-fold enrichment, Fisher’s Exact Test, *p* < 2.2 × 10^−16^, **Extended Data Figure 6a**). However, Jayavelu *et al.* only tested 3% of our silencer cCREs in their assay, suggesting that we have identified many novel silencers (**Extended Data Figure 6b**).

A major distinction of our silencer annotation lies in our cCRE-based approach. Previous annotations typically centered on open chromatin regions within the specific cell type where silencer activity was assayed or in a limited number of biosamples. Only 14% of our silencer cCREs show high chromatin accessibility in K562 cells (**Extended Data Figure 6c, Supplementary Note 4.3**). Instead, silencer cCREs have high chromatin accessibility in early-stage cell types—embryonic stem cells, induced pluripotent stem cells, and *in vitro-*derived progenitor cells—and fetal tissues (**Extended Data Figure 6d-f, Supplementary Note 4.3, Supplementary Table 8f, g**). Therefore, by using chromatin accessibility measurements in a comprehensive collection of biosamples, we are able to identify silencer elements that would otherwise be missed.

### Integrating the Registry of cCREs with other ENCODE Encyclopedia annotations aids in identifying disease-associated genes

The ENCODE Encyclopedia encompasses a comprehensive set of sequence, element, gene, and interaction annotations^20^. The Registry of cCREs, one component of the Encyclopedia, overlaps the vast majority of these other annotations and can serve as an anchor for comparative and integrative analyses (**Supplementary Note 5, Supplementary Tables 9-11**), allowing researchers to study complex loci and better understand transcriptional regulation. To aid in such investigations, we previously developed the web-based search and visualization tool SCREEN (Search candidate cis-Regulatory Elements by ENCODE) for users to query, explore, and visualize cCREs and their underlying data (**Figure 1, Supplementary Figure 9**). During ENCODE4, we extensively re-engineered SCREEN to improve performance and usability (**Supplementary Note 6**).

The Registry of cCREs and SCREEN are particularly useful for aiding the interpretation and prioritization of phenotype-associated genetic variants. Genome-wide association studies (GWAS) have linked over 97,000 genetic variants with hundreds of traits and diseases^59^. Several groups, including ours, have demonstrated that regulatory element annotations can identify phenotype-relevant cell types and pinpoint causal variants in associated loci^1,60–68^. With our updated Registry of cCREs, we can expand the biological scope and accuracy of these annotations, facilitating the identification of novel trait-associated genes and genetic variants.

To demonstrate, we used the Registry of cCREs and SCREEN annotations to annotate and dissect the *RTBDN-MAST1* locus, which was previously associated with red blood cell traits by three independent studies (**Figure 4a, Extended Data Figure 7a**). Variants associated with red blood cell traits are enriched in cCREs active in K562 cells (**Figure 4b**, **Supplementary Table 12a**), which have similar properties and proteomic profiles to early-stage erythrocytes.^69,70^ Therefore, we prioritized K562-specific cCREs and Encyclopedia annotations to identify the causal gene in this region.

**Figure 4.**
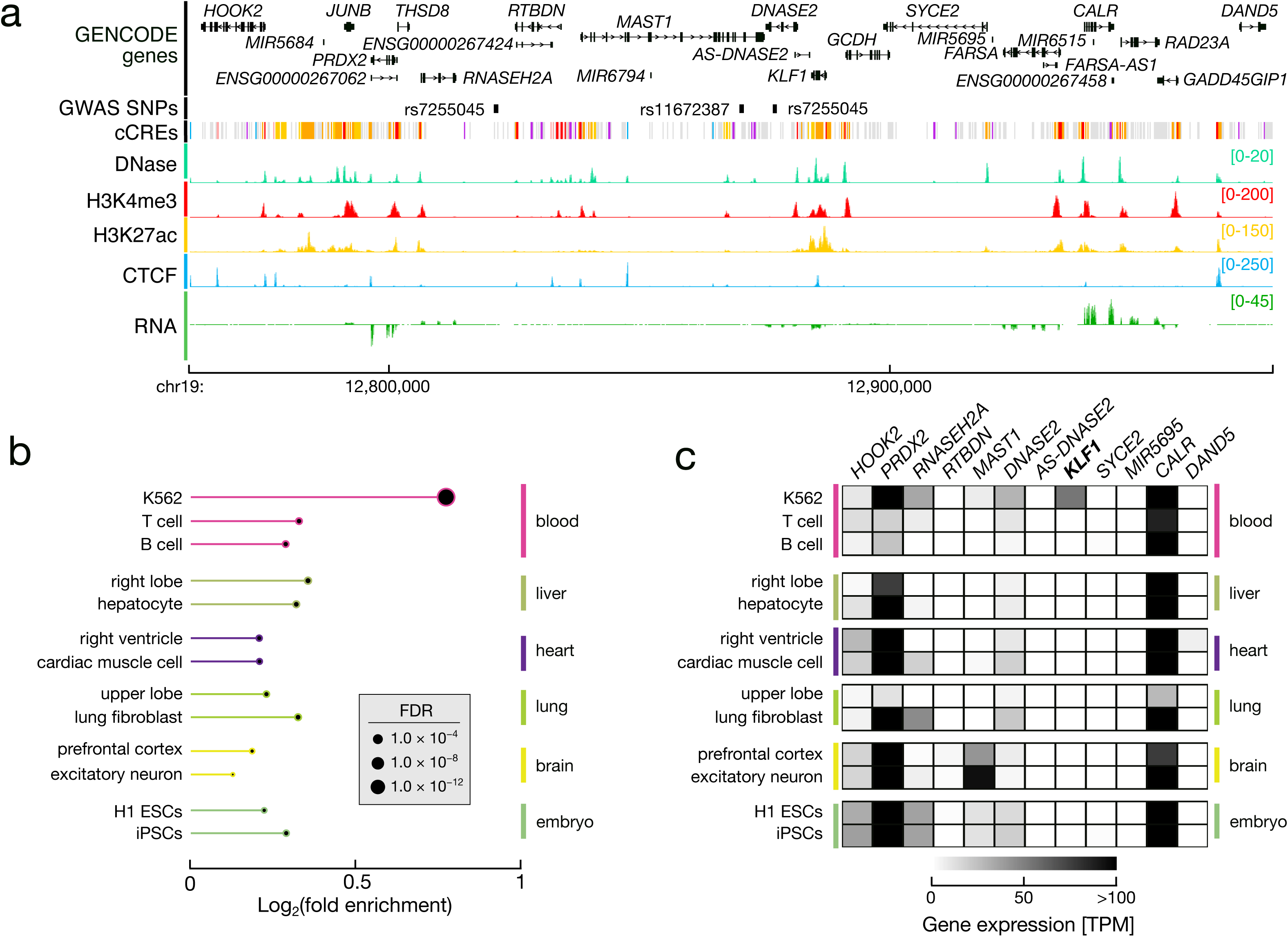
Using the Registry of cCREs to study transcriptional regulation. **a**, Genome browser view of three SNPs associated with red blood cell traits, rs7255045, rs11672387 and rs7255045, and nearby genes. Biochemical signals from K562 cells are shown: DNase in light green, H3K4me3 in red, H3K27ac in yellow, CTCF in blue, and RNA-seq in dark green, along with cCRE classifications in K562. **b**, Circle-barplots denoting the enrichment of variants associated with red blood cell traits in cCREs with high H3K27ac signals in specific cell and tissue types. Length of line denotes the log2 fold enrichment over control variants and size of the terminal circle indicates statistical significance of enrichment. Color of line denotes tissue/organ of origin of biosample. A representative set of biosamples is shown with one cell line and one tissue sample for each non-blood tissue/organ. **c**, Heatmap depicting the expression level of genes linked and/or proximal to variants associated with red blood cell traits across the same biosamples as **b**.

Nine K562 cCREs overlapped SNPs in high linkage disequilibrium (R^2^ > 0.7) with at least one reported red blood cell trait GWAS variant in this locus (referred to as lead variants; **Figure 4a, Extended Data Figure 7b**). To narrow down possible causal genes, we selected all genes that were connected to one of these cCREs in K562 cells using RNAPII and CTCF ChIA-PET links, intact Hi-C loops, and CRISPRi-FlowFISH perturbations. We also included the closest gene by linear distance to each of these nine cCREs along with all the original genes reported by each of the GWAS. In total, this resulted in a list of 12 potential causal genes (**Extended Data Figure 7b, Supplementary Table 12b**).

Of these candidate genes, we predicted *KLF1* to be the likely causal gene based on its known function and expression pattern (**Figure 4c, Extended Data Figure 7c, Supplementary Table 12b**). *KLF1* is a hematopoietic-specific transcription factor that regulates erythroid differentiation by inducing expression of beta-globin and other erythroid-specific genes.^58,71^ Of the 12 candidate genes, *KLF1* is the only gene with K562-specific expression, i.e., high expression in K562 and low expression in all other surveyed cell types (**Figure 4c, Extended Data Figure 7c, Supplementary Table 12c**). We expected a locus that is solely associated with red blood cell traits and no other phenotypes to be associated with a cell type-specific gene. These results support *KLF1* as the gene associated with red blood cell traits, a novel finding for this locus.

Although additional validation is needed to confirm exactly which variant or variants in the locus directly impact *KLF1* expression, we can use the Registry of cCREs to prioritize candidate variants for future testing. For example, rs2290688, which overlaps CA-CTCF cCRE EH38E3291318, is a likely candidate, as it is in high linkage disequilibrium with all three lead GWAS variants (**Extended Data Figure 7b**), and perturbing EH38E3291318 via CRISPRi-FlowFISH resulted in a decrease in *KLF1* expression (**Supplementary Note 7.1**). EH38E3291318 is a major anchor for CTCF-mediated 3D chromatin interactions; it overlaps CTCF ChIA-PET links in 100% of surveyed biosamples and Hi-C loops in 82% of surveyed biosamples, and also has high CTCF signal in 99% of surveyed biosamples (**Supplementary Table 12d-f**). We hypothesize that rs2290688 impacts 3D interactions at this locus, resulting in a rewiring of the regulatory landscape, ultimately impacting *KLF1* expression. Other potential variants for future validation include rs2280742, which overlaps a myeloid enhancer cCRE near *KLF1*, and rs2072597, a benign missense variant in *KLF1* (**Supplementary Note 7.2**, **Supplementary Figure 10, Supplementary Table 12g**). Using the Registry of cCREs, we are able to narrow a list of 63 high linkage disequilibrium variants down to a prioritized set of just a few likely candidates. In summary, our analysis of the *RTBDN-MAST1* locus revealed *KLF1* as a novel causal gene associated with red blood cell traits and demonstrates how the Registry of cCREs, coupled with other ENCODE Encyclopedia annotations, can improve our ability to annotate complex loci and prioritize phenotype-relevant variants.

## Discussion

The expanded Registry of cCREs offers a powerful tool for dissecting the intricate mechanisms of transcriptional regulation across diverse biological contexts. By integrating comprehensive data from functional assays and ENCODE annotations, the Registry enables researchers to study the interplay between cCREs and their impact on gene expression, providing deeper insights into the regulatory logic that governs cellular states and contributes to disease risk.

A key innovation of this updated Registry is its integration with functional characterization datasets, including STARR-seq, MPRA, and CRISPR perturbation assays. These high-throughput methods allow for the direct testing of sequence-driven regulatory potential, facilitating the identification of sequence features that drive regulatory activity. However, there are inherent limitations; for instance, episomal assays like STARR-seq and MPRA may not fully recapitulate the native chromatin context, leading to discrepancies in regulatory activity. Despite this, functional assays remain invaluable for large-scale screening and can guide more detailed, context-specific investigations.

Our findings have several important implications for future research. First, the observed cooperativity between neighboring cCREs suggests that future functional experiments should account for potential interactions between regulatory elements, as these interactions may play a crucial role in gene regulation. Second, elements with dual functions—such as REST+ cCREs, which act as enhancers in neurons and silencers in non-neuronal cells—underscore the need to study regulatory elements across multiple cellular contexts to fully understand their functional roles. Additionally, while cancer cell lines like K562 are commonly used for epigenomic studies, their aberrant gene expression and chromatin landscapes necessitate careful interpretation, especially when generalizing findings to other biological systems.

A significant outcome of our study is the identification and characterization of silencer elements, including those bound by the repressive transcription factor REST and exhibiting repressive activity in STARR-seq assays. Many of these silencers were previously overlooked due to their lower chromatin accessibility in the cell types where they function. While we have identified thousands of novel silencers, our current methods capture only certain subclasses with independent, episomal activities. A more comprehensive understanding will require the development of additional functional assays that can identify silencers operating within complex chromatin environments and context-dependent manners.

The ongoing development of the Registry of cCREs is essential for advancing the field of gene regulation. Future efforts should aim to integrate more comprehensive datasets, particularly those from single-cell assays and underrepresented tissue types, to enhance the resolution of regulatory maps. Additionally, understanding how cCREs interact with each other and with other genomic elements will be crucial for constructing detailed regulatory networks governing gene expression.

In conclusion, our study highlights the utility of the expanded Registry of cCREs as a resource for identifying, characterizing, and understanding complex regulatory loci. With its broad biological scope, detailed annotations, and integration with functional data, the Registry provides a valuable framework for studying CREs across various biological contexts. Continued expansion and refinement of this resource will drive the discovery of novel regulatory mechanisms and enhance our understanding of their implications in health and disease.

## Methods

### The Registry of cCREs

#### Anchoring cCREs

We anchored cCREs on two classes of elements: representative DNase Hypersensitivity Sites (rDHS) and transcription factor clusters. To annotate rDHSs, we refined and filtered DHS calls across biosamples and then iteratively clustered and selected the strongest DHS as previously described^1^. To annotate transcription factor clusters, we downloaded peak BED files for all transcription factor ChIP-seq experiments from the ENCODE portal with a FRIP score > 0.003. For each experiment, we used the “preferred default” peak file and resized peaks smaller than 150 bp and peaks larger than 350 bp to 150 and 350 bp, respectively, to match the cCRE size distribution. Then, using the same interactive clustering and selection process described above, we identified representative transcription factor peaks for each cluster. To complement our rDHS anchors, we selected all transcription factor clusters representing at least five experiments that did not overlap an rDHS. For each anchor, we then calculated the Z-scores of the log-transformed signal for each DNase within a biosample to normalize across datasets. cCREs were defined as regions with high DNase max-Z (>1.64) or overlap a transcription factor cluster.

#### Relevant scripts

0_Call-DHSs.sh

1_Process-DHSs.sh

2_Create-rDHSs.sh

3_Curate-TF-Clusters.sh

4_Calculate-Signal-Zscores.sh

5_Determine-Max-Zscores.sh

#### Classifying cCREs

We classified cCREs into eight classes by analyzing DNase-seq, H3K4me3, H3K27ac, and CTCF ChIP-seq signals and transcription factor binding across biosamples. For each cCRE, we calculated the Z-scores of the log-transformed signal for each mark within a biosample to normalize across datasets. The maximum Z-score (max-Z) for each mark was determined across all biosamples. Based on combinations of high or low signals for these marks, their genomic distance from annotated transcription start sites, and overlap with transcription factor clusters, cCREs were categorized into promoter-like, enhancer-like (TSS-proximal or distal), CA-H3K4me3, CA-CTCF, CA-TF, CA, or TF classes. Classifications were performed both cell type-agnostically and in specific biosamples, following previously described methods.

#### Relevant scripts

6_Classify-cCREs.sh

7_Call-Cell-Type-Specific.sh

#### Gene annotations

Unless otherwise stated, all analyses used GENCODE V40 basic gene annotations for human and GENCODE M25 basic annotations for mouse. ENCODE RNA-seq data was uniformly processed by the ENCODE DCC using GENCODE V29 comprehensive for human and GENCODE M21 comprehensive for mouse.

#### Estimating the number of mouse enhancer cCREs

To estimate the upper bound of the number of possible “missing” mouse enhancer cCREs, we subtracted the number of TSS distal CA cCREs annotated in biosamples with both DNase and H3K27ac data from the total number of annotated distal CA cCREs. This left only CA cCREs in biosamples lacking H3K27ac data which have the potential to be classified as enhancer cCREs if H3K27ac were profiled in those biosamples.

#### Relevant scripts

Figure-1c.Estimate-Mouse-Enhancers.sh

### cCRE sequence features

#### Variational Autoencoder

We analyzed cCREs classified as active in three cell lines—K562, HepG2, and HCT116—focusing on promoters, proximal and distal enhancers, and CA-H3K4me3 elements for each respective cell type. Each cCRE was adjusted to a fixed width of 300 bp, and its sequence was represented using one-hot encoding. A cell type-specific variational autoencoder (VAE) was trained for each cell type, utilizing a ten-dimensional latent space. The encoder architecture included a convolutional layer, max pooling, flattening, and a fully connected dense layer, while the decoder consisted of dense, reshape, upsampling, and convolutional layers. Training was performed for 25 epochs using Adam optimization, implemented in Keras/TensorFlow. After training, the mean values of the latent dimensions for each cCRE were projected into two dimensions using UMAP with 100 nearest neighbors.

#### Relevant scripts

[Jupyter Notebooks](https://github.com/weng-lab/ENCODE-cCREs/tree/master/Version-4/cCRE-Analysis/Supplementary-Figure-4)

#### GC content

For each cCRE we calculated its GC content by counting the number of “G” and “C” nucleotides that appear in its sequence and dividing by the total length of the cCRE

#### Relevant scripts

calculate_CGdinucleotide.py

#### Transcription factor motif overlap

We scanned cCREs for HOCOMOCO v11 motifs^72^ using FIMO^73^ with the default settings and the --text parameter.

### Functional Characterization Data

#### Whole genome STARR-seq

For peak-based analyses, we downloaded BED peak files from the ENCODE portal and intersected them with the Registry of cCREs using BEDTools^74^ (**Supplementary Table 5a**). For enrichment analysis (**Supplementary Figure 5ab**), we intersected peaks (ENCFF454ZKK) from experiment ENCSR661FOW with cCREs active in K562, all other cCREs and non-cCRE size-matched genomic regions (generated using the bedtools random function). For fragment-based analyses, we downloaded BAM files from the ENCODE portal and processed them with the CAPRA pipeline (detailed below).

#### Relevant scripts

Supplementary-Table-5a.STARR-Peak-Summary.sh

Supplementary-Figure-5ace.FCC-Positive-Overlap.sh

Supplementary-Figure-5bdf.FCC-Positive-Overlap.sh

#### Massively parallel reporter assays (MPRA)

For each MPRA experiment we downloaded “element quantification” files from the ENCODE portal. If element coordinates were on the hg19 genome, we lifted coordinates to the GRCh38 genome using the UCSC liftOver tool^75^. We stratified tested regions as to whether they completely overlapped a cCRE (bedtools -f 1 flag), partially overlapped a cCRE, or did not overlap any cCRE (bedtools -v flag). We considered all tested regions with a log2 fold enrichment greater than one as active. We calculated activity enrichment for each group against a baseline activity of all tested regions.

For enrichment analysis (**Supplementary Figure 5c,d**), we intersected tested regions (ENCFF677CJZ) from experiment ENCSR653LWA with cCREs active in K562, all other cCREs and non-cCRE size-matched genomic regions (generated using the bedtools random function).

#### Relevant scripts

[Supplementary-Table-5b.MPRA-Summary.sh](https://github.com/weng-lab/ENCODE-cCREs/blob/master/Version-4/cCRE-Analysis/Supplementary-Table-5a.STARR-Peak-Summary.sh)

Supplementary-Figure-5ace.FCC-Positive-Overlap.sh

Supplementary-Figure-5bdf.FCC-Positive-Overlap.sh

#### CRISPR perturbation screens

For CRISPR perturbation experiments (**Supplementary Table 5c**), we intersected coordinates of CRISPR guide RNAs with cCREs and calculated per-cCRE counts for input libraries and output measurements. We used DESeq2 to calculate cCREs with significant depletion or overrepresentation depending on the exact assay.

#### Relevant scripts

Supplementary-Table-5c.CRISPR-gRNA-Overlap.sh

#### CRISPRi-FlowFISH

For PrimeFlow readout experiments generated by the Engreitz lab^57^, we downloaded element quantification files from the ENCODE portal (**Supplementary Table 5d**). We lifted region coordinates to the GRCh38 genome using UCSC liftOver^75^ and intersected them with cCREs using BEDTools^74^.

For HCR-FlowFISH readout experiments generated by the Sabeti lab, we downloaded CASA elements^27^ and intersected them with cCREs using BEDTools. We used these elements for enrichment analysis (**Supplementary Figure 5ef**), where CASA elements were intersected with with cCREs active in K562, all other cCREs and non-cCRE size-matched genomic regions (generated using the BEDTools random function).

#### Relevant scripts

Supplementary-Table-5d.CRISPRi-FlowFISH-Overlap.sh

Supplementary-Figure-5ace.FCC-Positive-Overlap.sh

Supplementary-Figure-5bdf.FCC-Positive-Overlap.sh

### CAPRA quantifications

#### Pipeline

Using BEDtools^74^, the first step of the pipeline converts BAM files into fragment BED files for both the input DNA and output RNA fragments. Next, for each fragment BED file, fragments are intersected with cCREs. Fragments that overlap a single cCRE in its entirety are counted in the “solo” quantifications. Fragments that overlap two cCREs in their entirety are counted in the “double” quantifications. Fragments that partially overlap one or more cCREs are excluded. Quantifications for input DNA and output RNA fragments (including multiple replicates when available) are concatenated into a matrix for each quantification type—e.g. solo and double. Quantification matrices are processed by DESeq2 to identify cCRE with higher than expected RNA counts (denoting enhancer activity) or lower than expected RNA counts (denoting silencer activity).

#### GitHub Repo

https://github.com/Moore-Lab-UMass/CAPRA

#### Cell type-specific activity

Using the solo fragment STARR scores (normalized log_2_ fold enrichment of RNA over DNA) calculated by CAPRA we defined K562+ and HepG2 + STARR distal enhancers as follows:

- K562+ STARR distal enhancers: K562 STARR score > 0 <u>AND</u> K562 STARR score > HepG2 STARR score +1
- HepG2+ STARR distal enhancers: HepG2 STARR score > 0 <u>AND</u> HepG2 STARR score > K562 STARR score +1

#### Relevant scripts

Figure-2d.STARR-Cross-Cell-dELS.sh

#### Transcription factor enrichment

Using the HOCOMOCO motif sites identified by FIMO (see above) we calculated the number of K562+ and HepG2+ STARR distal enhancers that overlap a motif for each transcription factor. We considered all transcription factors with greater than five transcripts per million (TPM) in either K562 or HepG2 (N = 234). We compared motif overlap between the two cell types using Fisher’s Exact test with FDR correction (**Supplementary Table 6a**) and visualized the top five most enriched transcription factor motifs in each cell type that appear in at least 10% of the respective K562+ or HepG2+ STARR distal enhancers (Figure 2e).

Similar analysis was performed to look for motif enrichment in negative combinatorial pairs of cCREs. We selected all cCRE pairs meeting the following requirements (N = 1,275):

- One cCRE of the pair had independent STARR activity with a positive solo STARR score and p < 0.05
- The other cCRE of the pair had a negative solo STARR score
- The double STARR score of the pair was negative

We compared the overlap of motifs between cCREs with the positive and negative solo STARR scores, considering all transcription factors expressed in K562 (TPM > 5, N = 181) and using Fisher’s Exact test with FDR correction (**Supplementary Table 6c**).

#### Relevant scripts

Figure-2e.CTS-STARR-TF-Motif.sh

Supplementary-Table-6c.Repressive-Dual-cCRE-Motif-Enrichment.sh

### Silencer cCREs

#### Defining REST+ cCREs

Using BEDTools^74^, we intersected cCREs with the summits of ENCODE REST ChIP-seq peaks (**Supplementary Table 7c**) and selected all cCREs that overlapped the summits of at least five peaks. We then selected all cCREs that overlapped at least one REST motif site from FactorBook^76^ (**Supplementary Table 7c**). We then filtered out promoter cCREs (**Supplementary Note 3.1**) resulting in a total of 5,850 REST+ cCREs.

#### Relevant scripts

Figure-3a.Curate-REST-cCREs.sh

#### Testing REST+ regions with transgenic mouse enhancer assays

To supplement previously tested regions in the VISTA database, we performed transgenic mouse enhancer assay experiments for 26 REST+ regions (**Supplementary Table 7e**). These regions were conserved between mouse and human and had evidence of REST binding (i.e., ChIP-seq peaks) in both species. Nine of these regions overlapped REST+ enhancer/silencer cCREs, 15 overlapped REST+ silencer cCREs and 2 overlapped cCREs that overlapped REST ChIP-seq peaks but were not included in our REST+ silencer sets (**Supplementary Table 7e**).

The transgenic mouse assays were conducted using the FVB/NCrl strain of Mus musculus (Charles River) as previously described^77^. Candidate regions were amplified via PCR and inserted into a plasmid containing a minimal Hsp68 promoter and a lacZ reporter gene. The resulting constructs were injected into fertilized mouse eggs, which were then implanted into surrogate mothers. Embryos were harvested at embryonic day 11.5 (E11.5) and analyzed for β-galactosidase activity. Regions were classified as active enhancers if reproducible staining was observed in the same tissue in at least three embryos. Regions were considered inactive if no consistent staining was detected, provided that at least five embryos with transgene insertions were analyzed.

#### Overlap of REST+ cCREs with VISTA enhancers

We downloaded the coordinates of 1,947 tested regions from the VISTA Enhancer Database (available as of April 22, 2023), which included the aforementioned 26 regions, and lifted the coordinates of these regions to the GRCh38 genome using UCSC liftOver^75^. We then intersected VISTA regions with REST+ cCREs using BEDTools intersect, calculating the fraction of tested regions with positive activity. All analysis was performed relative to the number of unique VISTA regions and we used Fisher’s exact test to calculate statistical significance. For distal enhancer cCREs, we calculated enrichment of activity in specific tissues using Fisher’s exact test by comparing the activity of regions overlapping REST+ distal enhancer cCREs versus REST- distal enhancer cCREs.

#### Relevant scripts

Figure-3g.Supplementary-Table-7e.REST-Silencer-VISTA-Enrichment.sh

#### Defining STARR-silencer cCREs

Using STARR scores calculated by CAPRA from ENCSR661FOW, we annotated all cCREs with a negative STARR score and *p* < 0.01 as stringent STARR-silencers and all cCREs with a negative STARR score and *p* < 0.05 as robust STARR-silencers (**Supplementary Table 8a**).

#### Enrichment in chromatin signatures

For each biosample with DNase data (N = 1,325, **Supplementary Table 8f**) we calculated the fraction of silencer cCREs with high signal (z-score > 1.64, **Extended Data Figure 6d**). To determine enrichment, we calculated the log2 fold enrichment of the fraction of silencer cCREs over the fraction of all cCREs with high DNase signal. For a subset of silencer cCREs, REST+ silencer cCREs, we compare the enrichment of chromatin accessibility between fetal and adult biosamples from the same tissue or organ (e.g. muscle, lung, kidney, and brain, **Supplementary Table 8g**). Statistical significance was calculated using a Wilcoxon test.

#### Relevant scripts

Supplementary-Table-8f.Silencer-DNase-Enrichment.sh

We also intersected silencer cCREs with histone mark peaks from K562 cells (**Supplementary Table 7h,8d**) using BEDTools intersect. We downloaded histone mark peaks directly from the ENCODE portal. For REST+ cCREs, we compared peak overlap with cCRE active and inactive in K562, respectively. For stringent and robust STARR-silencers we compared peak overlap with cCREs with positive STARR scores and neutral STARR scores. We calculated p-values using Fisher’s Exact test with FDR correction.

#### Relevant scripts

Supplementary-Table-7h.REST-cCRE-Histone-Enrichment.sh

[Supplementary-Table-8d.STARR-Silencer-Histone-Enrichment.sh](https://github.com/weng-lab/ENCODE-cCREs/blob/master/Version-4/cCRE-Analysis/Supplementary-Table-7h.REST-cCRE-Histone-Enrichment.sh)

#### Overlap with previous silencer collections

We downloaded K562 silencers from four previous studies and formatted them into BED files. This included elements from Supplemental Table 2 from Huan *et al*.^48^, Supplementary Data 2 from Jayavelu *et al.*^49^, Supplementary Table 1 from Pang *et al.*^50^, and Supplementary Data 1 from Cai *et al.*^52^. When necessary, region coordinates were lifted to the GRCh38 genome using UCSC liftOver^75^. We intersected each collection with the Registry of cCREs using BEDTools^74^. We also compared silencer calls between each study by calculating the overlap coefficient.

#### Relevant scripts

Supplementary-Table-7a.Silencer-Comparison-cCRE.sh

Supplementary-Table-7b.Silencer-Comparison.sh

#### Expression of nearby genes

Using BEDTools^74^ we identified the closest GENCODE annotated protein coding gene for each cCRE as measured by linear distance to the nearest TSS. Using gene quantifications in K562 (ENCFF421TJX, available on the ENCODE portal), we extracted transcripts per million (TPM) quantifications for each gene and stratified by cCRE class/group.

#### Relevant scripts

Supplementary-Figure-8c.Silencer-Gene-Expression.sh

### Dissecting GWAS loci

#### VESPA pipeline

As previously described^1^, the VESPA pipeline takes lead variants reported by GWAS and generates a set of matched controls based on variant minor allele frequency and genomic context. For both sets of GWAS and control variants, VESPA then retrieves variants in high linkage disequilibrium (LD, R^2^>0.7) based on study population, creating “LD blocks”. VESPA then intersects variants with cCREs. For studies with a sufficient number of LD blocks (at least 25 lead variants), we can calculate cell type/tissue enrichment by subsetting cCREs based on activity in specific biosamples and comparing overlap between GWAS and control variants, accounting for LD by only counting each LD block once per intersection. For this study, we used H3K27ac signals across 562 biosamples. In total, we curated variants from 3,751 unique trait-study combinations, 396 of which also have biosample recommendations, the results of which are available on SCREEN.

#### GitHub Repo

https://github.com/Moore-Lab-UMass/VESPA

#### Candidate GWAS genes

We identified candidate GWAS genes using three complementary approaches:

- *Closest gene*: Using BEDTools closest^74^, we identified the closest gene (either protein coding or non-coding) to each candidate cCRE as measured by linear distance to the nearest transcription start site.
- *3D chromatin interactions*: We intersected cCREs with the anchors of 3D chromatin loops—RNAPII ChIA-PET, CTCF ChIA-PET and Hi-C—from K562 cells. To link a candidate cCRE to a gene, we required one end of the loop to overlap the cCRE and the other end of the loop to fall within 2kb of an annotated transcription start site.
- *CRISPRi-FlowFISH*: We intersected the GRCh38 mapped coordinates with candidate cCREs and selected all cCRE-gene pairs with an FDR < 0.05.

For the cross-biosample, cross-gene expression analysis (Figure 4c) gene quantification files were downloaded directly from the ENCODE portal (experiment and file IDs in **Supplementary Table 12b**). For *KLF1* gene expression data was downloaded directly from SCREEN (**Extended Data Figure 7c, Supplementary Table 12c**).

## Data Availability

All cCRE annotations (cell type-agnostic and cell type-specific) are available on SCREEN (screen.wenglab.org). All other ENCODE data are available on the ENCODE portal (encodeproject.org) and are referenced by experiment and file accession. Supplementary Data are also available at https://users.moore-lab.org/ENCODE-cCREs/Supplementary-Data/.

All data used and generated in this study adhere to ENCODE guidelines and comply with institutional standards, including approval by the appropriate Institutional Review Board (IRB) for human-related data and Institutional Animal Care and Use Committee (IACUC) for animal research.

## Code Availability

All code is available on the following repositories:

https://github.com/weng-lab/ENCODE-cCREs/tree/master/Version-4
https://github.com/Moore-Lab-UMass/CAPRA
https://github.com/Moore-Lab-UMass/VESPA
https://github.com/weng-lab/SCREEN2.0

Specific scripts are also referenced in the corresponding Methods sections.

## Acknowledgements

This study was funded by National Institutes of Health grants U24HG009446 and U24HG012343 (Z.W.), UM1HG009390 (B.E.B.), UM1HG009382 (M.L), UM1HG009442 (M.S.), UM1HG009443 (A.M.), UM1HG009421 (L.A.P), UM1HG009428 (T. R.), and U01HG009380 (M.B.) The research of D.E.D, A.V, and L.A.P. was conducted at the E.O. Lawrence Berkeley National Laboratory and performed under U.S. Department of Energy Contract DE-AC02-05CH11231, University of California.

We are grateful to the ENCODE Consortium for its collective efforts in enabling large-scale data production, advancing methods development, and facilitating data analysis, which provided the foundation for this work. We thank the ENCODE Registry of cCRE Working Group (**Supplementary Note 8.1)** for providing valuable feedback on this work and also acknowledge the ENCODE TSS Annotation Working Group (**Supplementary Note 8.2)** for their contributions to improving transcription start site annotations. We also extend our thanks to the ENCODE Biosamples Working Group (**Supplementary Note 8.3)** for their critical role in coordinating sample generation, which provided essential resources for this study. Finally, we would like to thank Mansi Khandpekar and Andres Colubri for their help in redesigning the SCREEN logo.

## Ethics declarations (competing interests)

J.M.E. is an inventor on patents and patent applications related to CRISPR screening technologies, has received materials from 10× Genomics unrelated to this study, and has received speaking honoraria from GSK plc. B.E.B. discloses financial interests in HiFiBio, Arsenal Biosciences, Chroma Medicine, Cell Signaling Technologies and Design Pharmaceuticals. M.P.S. is a co-founder and on the advisory boards of Personalis, Qbio, January AI, SensOmics, Filtricine, Protos, Mirvie, Onza, Marble Therapeutics, Iollo, and NextThought AI. He is also on the advisory boards of Jupiter, Applied Cognition, Neuvivo, Mitrix, and Enovone. A.K. is a consulting fellow with Illumina; a member of the SABs of OpenTargets (GSK), PatchBio, and SerImmune; and a co-founder of RavelBio. Z. W. is a cofounder of Rgenta Therapeutics and serves on its scientific advisory board. The other authors declare no competing interests.

## Author contributions

J.E.M. and Z.W. conceived and designed the study. A.K., A.M., M.S, T.R., and M.L. provided regular feedback on computational methodology, biological interpretation, and manuscript preparation. J.E.M. H.E.P., and K.F., performed the data analysis with contributions from S.I.E, G.A., M.G., N.S., and Y.F. Additional supporting analysis was performed by A.P., V.R., F.R., B.B., M.Y., E.W., M.R-R., M.R-M., J.X., T.G. under the supervision of J.E.M., Z.W., M.A.B., R.G., M.B.G., A.M., and A.K., H.E.P., N.P., and J.F, developed the SCREEN web portal with contributions from M.C.L., J.M., M.G., E.C., and R.S. N.F, C.B.E, E.G. and B.B. led the generation of histone modification data and coordinated cross-consortium data generation efforts. D.E.D, A.V, and L.A.P. carried out transgenic mouse enhancer assays and contributed related biological insights. T.R. and J.E. provided functional characterization data and contributed related biological insights. J.E.M. and Z.W. wrote the manuscript with input from all authors. All authors reviewed and approved the final version of the manuscript.

