## Extended Data Figures for "An Expanded Registry of Candidate cis-Regulatory Elements for Studying Transcriptional Regulation"

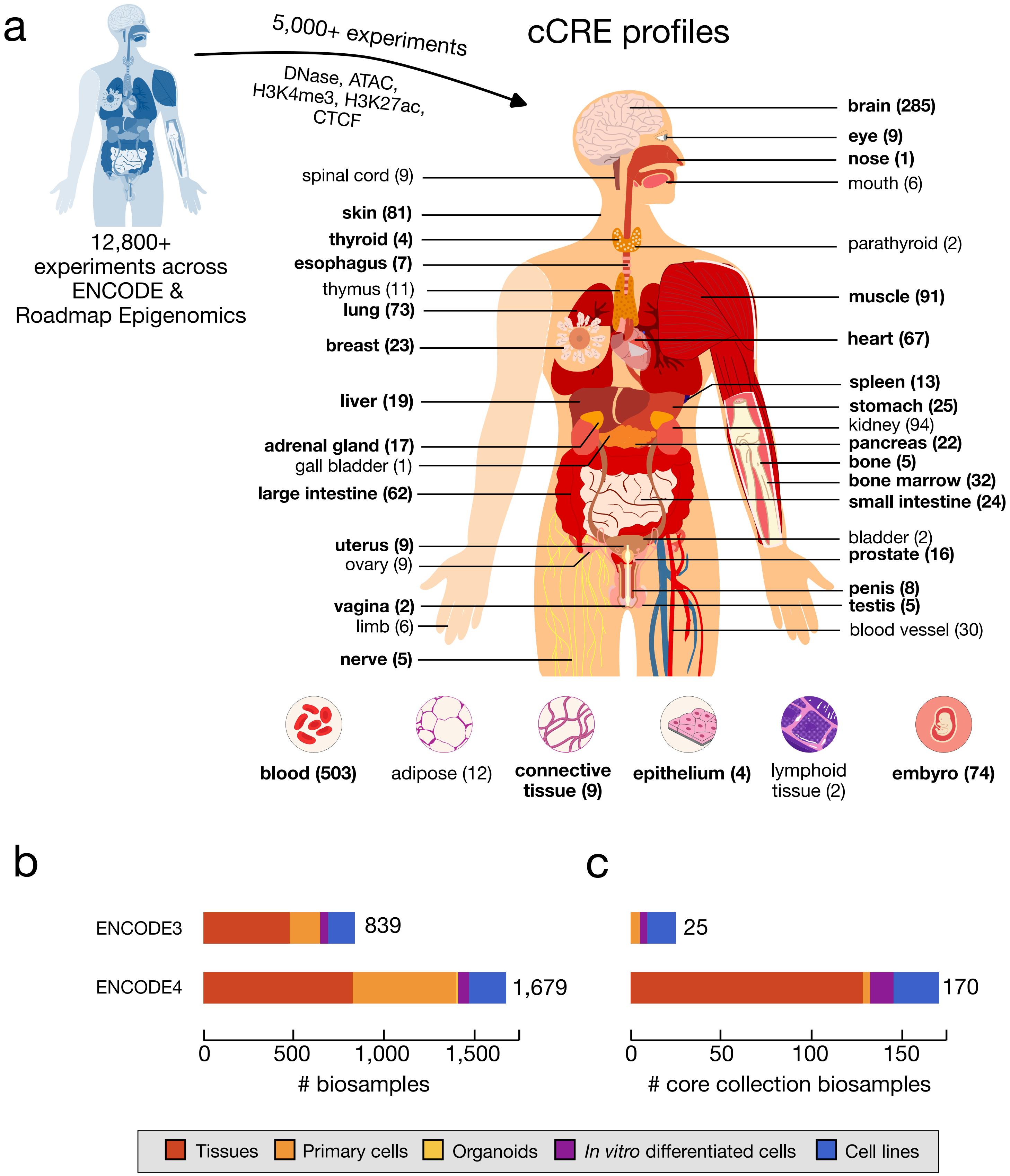
a
12,800+ experiments across ENCODE & Roadmap Epigenomics
5,000+ experiments
DNase, ATAC, H3K4me3, H3K27ac, CTCF
cCRE profiles
brain (285)
eye (9)
nose (1)
mouth (6)
spinal cord (9)
skin (81)
thyroid (4)
parathyroid (2)
esophagus (7)
thymus (11)
lung (73)
muscle (91)
breast (23)
heart (67)
spleen (13)
liver (19)
stomach (25)
kidney (94)
adrenal gland (17)
pancreas (22)
gall bladder (1)
bone (5)
bone marrow (32)
large intestine (62)
small intestine (24)
uterus (9)
bladder (2)
ovary (9)
prostate (16)
penis (8)
vagina (2)
testis (5)
limb (6)
blood vessel (30)
nerve (5)
blood (503)
adipose (12)
connective tissue (9)
epithelium (4)
lymphoid tissue (2)
embyro (74)
b
ENCODE3
839
ENCODE4
1,679
0
500
1,000
1,500
### biosamples
c
25
170
0
50
100
150
### core collection biosamples
Tissues
Primary cells
Organoids
In vitro differentiated cells
Cell lines

#### Extended Data Figure 1 | Increased biosample coverage of the Registry of cCREs

**a,** Body map denoting the number of biosamples in the Registry of cCREs deriving from each major tissue or organ. Tissues/organs in bold denote that biosamples in this category are part of the core collection (having DNase, H3K4me3, H3K27ac, and CTCF), and the numbers indicate the number of biosamples belonging to each category. **b,** Barplots showing the number of biosamples in ENCODE3 vs ENCODE4 versions of the Registry of cCREs, stratified by biosample type: tissues in red, primary cells in orange, organoids in yellow, *in vitro*-differentiated cells in purple, and cell lines in blue. **c,** Barplots showing the number of biosamples in each version of the Registry of cCREs that have all four core marks (DNase, H3K4me3, H3K27ac, and CTCF), stratified by biosample type and colored as defined in **b**.

Moore *et al.* Extended Data Figure 2

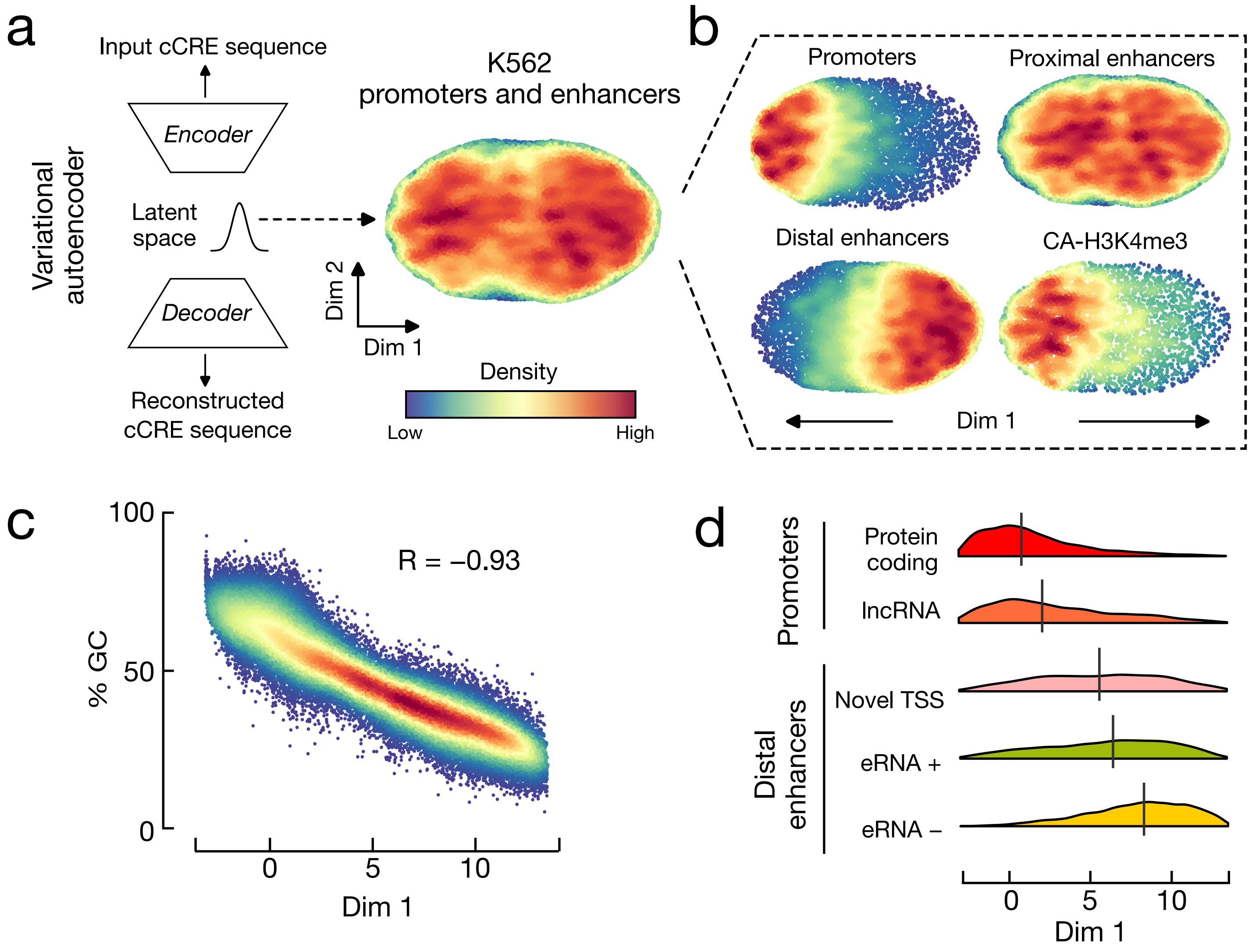

**Extended Data Figure 2 | Biochemical signal-centric cCRE classification is concordant sequence-based clustering**

**a,** Schematic of variational autoencoder used to extract sequence features of promoter and enhancer cCREs. After the model is trained, the latent space is sampled for each cCRE and then projected onto two-dimensions. **b**, Density scatterplots of a two-dimensional UMAP projection of the trained variational autoencoder's latent space for each cCRE, stratified by cCRE class. Red corresponds to high density and dark blue corresponds to low density. **c,** Density scatterplot depicting the correlation between the first visualized dimension in **a** and **b** and the percent GC content of input cCRE sequence (PCC, $R = -0.93$). **d**, Density plots showing the distribution of dimension 1 values (sequence scores) for promoters (top) stratified by gene class (protein coding: red, lncRNA: orange) and distal enhancers (bottom) stratified by overlap with experimentally-derived transcription start sites (long-read RNA: pink, PRO-cap: green, neither: yellow). All distributions are significantly different (Pairwise Wilcox test with FDR correction, $FDR < 1.0 \times 10^{-6}$).

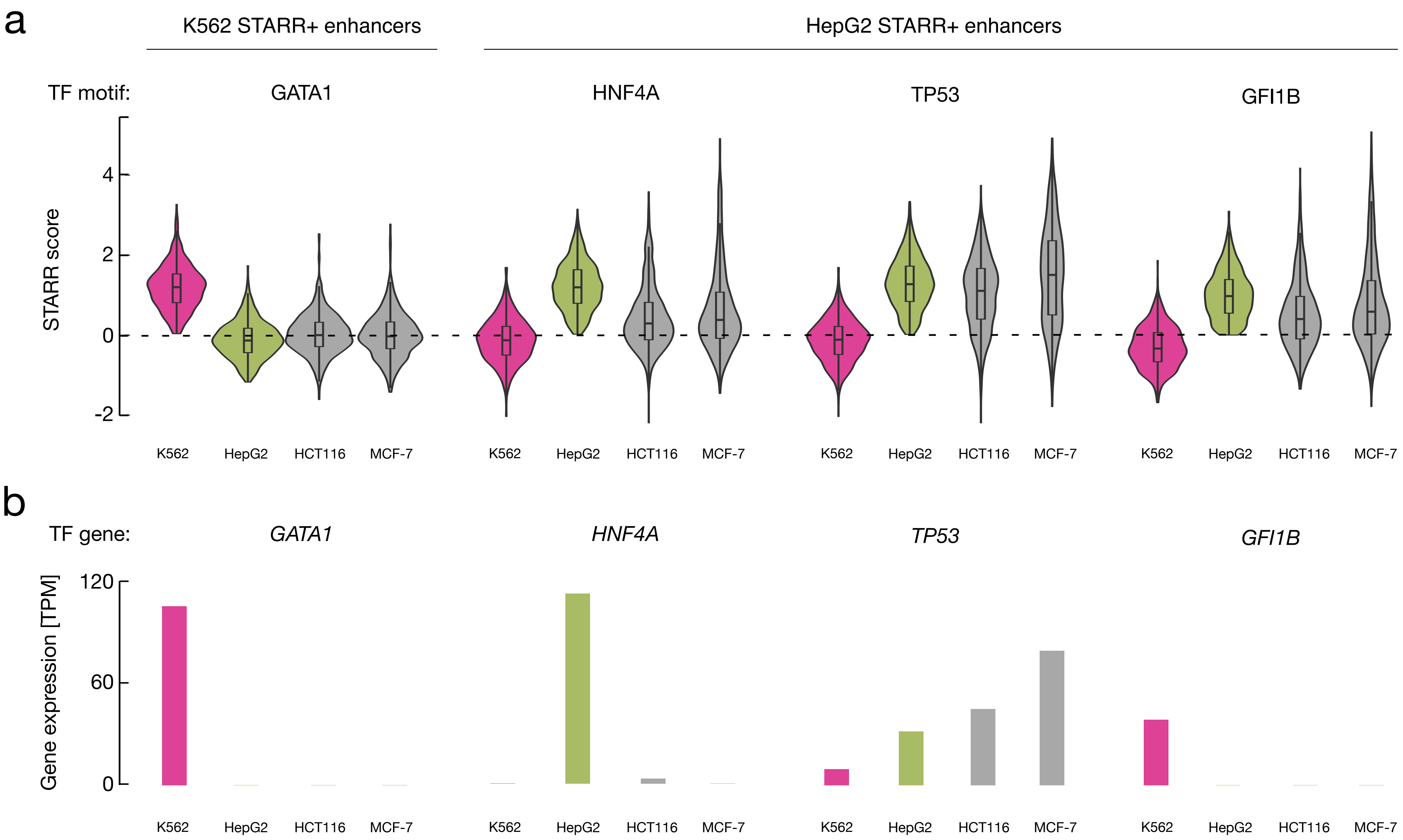
a
K562 STARR+ enhancers
HepG2 STARR+ enhancers
TF motif:
GATA1
HNF4A
TP53
GFI1B
STARR score
4
2
0
-2
K562
HepG2
HCT116
MCF-7
b
TF gene:
GATA1
HNF4A
TP53
GFI1B
Gene expression [TPM]
120
60
0
K562
HepG2
HCT116
MCF-7

**Extended Data Figure 3 | STARR activity of cCREs is concordant with cell type-specific molecular profiles**

**a,** Nested violin boxplots showing STARR scores for each of K562 STARR+ distal enhancer cCREs that overlap GATA1 motifs (left) and HepG2 STARR+ distal enhancer cCREs that overlap HNF4A, TP53, and GFI1B motifs, respectively (right). STARR scores are from four whole genome STARR-seq experiments including K562 (pink), HepG2 (green), HCT116 (gray), and MCF-7 (gray). **b**, Barplots depicting gene expression in transcripts per million (TPM) of *GATA1*, *HNF4A*, *TP53* and *GFI1B*, respectively, in K562 (pink), HepG2 (green), HCT116 (gray), and MCF-7 (gray) cell lines.

Moore *et al*. Extended Data Figure 4

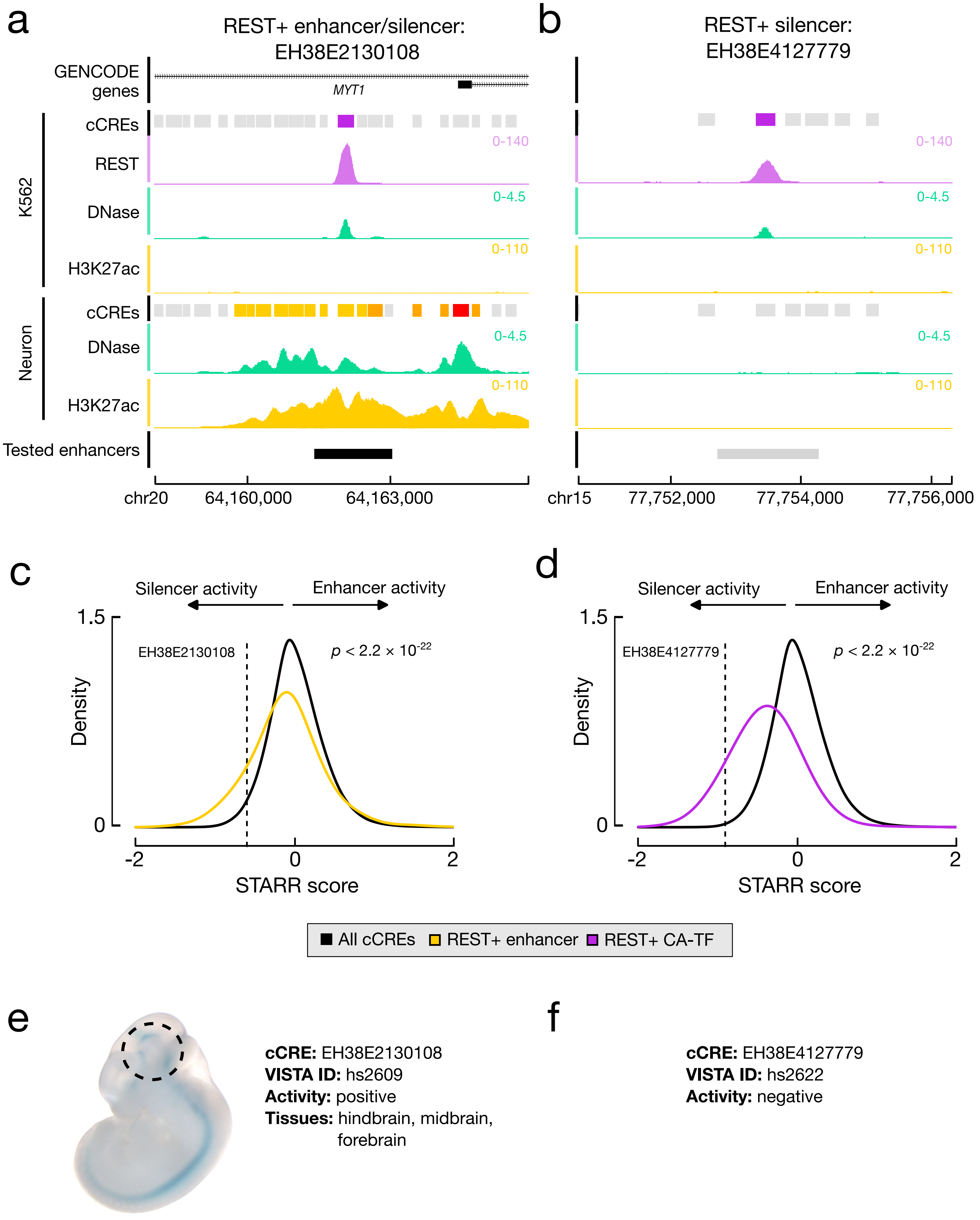

**Extended Data Figure 4 | Example cases of REST+ cCRE classes**

**a,** Genome browser view of REST+ enhancer/silencer EH38E2130108. EH38E2130108 has high REST ChIP-seq signal (purple), moderate DNase signal (green), and low H3K27ac signal (yellow) in K562, which results in it being classified as CA-TF in K562. In *in vitro*-differentiated neurons, EH38E2130108 is classified as an enhancer because it has high DNase and H3K27ac signals. The black bar denotes the genomic region tested in the transgenic mouse enhancer assay. **b**, Genome browser view of REST+ CA-TF EH38E4127779 with biochemical signals as defined in **a**. EH38E4127779 has high REST ChIP-seq signal, moderate DNase signal, and low H3K27ac signal in K562, which results in it being classified as CA-TF in K562. In *in vitro*-differentiated neurons, EH38E4127779 has no enhancer signatures and tested negative in the transgenic mouse enhancer assay. **c,** Density plot of the distributions of STARR scores calculated by CAPRA for all cCREs (black) and REST+ distal enhancer cCREs (yellow). Dashed line indicates STARR score for EH38E2130108. **d,** Density plot of the distributions of STARR scores calculated by CAPRA for all cCREs (black) and REST+ CA-TF cCREs (purple). Dashed line indicates STARR score for EH38E4127779. **e**, Representative results from transgenic mouse enhancer assay for EH38E2130108. Dashed circle indicates activity in hindbrain, midbrain, and forebrain. **f**, No positive results from testing EH38E4127779 in the transgenic mouse enhancer assay.

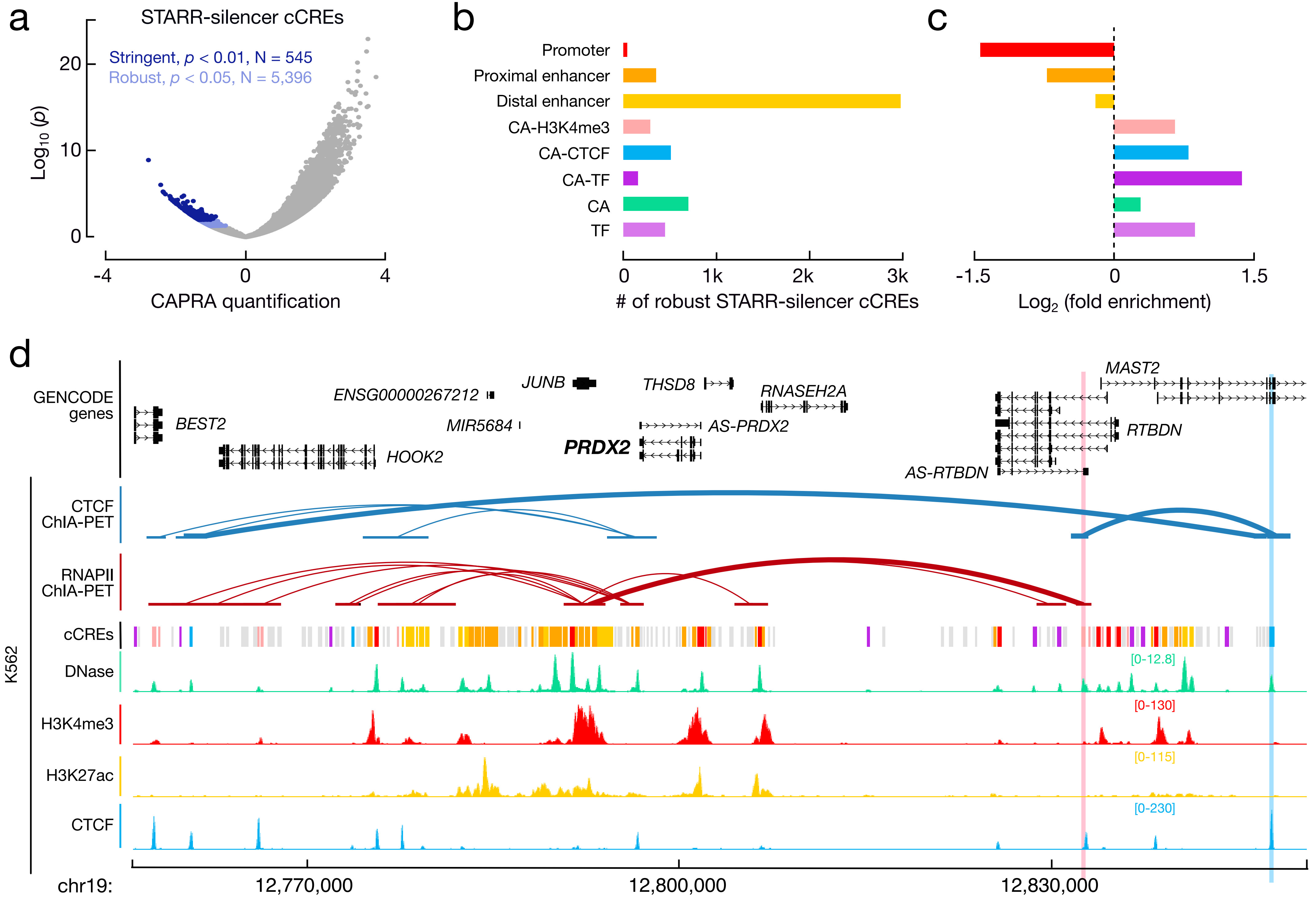
a
STARR-silencer cCREs
Stringent, p < 0.01, N = 545
Robust, p < 0.05, N = 5,396
Log10 (p)
0
10
20
-4
0
4
CAPRA quantification
b
Promoter
Proximal enhancer
Distal enhancer
CA-H3K4me3
CA-CTCF
CA-TF
CA
TF
0
1k
2k
3k
### of robust STARR-silencer cCREs
c
-1.5
0
1.5
Log2 (fold enrichment)
d
GENCODE genes
BEST2
HOOK2
ENSG00000267212
MIR5684
JUNB
PRDX2
THSD8
AS-PRDX2
RNASEH2A
MAST2
RTBDN
AS-RTBDN
K562
CTCF ChIA-PET
RNAPII ChIA-PET
cCREs
DNase
H3K4me3
H3K27ac
CTCF
[0-12.8]
[0-130]
[0-115]
[0-230]
chr19:
12,770,000
12,800,000
12,830,000

#### Extended Data Figure 5 | Identification and characterization of silencer cCREs

**a,** Volcano plot depicting effect size versus significance of CAPRA quantifications for all cCREs. Stringent STARR-silencer cCREs ($p < 0.01$, N = 545) are in dark blue and robust STARR-silencer cCREs ($p < 0.05$, N = 5,396) are in light blue. **b,** Barplots depicting the number of robust STARR-silencer cCREs stratified by cCREs class. **c,** Barplots depicting the enrichment for cCRE classes of robust STARR-silencer cCRE compared to the entire Registry. **d**, Genome browser view showing physical 3D genome interactions between silencer EH38E4193243 (pink highlight) and CTCF-anchor EH38E3291318 (blue highlight) and a domain of actively transcribed genes. Targeting either silencer EH38E4193243 or CTCF-anchor EH38E3291318 results in increased expression of *PRDX2* (bold). CTCF ChIA-PET links (dark blue) directly connect silencer EH38E4193243 and CTCF-bound EH38E3291318. CTCF-anchor EH38E3291318 is also connected with upstream cCREs, forming a domain which brings silencer EH38E4193243 near the active promoters, as evidenced by RNAPII ChIA-PET links (dark red). Biochemical signals (DNase: green, H3K4me3: red, H3K27ac: yellow, CTCF: blue) and cCRE classifications for K562 cells are displayed.

Moore *et al.* Extended Data Figure 6

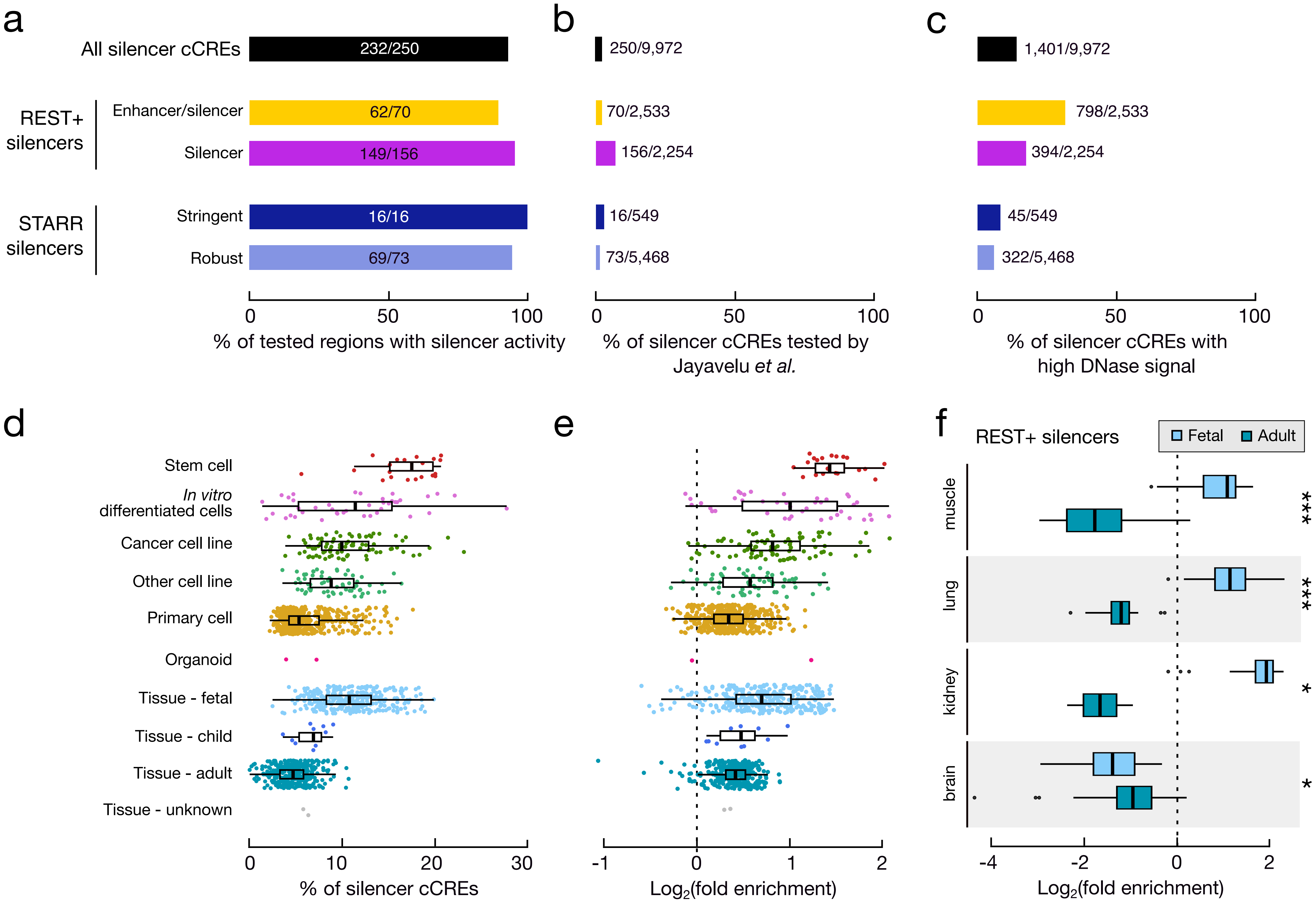

**Extended Data Figure 6 | Concordance and characteristics of silencer cCREs**

**a,** Barplots depicting the percentage of silencer cCREs tested by Jayavelu *et al.* that were deemed to have silencer activity. Metrics for all silencer cCREs are in black, REST+ enhancer/silencer cCREs in yellow, REST+ silencer cCREs in purple, stringent STARR-silencer cCREs in dark blue, and robust STARR-silencer cCREs in light blue. **b**, Percentage of silencer cCREs tested by Jayavelu *et al.* stratified as described in **a**. **c,** Percentage of silencer cCREs with high DNase signal in K562 cells, stratified as described in **a**. **d,** Boxplots with scatterplot overlays depicting the percentage of silencer cCREs with high chromatin accessibility in each biosample, grouped by biosample type. **e,** Boxplots with scatterplot overlays depicting the log2 fold enrichment for high chromatin accessibility at silencer cCREs compared to all cCREs, grouped by biosample type. **f,** Boxplots depicting log2 fold enrichment for high chromatin accessibility at REST+ silencer cCREs compared to all cCREs for matched fetal and adult tissues. Differences between matched adult and fetal tissues were significantly different (Paired Wilcox test with FDR correction, * indicates FDR $< 0.05$, *** indicates FDR $< 0.001$).

Moore *et al.* Extended Data Figure 7

**a**

| Lead SNP | Author | PMID | *p* | Trait |
|---|---|---|---|---|
| rs2418568 | Astle *et al.* | 27863252 | $6 \times 10^{-9}$ | Immature fraction of reticulocytes |
| rs7255045 | Ganesh *et al.* | 19862010 | $2 \times 10^{-12}$ | Mean corpuscular volume |
| | van Rooij *et al.* | 28017375 | $5 \times 10^{-33}$ | Mean corpuscular volume |
| rs11672387 | Astle *et al.* | 27863252 | $4 \times 10^{-16}$ | High light scatter reticulocyte count |
| | Astle *et al.* | 27863252 | $1 \times 10^{-21}$ | High light scatter reticulocyte percentage of red cells |
| | Pilling *et al.* | 28957414 | $5 \times 10^{-10}$ | Red cell distribution width |

**b**

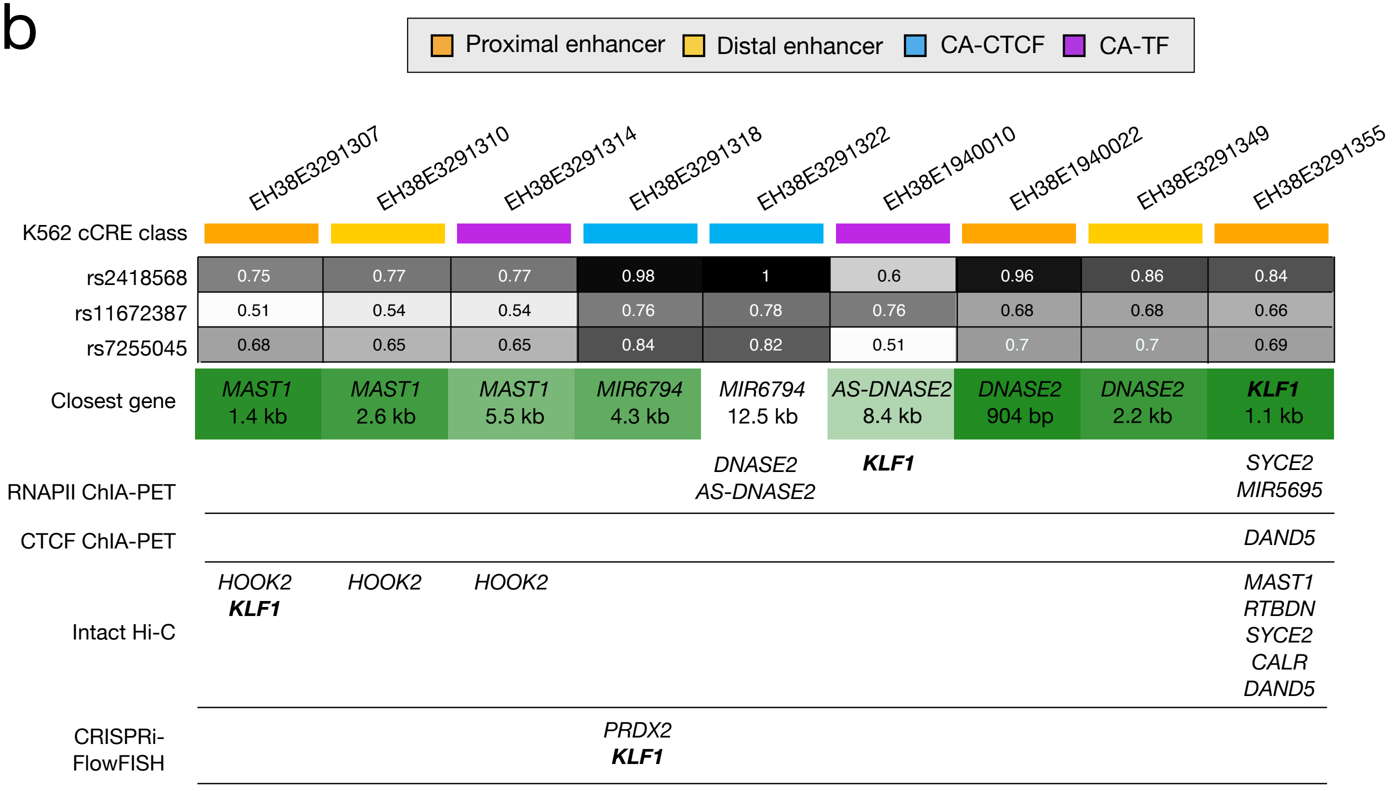


**c**

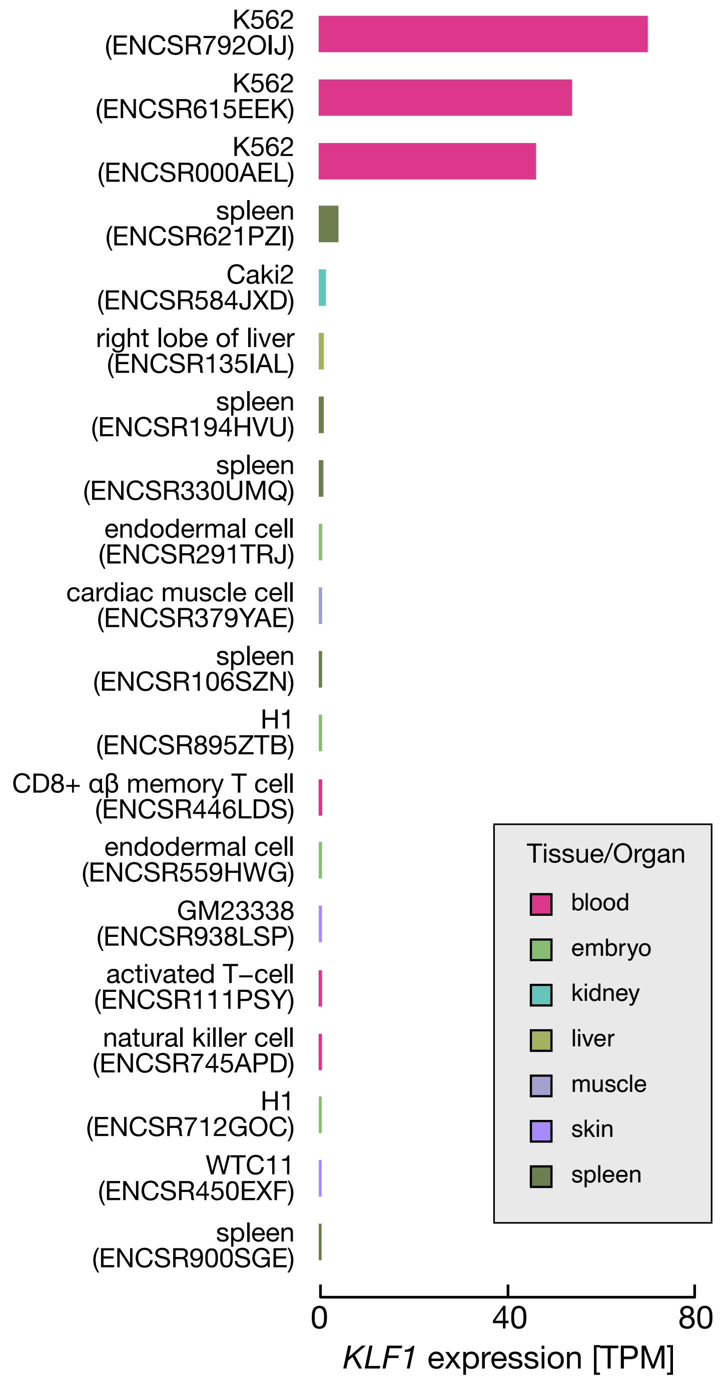

**Extended Data Figure 7 | Using the Registry of cCREs to identify causal GWAS genes**

**a,** Summary of lead SNPs in the *RTBDN-MAST1* locus identified by GWAS. **b**, Candidate genes for cCREs overlapping SNPs in high linkage disequilibrium ($R^2 > 0.7$, identified by VESPA pipeline) with each of the three lead SNPs reported in **a**. Color denotes the cCREs classification in K562 (proximal enhancer: orange, distal enhancer: yellow, CA-CTCF: blue, CA-TF: purple). Gray boxes indicate the linkage disequilibrium $R^2$ with each of the lead GWAS SNPs. Closest gene by linear distance to the nearest TSS is shown in green boxes with color denoting distance. Additional candidate gene targets identified by 3D chromatin interactions (ChIA-PET and intact Hi-C loops) or CRISPRi-FlowFISH are also displayed. **c**, Barplots depicting *KLF1* expression across the top 20 biosamples with highest expression in transcripts per million (TPM). Bars are colored by tissue/organ of origin as defined on SCREEN.
