## Supplementary Information for "An Expanded Registry of Candidate cis-Regulatory Elements for Studying Transcriptional Regulation"

#### Supplementary Tables

Supplementary Table 1 | Summary of datasets in Version 4 of the Registry of cCREs
Supplementary Table 2 | Overlap of CRE-associated biochemical annotations and rDHSs
Supplementary Table 3 | Multi-mapping cCREs
Supplementary Table 4 | Enrichment of Version 4-specific cCREs
Supplementary Table 5 | Functional characterization of cCREs
Supplementary Table 6 | Characterizing cell type and element-specific STARR-seq activity
Supplementary Table 7 | Characteristics of REST+ cCREs
Supplementary Table 8 | Characteristics of STARR-silencer cCREs
Supplementary Table 9 | ENCODE Encyclopedia Element Annotations
Supplementary Table 10 | ENCODE Encyclopedia Transcription Annotations
Supplementary Table 11 | ENCODE Encyclopedia Interactions
Supplementary Table 12 | Identifying genes associated with red blood cell traits

#### Supplementary Data

Supplementary Data 1 | ENCODE4 GRCh38 cCREs
Supplementary Data 2 | ENCODE4 mm10 cCREs
Supplementary Data 3 | ENCODE4 GRCh38 Multi-mapping cCREs
Supplementary Data 4 | ENCODE4 mm10 Multi-mapping cCREs
Supplementary Data 5 | CAPRA Quantifications (Solo)
Supplementary Data 6 | CAPRA Quantifications (Double)

### Supplementary Notes

#### Supplementary Note 1 - Detailed updates to the Registry of cCREs

##### 1.1 Defining element anchors

Similar to previous versions[1], the ENCODE4 Registry of the cCREs is primarily anchored on representative DNase hypersensitivity sites (rDHSs). For ENCODE4, the 2,863,480 rDHSs in humans result from the integration of 178,746,635 individual DHS across 1,438 DNase profiles and the 1,338,731 rDHSs in humans result from the integration of 54,542,655 individual DHS across 466 DNase profiles.

In general, rDHSs capture the vast majority of biochemically active sites across the genome; they had high overlap with experimentally derived TSSs from CAGE and RAMPAGE (overlaps of 90 and 95%, respectively, **Supplementary Table 2a**), active histone mark peaks including H3K4me3 (average overlap of 96%, **Supplementary Table 2b**), H3K27ac (92%,

**Supplementary Table 2c**), and H3K4me1 (91%, **Supplementary Table 2d**) and transcription factor ChIP-seq peaks (average overlap of 99%, **Supplementary Table 2d**). However, there was a subset of transcription factors whose peaks had lower overlap with rDHSs (**Supplementary Figure 1a, Supplementary Table 1**). These included ZNF274 (on average 71% of peaks overlap with rDHSs), MAFK (73%), MAFF (73%), ATF2 (73%), ATF7 (84%), and CREB1 (85%). We analyzed overlap between these non-rDHS transcription factor peaks and defined five primary clusters (**Supplementary Figure 1b**). While there are a relatively few of these regions across the genome, they have higher conservation than non-cCRE regions suggesting they may have a functional role (**Supplementary Figure 1c**). Therefore, to capture these putative regulatory regions in our Registry, we supplemented our rDHSs with representative transcription factor peaks (transcription factor rPeaks)—peaks that represent binding sites for at least three TFs (**Supplementary Figure 1d**).

For the human genome, we curated 821,672 transcription factor rPeaks by combining 2,509 transcription factor ChIP-seq datasets; 86,748 (11%) did not overlap an rDHS and were included in our anchoring scheme. For the mouse genome, we curated 171,092 transcription factor rPeaks by combining 167 transcription factor ChIP-seq datasets; 7,658 (4%) did not overlap an rDHS and were included in our anchoring scheme.

##### 1.2 Multi-mapping sites

Most ENCODE uniform processing pipelines, including those for DNase-seq and ChIP-seq, filter out reads that map to multiple genomic loci known as multi-mapping reads. This filtering was implemented to reduce false positive peak calls and mapping artifacts. However, focusing exclusively on uniquely mapping reads creates "holes" in the Registry, particularly at recently duplicated genes.

To build a more comprehensive collection of cCREs, we incorporated multi-mapping reads when calling DHSs at a curated set of genomic loci. We identified 1,799 loci in the human genome and 1,470 loci in the mouse genome by analyzing high-confidence ATAC-seq peaks that did not overlap rDHSs. Unlike DNase-seq, the ATAC-seq pipeline allows reads to map to the genome up to four times, selecting the best matching location. When reads map equally well to multiple locations, they are randomly assigned. As a result, ATAC-seq can detect peaks in highly similar genomic regions, such as recently duplicated genes, missed by traditional DHS calling (which considers only uniquely mapping reads).

To address these gaps, we called DHSs using multi-mapping reads specifically at the curated loci. This approach added an average of 2,292 DHSs per experiment, resulting in a total of 24,160 additional rDHSs in the human genome (**Supplementary Data 3**) and 40,537 in the mouse genome (**Supplementary Data 4**). These DHSs, which do not overlap existing cCRE anchors, are referred to as *multi-mapping* cCREs.

In humans, the majority of the multi-mapping cCREs were near recently duplicated genes. Acharya *et al*.[78] annotated 11,710 human genes as either arising from whole genome duplication events that took place early during vertebrate evolution (N=7,070, 60%) or more recent small scale duplication events (N=4,640, 40%). Using our standard collection of non-multi-mapping cCREs, we identified promoter cCREs for 96% of whole genome duplication genes but only 69% of small scale duplication genes (**Supplementary Figure 2a,b**, Fisher's exact test, $p < 2.2 \times 10^{-16}$). By including *multi-mapping* cCREs, we now annotate promoters for 667 additional GENCODE V40 genes, of which 96 are small scale duplication genes and 12 are whole genome duplication genes (**Supplementary Figure 2a,b**).

For example, *EIF3C* and *EIF3CL*, encode subunits of the eukaryotic translation initiation factor 3 (eIF3) complex and arose from a small scale duplicate event (**Supplementary Figure 2c,d**). Due to their high sequence similarities, the regions around their promoters have low mappability, as measured by Umap mappability software[79]. By including peaks called from multi-mapping reads, we can now annotate dozens of cCREs in the region, including the promoter cCREs for both genes, that were previously missed. This example highlights how including multi-mapping reads in our analysis provides a more complete annotation of cCREs, particularly at loci with low mappability, offering valuable insights into regulatory elements at recently duplicated genes that would otherwise remain obscured.

#### 1.3 cCRE Data Collections

In both SCREEN's visualization, downloadable files, and UCSC genome browser track hubs, we annotate biosamples based on available data.

##### *Core collection*

Thanks to the extensive coordination efforts by the ENCODE4 Biosample Working Group (**Supplementary Note 7**), 170 biosamples were profiled for DNase, H3K4me3, H3K27ac, and CTCF. We refer to these samples as the biosample-specific *Core Collection* of cCREs. These samples derive from a variety of tissues and organs and mainly comprise primary tissues and cells (**Extended Data Figure 1a,c**). We suggest that users prioritize these samples for their analysis as they contain all the relevant marks for the most complete annotation of cCREs.

##### *Partial Data Collection*

To supplement the Core Collection, 1,154 biosamples have DNase in addition to various combinations of the other marks (but not all three). Though we are unable to annotate the full spectrum of cCRE classes in these biosamples, having DNase enables us to annotate element boundaries with high resolution. Therefore, we refer to this group as the *Partial Data Collection*. In these biosamples, we classify elements using the available marks. For example, if a sample lacks H3K27ac and CTCF, its cCREs can only be assigned to the promoter, CA-H3K4me3, and CA groups, not the enhancer nor CA-CTCF groups. The *Partial Data Collection* contains unique tissues and cell states that are not represented in the *Core Collection*, such as fetal brain tissue and stimulated immune cells that may be of high interest to some researchers. Therefore, if users are interested in cCRE annotations in such biosamples, we suggest leveraging the cell type-agnostic annotations or annotations from similar biosamples in the Core Collection, to anchor their analyses.

##### *Ancillary Collection*

For the 354 biosamples lacking DNase data, we do not have the resolution to identify specific elements and we refer to these annotations as the *Ancillary Collection*. In these biosamples, we simply label cCREs as having a high or low signal for every available assay. We highly suggest that users do not use annotations from the *Ancillary Collection* unless they are anchoring their analysis on cCREs from the *Core* or *Partial Data Collections*.

***Custom Classification***

While our classification schemes place each cCRE into specific, individual classes, the signal strengths for all recorded epigenetic features are retained for each cCRE in the Registry and can be used for customized searches by users. For example, users may want promoters that have high DNase, H3K4me3, and H3K27ac to distinguish from poised promoters that often lack H3K27ac signal. Additionally, by default, all chromatin accessibility annotations use DNase signal. If users prefer to use ATAC signal, this can also be accomplished using the SCREEN API.

#### 1.4 Classification of cCREs

Many uses of cCREs are based on the regulatory role associated with their biochemical signatures. Analogous to GENCODE's catalog of genes, which are defined irrespective of their varying expression levels and alternative transcripts across different cell types, we provide a general, cell type-agnostic classification of cCREs. This classification is based on each element's dominant biochemical signals across all available biosamples and its proximity to the nearest GENCODE transcription start site (**Figure 1b**).

In addition to the cell type-agnostic classification (described in the main text), we also evaluated the biochemical activity of each cCRE in individual biosamples using the corresponding biosample-specific DNase, ATAC, H3K4me3, H3K27ac, and CTCF data. This allows us to annotate active cCREs in individual biosamples. Elements with low DNase Z-scores in individual biosamples are deemed to be inactive and are labeled with "Low Chromatin Accessibility."

Because of the uneven distribution of TF data across biosamples and our desire to reduce false positive annotations, we do not specifically annotate the TF class of elements in individual cell types. We instead provide an aggregate list of these TF cCREs annotated with their supporting

TF ChIP-seq peaks. For cell types with TF ChIP-seq data, we annotate CA-TF elements since the accessible chromatin corroborates the TF binding.

#### 1.5 Expanded Registry of cCREs maintains specificity for functional elements

Our updated Registry saw the addition of 1,419,467 new cCREs. This increase can be attributed to both updates in our computational pipeline—which accounts for 524,533 new cCREs—and an increase in biosample coverage—which accounts for 894,934 new cCREs (**Supplementary Figure 3a**). In previous versions of the Registry, we performed an additional filtering step as detailed in The ENCODE Project Consortium *et al.* Supplementary Note 1[1]. However, such stringent filtering resulted in underrepresentation of elements from cell types and tissues that were missing histone mark data, such as fetal tissue samples profiled during the Roadmap Epigenomics project (**Supplementary Table 4a**). Because these "filtered" cCREs were enriched for STARR-seq activity (**Supplementary Figure 3b,** 4.3 fold enrichment, Fisher's Exact test, $p < 2.2 \times 10^{-16}$) and were evolutionarily conserved[80] (average of 0.14 vs -0.03 at center, Wilcoxon rank sum test, $p < 2.2 \times 10^{-16}$ , **Supplementary Figure 3c**) compared to non-cCRE regions, we removed this additional filtering step in our updated pipeline. The 894,934 cCREs added to the Registry due to an increase in biosample coverage were also more likely to have STARR-seq activity (**Supplementary Figure 3b,** 5.4 fold enrichment, Fisher's Exact test, $p < 2.2 \times 10^{-16}$) and had higher evolutionary conservation (average of 0.04 at center, Wilcoxon rank sum test, $p < 2.2 \times 10^{-16}$ , **Supplementary Figure 3c**) than non-cCRE regions, suggesting that although our Registry has more than doubled its size, it still retains specificity for functional genomic elements.

To further evaluate the comprehensiveness and specificity of the ENCODE4 Registry of cCREs, we compared our collection with cis-regulatory annotations from two recent publications. Li *et al.* performed single nucleus ATAC-seq on over one million cells from 42 brain regions. In total, they identified 107 distinct cell types and subsequently 544,729 candidate cis-regulatory elements active in the brain[32]. In their original report, they found only about half of their elements (51%) overlapped ENCODE3 cCREs. With the expanded Registry, 85% (N=462,009) of their elements overlap ENCODE4 cCREs (**Supplementary Table 4b**) and this increase in coverage was primarily due to newly annotated cCREs active in brain tissues (**Supplementary**

**Figure 3d**, **Supplementary Table 4c**). On average, 46 thousand of the Li *et al.* brain candidate cis-regulatory elements were active in an individual cell type and ENCODE4 cCREs overlap 96% of brain candidate cis-regulatory elements active in one of the 107 cell types (**Supplementary Table 4b**). These results suggest that our collection covers the majority of regulatory regions active in a diverse collection of brain cell types.

Loupe *et al.* performed ChIP-seq for more than 100 transcription factors across nine brain regions, surveying both bulk tissue and nuclei sorted for neuronal markers[33]. They previously reported that about 70% of their collection of 237,324 thousand transcription factor peaks (termed union peaks) overlapped ENCODE cCREs (**Supplementary Table 4d**). We compared their list of union peaks with our updated collection of cCREs and 223,548 of their elements (94%) overlap ENCODE4 cCREs (**Supplementary Table 4d**). Again, this increase in coverage was primarily due to our expansion in brain-specific cCREs (**Supplementary Figure 3e, Supplementary Table 4e**). On average, 85,000 union peaks were bound by transcription factors in an individual brain region or cell type. ENCODE4 cCREs overlap 98% of these experiment-specific elements suggesting once again that our collection covers the majority of regulatory regions active in a diverse collection of brain cell types and regions (**Supplementary Table 4d**).

##### 1.6 Using variational autoencoders to extract cCRE sequence features

We trained three cell type-specific Variational Autoencoders (VAEs) on the sequences of promoter, enhancer, and CA-H3K4me3 cCREs active in K562, HepG2, and HCT116 cells (**Extended Data Figure 2a**). The latent representations learned by the VAEs were then reduced to two dimensions using UMAP for visualization. In all three cell types, promoter and distal enhancer cCREs separated along the first dimension (**Extended Data Figure 2b, Supplementary Figure 4**). Proximal enhancers were evenly distributed across dimension one in K562 and centrally distributed for HepG2 and HCT116. CA-H3K4me3 cCREs clustered near promoters in K562 and HCT116 but were positioned more centrally in HepG2. Differences in clustering for proximal enhancer and CA-H3K4me3 cCREs may reflect biological variations between cell types or could result from differences in data quality.

GC content, defined as the percentage of guanine (G) and cytosine (C) nucleotides in a cCRE, was strongly correlated with the first dimension across all three cell types with Pearson correlation coefficients of -0.93 for K562, -0.91 for HepG2, and -0.88 for HCT116, respectively ($p < 2.2 \times 10^{-16}$). Other sequence features had lower correlation coefficients such as CpG density (-0.78, -0.75, and -0.74, respectively) and normalized CG[81] (-0.61, -0.63 and -0.59, respectively). This indicates that GC content is a key sequence feature captured by our VAE models (**Extended Data Figure 2c**).

We then investigated how the regulatory potential of cCREs differs across subclasses of promoter and enhancer cCREs (**Extended Data Figure 2d**). We stratified promoter cCREs based on whether they were promoters for protein coding or lncRNA transcripts, as defined by GENCODE. Promoter cCREs of protein coding genes (N=17,080) had lower values for dimension one compared to promoter cCREs of lncRNA genes (N=4,175, median of 0.8 vs. 2.0, Pairwise Wilcoxon rank sum test with FDR correction, FDR $< 2.2 \times 10^{-16}$). We also stratified distal enhancer cCREs into three distinct groups based on their overlap with transcription sites: (1) cCREs that overlapped the 5' ends of long RNA-seq reads, presumably the transcription start sites of novel, stable transcripts (N=1,895), (2) cCREs that overlap PRO-cap peaks but not the 5' end of long RNA-seq reads (N=3,916), and (3) all other distal enhancers (N=29,677). Enhancer cCREs that overlapped novel TSSs had the lowest dimension one values (5.5, FDR $< 1 \times 10^{-6}$ ) followed by enhancer cCREs with eRNAs (6.4, FDR $< 2.2 \times 10^{-16}$) and all other distal enhancer cCREs (8.3).

##### 1.7 Mouse enhancers

Like previous versions of the Registry of cCREs, our expanded ENCODE4 Registry of cCREs has far more human data than mouse data. In total, the current Registry integrates 3,202 experiments from 1,679 human biosamples compared to only 591 experiments across 367 mouse biosamples. There was no significant difference in the number of DHSs called from an individual human or mouse DNase-seq experiment with medians of 98,569 and 105,538, respectively (Wilcoxon rank sum test, $p$=0.54), suggesting that similar numbers of cCREs would be annotated across the human and mouse genomes if experiment and biosample coverage were the same.

Additionally, the majority of mouse data generated during ENCODE4 was primarily chromatin accessibility and transcription assays; histone mark ChIP-seq was only generated for four biosamples (hippocampus, cerebral cortex, heart and gastrocnemius). Therefore, the mouse registry has a higher proportion of CA cCREs—cCREs with high levels of chromatin accessibility that lack histone mark signals—compared to the human registry. We hypothesize that many of these CA cCREs would be classified as enhancer cCREs if we had H3K27ac ChIP-seq data in the same tissues. Therefore, to estimate an upper bound on the number of CA cCREs that could be enhancers, we counted all annotated CA cCREs that exclusively derive from biosamples lacking histone mark data, which resulted in 188,912 CA cCREs, 65% of the total annotated mouse CA cCREs (**Figure 1c**).

#### Supplementary Note 2 - Functional characterization analysis

##### 2.1 Integrating functional characterization data with the Registry of cCREs

###### ***Whole genome STARR-seq***

STARR-seq (Self-Transcribing Active Regulatory Region sequencing) is a high-throughput reporter assay that quantifies enhancer activity of DNA sequences by placing them in the 3' untranslated region (UTR) of a reporter gene downstream of a minimal promoter[82]. Active enhancers are part of the resulting transcript and the relative abundance of their transcripts is a measure of enhancer activity. For whole genome STARR-seq, the input library of tested regions derives from fragmented genomic DNA. Candidate enhancers are traditionally identified through peak calling, looking for regions with enrichment for RNA fragments over input DNA fragments.

We integrated the Registry of cCREs with peak calls from seven whole genome STARR-seq experiments generated during ENCODE4 (**Supplementary Table 5a**). Between 69-91% of STARR peaks overlap cCREs with variation in overlap attributed to peak calling method and STARR-seq protocol. Because the median peak width of STARR peaks is greater than the width of cCREs, many STARR peaks overlap multiple cCREs. Therefore to quantify the STARR activity of individual cCREs and annotate both positive and negative activity patterns, we developed a novel computational method to annotate cCRE-specific STARR activity.

***MPRA***

Massively Parallel Reporter Assays (MPRAs) are high-throughput techniques that measure the regulatory activity of thousands of DNA sequences simultaneously by linking each sequence to a reporter gene and quantifying its expression in cells[83]. We integrated the Registry of cCREs with results from 62 ENCODE4 MPRA experiments spanning six cell types (**Supplementary Table 5b**). For each experiment, we compared the activity of tested MPRA regions that (1) fell entirely within a cCRE, (2) partially overlapped a cCRE or (3) did not overlap any cCRE. Tested regions that fell entirely within a cCRE had the highest positivity rates (an average of 23% of tested regions had a $Log_2FC > 1$) compared to those that partially overlapped cCREs (14%) and those that did not overlap cCREs (7%, **Supplementary Table 5b**). These results demonstrate that cCREs are enriched for functional activity and that their boundaries effectively delineate regions of regulatory potential.

Additionally, for MPRA regions that fell entirely within cCREs, those that overlapped cCREs and had high chromatin accessibility were, on average, almost three times more likely to be active than those that overlapped cCREs with low levels of chromatin accessibility (**Supplementary Figure 5c,d**, **Supplementary Table 5b**). These results highlight the high specificity of the Registry of cCREs for capturing cell type-specific regulatory activity.

***CRISPRi-FlowFISH***

CRISPRi-FlowFISH is a high-throughput experimental method that combines CRISPR interference, RNA fluorescence in situ hybridization (FISH), and flow cytometry to systematically perturb distal regulatory elements in the genome, enabling quantitative assessment of their effects on target gene expression. The Sabeti lab used CRISPRi Flow-FISH screens with HCR-FlowFISH to identify CRE-gene pairs around 20 genes in K562 cells (**Supplementary Table 5c**)[27]. Guide RNAs were selected using two approaches (1) targeting features associated with CREs such as chromatin accessibility and histone mark peaks and (2) dense tiling to detect elements with functional effects in an unbiased manner. The second approach, which was done for 18 of the experiments, enabled us to evaluate enrichment of

K562 cCREs, all cCREs and non-cCRE regions in functionally active elements (**Supplementary Figure 5ef**).

The Engrietz lab used CRISPRi Flow-FISH screens with PrimeFlow readout[57] to identify CRE-gene pairs around 33 genes in K562 cells and, for some genes, up to four additional cell lines (**Supplementary Table 5c**). These experiments targeted DNase peaks and therefore nearly all tested elements were cCREs. One caveat to consider when interpreting results from either set of experiments is that a negative result in CRISPRi FlowFISH does not necessarily indicate that a cis-regulatory element (CRE) is inactive; it may regulate a gene that was not included in the measurement.

##### *Additional CRISPR perturbations*

Over 66 additional CRISPR-based experiments were performed by functional characterization centers during ENCODE4, 53 in human and 13 in mouse (**Supplementary Table 5c**). While there was a lot of variation in experimental design such as utilizing different CRISPR constructs—cutting, inhibition, activation—and guide RNA design approaches—full tiling vs targeting peaks—we developed a systematic approach for characterizing cCRE activity in each of these assays. Briefly, we intersected cCREs with guide RNAs and quantified per cCRE levels in each experiment and then determined statistical significance using DESeq2. Results are available to view and download available on SCREEN.

#### 2.2 Comparing whole genome STARR-seq datasets

The White and Reddy labs both generated whole-genome STARR-seq data as part of the ENCODE4 project, including on the K562 cell line. Although both labs followed similar protocols, we observed notable differences in the STARR scores of cCREs calculated from the lab-specific datasets, which we attribute to subtle differences between methodologies. The correlation of STARR scores between the two datasets was modest (**Supplementary Table 5d**, Pearson correlation coefficient=0.25) when considering all cCREs with at least ten reads in both input DNA libraries (N = 507,547, $p < 2.2 \times 10^{-16}$). We further compared the STARR scores across different classes of cCREs, particularly those with biochemical signatures in K562 cells (i.e., K562 cCREs). STARR scores were significantly correlated for K562 distal enhancers (N =

4,085, R = 0.50, $p < 2.2 \times 10^{-16}$), but not for K562 promoter cCREs (N = 382, R = –0.01, p = 0.9).

We hypothesize that these differences may be due to variation in GC content, given our previous findings that GC content was the primary sequence feature distinguishing promoter and enhancer cCREs. To test this, we compared the correlation between STARR scores for K562 distal enhancer cCREs with either high or low GC content (**Supplementary Table 5d**). We observed that cCREs with high GC content had a lower correlation between lab datasets compared to those with lower GC content (correlation coefficients of 0.23 vs. 0.59, respectively). Similarly, high GC content promoters had a lower correlation between lab datasets compared to those with lower GC content (correlation coefficients of 0.14 vs. 0.63, respectively).

Due to the discrepancies between assays, we prioritized the data from each lab for specific types of analyses. Since the White lab performed STARR-seq across multiple cell types, we used their data to analyze cell type specificity of cCREs and identify sequence features predictive of activity (**Figure 2d,e**, **Extended Data Figure 3**). Conversely, because the Reddy lab data appeared less influenced by GC content, we used it to compare activity across elements (**Figure 2c**), assess combinatorial effects (**Figure 2f,g**), and identify potential silencers (**Figure 3h**, **Extended Data Figure 5,6**).

Despite the differences in the assays, we consistently observed that the GFI1B motif was significantly associated with silencer activity in both datasets. In particular, cCREs with the GFI1B motif showed lower-than-expected activity in both the White lab (–0.23 vs. –0.11, Wilcoxon test, $p < 2.2 \times 10^{-16}$) and Reddy lab (–0.16 vs. –0.04, Wilcoxon test, $p < 2.2 \times 10^{-16}$) data. This convergence across datasets suggests that, despite methodological differences, common core regulatory features such as the GFI1B binding may underlie silencer activity.

These results underscore the robustness of certain regulatory elements across varying experimental conditions and highlight the importance of considering both shared and distinct characteristics in interpreting STARR-seq data. Future work should focus on resolving methodological differences and developing unified models to better capture the complex regulatory landscape reflected in these assays.

##### 2.3 Combinatorial effects of cCRE activity

Using the CAPRA pipeline, we assessed the combinatorial effects of cCREs by comparing the activity of individual cCREs, derived from solo fragments (which span a single cCRE in its entirety), to that of double fragments (which span two cCREs in their entirety). Due to the strict overlap requirements, only a maximum of 26% cCREs were profiled with double fragments per experiment. We observed that the distribution of STARR scores for double fragments had a lower mean and smaller standard deviation compared to that of solo fragments (**Supplementary Figure 7a,** Wilcox test, $p < 2.2 \times 10^{-16}$), indicating potential fragment length biases in the STARR-seq data. Despite this, we chose not to normalize CAPRA STARR scores for fragment length, as our primary interest was in identifying instances of increased combinatorial activity between pairs. We opted for a more conservative approach to avoid overestimating such combinatorial effects.

When comparing STARR scores for cCREs using solo fragments versus double fragments, we found that activity levels for the majority of tested cCREs were correlated. This correlation strengthened as we applied more stringent thresholds for the minimum number of input DNA fragments, increasing from 0.16 for a minimum of one fragment to 0.42 for a minimum of ten fragments (**Supplementary Figure 7b-d**). However, there were notable exceptions where cCREs showed either higher or lower than expected activity. An example of higher-than-expected activity is highlighted in **Figure 2f-h** and discussed in the main text. Here, we focus on cases with lower-than-expected combined activities.

We identified 1,275 cCRE pairs with lower-than-expected combined activity, defined as instances where one cCRE exhibited high activity as a solo fragment (a positive STARR score with $p < 0.05$), but displayed lower activity when combined with another cCRE (double fragments), the STARR score dropped below zero. We then analyzed sequence motifs overlapping each of these cCREs. The cCREs that exerted a dominant repressive effect were enriched for motifs associated with transcription factors, some of which are known to have repressive activity, including GFI1B[45], MEF2D[84], and ZNF121[85] (**Supplementary Table 6c**).

One notable example includes two distal enhancer cCREs, EH38E3837011 and EH38E3837012 (**Supplementary Figure 7e,f**). These elements are located approximately 8 kb downstream and within the body of the *TTPA* gene, which encodes the alpha tocopherol transfer protein that regulates vitamin E levels. *TTPA* is primarily expressed in the liver, and both EH38E3837011 and EH38E3837012 display enhancer signatures exclusively in liver-derived biosamples. EH38E3837011 has a STARR score of 1.05 ($p$ = 0.005), while EH38E3837012 has a STARR score of –0.31. When tested in combination, the combined region also displays low activity, with a STARR score of –0.28, suggesting that EH38E3837012 exerts a dominant repressive effect on EH38E3837011. We hypothesize that this repression may be due to EH38E3837012 overlapping motifs for transcription factors with known repressive functions, including SMAD3[86], MAFK[87], and TCF7L2[88].

These findings highlight the complex interplay between cCREs, particularly in cases where combinatorial effects result in repressive activity that cannot be predicted from individual cCREs alone. Future work will prioritize a larger-scale investigation of such effects by designing STARR-seq experiments with varying fragment lengths to identify additional synergistic or antagonistic cCRE pairs. Additionally, we plan to explicitly test key hypotheses, such as the role of repressive motifs in modulating enhancer activity, through targeted experiments. This approach will help refine our understanding of the regulatory landscape and the mechanisms by which cCREs interact to influence gene expression.

#### Supplementary Note 3 - REST Silencers

##### 3.1 Defining REST+ cCREs

To refine our analysis of REST+ cCREs and their potential roles as silencers, we excluded REST+ promoter cCREs from our silencer annotations. In general, these elements did not exhibit characteristics consistent with neuron-restrictive silencer elements (NRSEs). REST+ promoter cCREs demonstrated significantly higher than expected STARR scores, with a median value of 0.6 compared to 0.11 for REST– promoter cCREs and –0.02 for all cCREs (Wilcoxon test, $p < 2.2 \times 10^{-16}$). Additionally, genes near REST+ promoter cCREs had higher expression levels (median TPM 16.8) than other annotated promoter cCREs in K562 cells (median TPM 11.7). These findings are inconsistent with a repressive role, and therefore, REST+ promoter cCREs were excluded from further analyses.

In contrast, REST+ proximal enhancer cCREs displayed a range of functional activities, reflecting their diverse regulatory roles. While some of these cCREs showed characteristics akin to silencer elements, others exhibited active behavior, similar to REST+ promoter cCREs. To address this variability, we applied an additional filtering criterion: REST+ proximal enhancer cCREs were only classified as silencers if they demonstrated silencer activity in STARR-seq assays (i.e., classified as a robust STARR-silencer). This approach ensured that we identified cCREs with clear functional evidence of repression.

##### 3.2 Chromatin enrichment analysis of REST+ enhancers

REST+ enhancer cCREs lacked typical enhancer signatures in the cell types where REST was bound. To identify the biosamples in which these cCREs function as enhancers, we analyzed the enrichment of H3K27ac signals in REST+ enhancer cCREs. Notably, REST+ enhancer cCREs showed higher-than-expected H3K27ac signals in brain tissues and neuron-related biosamples (**Supplementary Table 7h**), indicating functional activity in neurons, as demonstrated by transgenic mouse enhancer assays.

REST+ enhancer cCREs were also enriched for H3K27ac signals in endocrine tissues, such as the pancreas (**Supplementary Table 7h**), where REST plays a critical role in regulating pancreatic cell differentiation. For example, Rovira *et al*. demonstrated that REST inhibits pancreatic endocrine differentiation during embryonic development by repressing the formation of endocrine cells from progenitor cells[89]. Finally, we observed enrichment for H3K27ac signal in HCT116 cells, a colon cancer cell line (**Supplementary Table h**). This included samples that were part of the ENCODE4 degron series, demonstrating the reproducibility of this result across multiple HCT116 experiments. Although the enrichment of H3K27ac signals in HCT116 cells is intriguing, the functional role of REST+ enhancer cCREs in this colon cancer cell line is not yet fully understood, and future studies could focus on elucidating the specific mechanisms through which REST+ enhancers contribute to gene regulation in HCT116. Overall, our findings suggest that REST+ enhancer cCREs function as enhancers in neurons, endocrine cells, and the HCT116 cell line but act as silencers in other cell types.

#### Supplementary Note 4 - Identifying additional silencer cCREs and comparing them with existing collections of silencer elements

##### 4.1 CRISPRi-FLOWFish validation of STARR-silencers

Two STARR-silencer cCREs were tested in CRISPRi-FlowFISH experiments in K562 cells. The first, EH38E4193243, is a proximal enhancer for the *RTBDN* gene. As previously mentioned, EH38E4193243 is a REST+ enhancer/silencer and a STARR-silencer (**Main Text**), and targeting it with CRISPRi resulted in increased expression of *PRDX2* (effect size = –0.026 , FDR = 0.002) suggesting endogenous silencer activity.

Interestingly, perturbing nearby cCRE EH38E3291318, resulted in a similar decrease in *PRDX2* expression (effect size = –0.030 , FDR = 0.0004). In K562 cells, EH38E3291318 is a CA-CTCF cCRE that mediates 3D chromatin interactions. These CTCF-mediated interactions include a 14 kb loop with silencer cCRE EH38E4193243 and a 130 kb loop with another CA-CTCF cCRE EH38E1939855 (**Figure 5f**). The long range interaction between EH38E3291318 and EH38E1939855 creates a domain encompassing seven actively expressed genes and 82 active cCREs in K562 (**Figure 5f**). Within this CTCF-mediated domain, silencer EH38E4193243 is positioned near active promoters, which is evidenced by by a RNAPII ChIA-PET link connecting it with the promoter of *JUNB* (**Figure 5f**). Therefore the CRISPRi-FLOWFish results demonstrate that inhibiting either the silencer activity of EH38E4193243 or the 3D interactions maintained by CA-CTCF cCRE EH38E3291318 results in an increase in expression of *PRDX2*.

Despite silencer EH38E4193243 being proximal to the *JUNB* promoter in three dimensional space, there were no significant differences in *JUNB* expression when either silencer EH38E4193243 or CA-CTCF cCRE EH38E1939855 was perturbed (**Figure 5e**). This could be due to several factors such as the much higher overall expression or false negative interactions. Understanding the rules governing such selectivity and specificity warrant future investigation and the Registry of cCREs serves as the ideal framework for such future studies.

The other STARR-silencer cCRE tested by CRISPRi-FLOWFish was EH38E3019358. EH38E3019358 is a CA-TF cCRE in K562. It overlaps ChIP-seq peaks for erythroid transcription factors GATA1 and TAL1, as well as transcription factors with known repressive activities including TCF12[90], ZEB2[91], and IKZF1[92] (**Supplementary Table 8e**). While we did not observe any significant changes in gene expression when this cCRE was targeted, this could be due to its target gene not being assayed in these particular CRISPRi-FLOWFish experiments. The two genes measured at this locus were *NFE2* (85 kb away) and *ITGA5* (209 kb away) and were not linked to EH38E3019358 via any 3D chromatin interactions. Instead, possible target genes for EH38E3019358 may include *SMUG1*, *CBX5*, or the *HOXC* locus which are all connected to EH38E3019358 via Hi-C loops or ChIA-PET interactions (**Supplementary Table 8e**).

##### 4.2 Overlap of silencer cCREs with previously published collections

Our collection of silencer cCREs has fairly low overlap with previously published collections of silencers, suggesting that we have identified thousands of novel silencers in the human genome. Jayavelu *et al.*[49] tested 7,539 "uncharacterized CREs"—that is open chromatin regions that were at least 2.5kb from a TSS and did not overlap either a H3K4me3 or CTCF ChIP-seq peak—in an MPRA/STARR-seq assay, which we refer to as *Jayavelu elements*. They identified candidate silencers by selecting all elements with a mean activity level below control regions, resulting in a total of 2,997 candidate silencers, which we refer to as *Jayavelu silencers*. In total, 83% of Jayavelu elements and 81% of Jayavelu silencers overlapped cCREs. Of the 2,586 cCREs that overlapped Jayavelu silencers, only 232 (9%) were annotated as silencer cCREs (**Extended Data Figure 6a,b**). However, these 2,586 cCREs did have lower STARR scores than cCREs that overlapped non-silencer Jayavelu elements (median STARR score of -0.06 vs 0.14, $p < 2.2 \times 10^{-16}$) and randomly selected cCREs (-0.06 vs. -0.02, $p < 2.2 \times$

$10^{-16}$), suggesting that we may identify concordant STARR-silencer cCREs with less stringent thresholds. It is also worth noting that 250 silencer cCREs overlapped Jayavelu elements and 232 (93%) were Jayavelu silencers. Stratifying by specific categories of silencer cCREs, 100% of tested stringent STARR-silencer cCREs, 95% of tested robust STARR-silencer cCREs, 96% of REST+ silencer cCREs, and 90% of REST+ enhancer/silencer cCREs were called as Jayavelu silencers (**Extended Data Figure 6a**).

Pang *et al.*[50] tested 177,046 open chromatin regions identified through FAIRE-seq, which we refer to as *Pang elements,* in a repressive ability of silencer elements (ReSE) functional screen. They identified 2,662 candidate silencers, which we refer to as *Pang silencers*. In total, 56% of Pang elements and 57% of Pang silencers overlapped cCREs. Of the 1,793 cCREs that overlapped Pang silencers, only 11 (0.6%) were annotated as silencer cCREs. These 1,793 cCREs actually had slightly higher STARR scores than cCREs that overlapped non-silencer Jayavelu elements (median STARR score of 0.06 vs. 0.05, $p = 0.07$) and significantly higher STARR scores than randomly selected cCREs (0.06 vs. -0.02, $p < 2.2 \times 10^{-16}$). Of the 624 silencer cCREs that overlapped Pang elements, only 11 (93%) were Jayavelu silencers.

Huang *et al.*[48] identified silencers by selecting DNase hypersensitivity sites that overlapped H3K27me3 and were negatively correlated with gene expression. They identified 8,084 candidate silencers, which we refer to as *Huang silencers*. In total, 93% of Huang silencers overlapped cCREs. Of the 22,465 cCREs that overlapped Huang silencers, only 135 (0.6%) were annotated as silencer cCREs. Additionally the 22,465 cCREs that overlapped Huang silencers had significantly higher STARR scores than randomly selected cCREs (0.02 vs. -0.02, $p < 2.2 \times 10^{-16}$).

Cai *et al.*[52] identified silencers by annotating H3K27me3-rich regions which they named MMRs. They identified 1,677 MMRs, which we refer to as *Cai silencers*. In total, nearly 100% of Cai silencers overlapped cCREs. Of the 222,109 cCREs that overlapped Pang silencers, 1,138 (0.5%) were annotated as silencer cCREs. Additionally the 1,138 cCREs that overlapped Cai silencers had significantly higher STARR scores than randomly selected cCREs (0.004 vs. -0.02, $p < 2.2 \times 10^{-16}$).

Overall, our analysis indicates that silencer cCREs identified through our approach represent a distinct class of regulatory elements compared to previous studies. While Jayavelu *et al.*'s use of a similar MPRA/STARR-seq-based method resulted in a higher degree of overlap, the divergence seen with other datasets, such as those from Pang et al, Huang et al, and Cai et al, could reflect either technical biases in silencer identification strategies, biological variability across cell types and conditions, or multiple classes of silencer-like elements. These findings underscore the need for further investigation to reconcile these differences and better understand the diverse mechanisms of gene repression, potentially through additional experimental validation and integration of multi-omics datasets.

##### 4.3 Chromatin accessibility enrichment

Only 14% of silencer cCREs had high levels of chromatin accessibility in K562 cells (z-score > 1.64, **Extended Data Figure 6c**). Stratifying by subclasses, 32% of REST+ enhancer/silencer cCREs had high chromatin accessibility, followed by REST+ silencer cCREs (17%), robust STARR-silencer cCREs (8%) and stringent STARR-silencer cCREs (6%). Silencer cCREs did have slightly higher levels of chromatin accessibility in K562 compared to randomly selected cCREs (median z-score 0.26 vs. -0.08, Wilcoxon test, $p < 2.2 \times 10^{-16}$). These results suggest silencer cCREs maintain a baseline level of accessibility that may be sufficient for transcription factor binding, even if not as broadly accessible as promoters or enhancers.

Since silencer cCREs had low chromatin accessibility in the cell type in which they act as silencers—whether defined by STARR-activity or REST binding—we were interested in determining in which cell types these cCREs had DNase signals. Compared to non-silencer cCREs, silencer cCREs were enriched for chromatin accessibility in early-stage cell types such as pluripotent cell lines (ESCs and iPSCs) and some *in vitro* differentiated cell lines (**Extended Data Figure 6d,e**, **Supplementary Table 8f**). Tissues from fetal samples were more likely than adult tissues to have open chromatin at silencer cCREs, presumably because fetal tissues comprise fewer mature cell types (**Extended Data Figure 3d,e**). This pattern was particularly significant for REST+ silencer cCREs (**Extended Data Figure 3f, Supplementary Table 8g**). Fetal tissues had significantly higher chromatin accessibility at REST+ silencer cCREs compared to matched adult tissue, except for brain tissues, where REST+ silencer cCREs had low chromatin accessibility in both fetal and adult tissues.

There were also cancer cell lines, such as Ishikawa (endometrial adenocarcinoma) and T47D (breast cancer), that showed enriched chromatin accessibility for silencer cCREs (**Extended Data Figure 6de**, **Supplementary Table 8f**). This may be due to the tendency of certain cancers to revert to a more pluripotent-like state, characterized by the reactivation of developmental pathways[93] and perhaps a broader chromatin landscape. Future work will aim to determine whether these changes in chromatin accessibility at silencers are specific to particular cancer types and to assess their functional impact on gene regulation and cellular behavior.

#### Supplementary Note 5 - Overlap of the Registry of cCREs with other classes of Encyclopedia annotations

The ENCODE Encyclopedia comprises four levels of annotations: sequence, elements, genes, and interactions. The Registry of cCREs is an integral component of the element annotations and, when combined with other annotations, provides a more comprehensive picture of gene regulation by capturing the interplay between regulatory elements and their functional contexts.

##### 5.1 Chromatin states (elements)

We previously demonstrated that cCREs and chromatin states, such as those generated by ChromHMM, are highly concordant[1]. These two approaches for identifying cis-regulatory regions are complementary, with cCREs providing high spatial resolution and chromatin states providing finer-grained histone mark combinations. Annotating cCREs with chromatin states can reveal subtle differences in regulatory element activity, while maintaining high resolution elements. For example, Borsari *et al.*, integrated temporally derived chromatin states with cCREs to study the transdifferentiation of human pre-B cells into macrophages and discovered that a precise ordering of histone modifications was required for gene activation[94].

Therefore, to complement the Registry of cCREs, we generated ChromHMM states for 330 epigenomes consisting of H3K4me3, H3K27ac, H3K4me1, H3K36me3, H3K27me3, and H3K9me3 histone marks (**Supplementary Table 9**). Because we observed differences in signal thresholding between models trained on ChIP-seq or Mint ChIP-seq data, we trained two assay-specific models (**Supplementary Methods**). We trained a ChIP-seq model on 20 epigenomes, which were generated across all project stages, and used the model to generate chromatin states for 210 epigenomes. For Mint ChIP-seq data, we trained the model on 20 epigenomes derived from a combination of primary cells, *in vitro* differentiated cells and cell lines, which we used to generate chromatin states for 120 epigenomes. These chromatin states can be viewed and are available for download on SCREEN.

##### 5.2 ENTEx (sequence & elements)

The EN-TEx resource mapped functional genomic datasets to diploid genomes from four individuals, revealing over a million allele-specific loci, demonstrating the importance of local nucleotide sequences, and enabling improved predictive modeling of eQTLs in challenging tissues[95]. As part of the EN-TEx resource we annotated cCREs with allele-specific biochemical activity and tissue-specific activity. Here we updated these annotations for V4 cCREs.

Of the 514,427 cCREs overlapping variants that were heterozygous in at least one of the four ENTEx donors, 26% (131,215) had allelic imbalance for either one of the six surveyed histone marks (H3K4me3, H3K4me1, H3K27ac, H3K36me3, H3K27me3, H3K9me3) or one of the three DNA binding proteins (CTCF, EP300, POLR2A). For 28 tissues, we also annotated active cCREs via overlap with the histone modifications H3K4me1, H3K4me3 and H3K27ac, annotating 31% (N = 717,095) of V4 cCREs. These annotations are available to view and download via SCREEN.

##### 5.3 Annotated GENCODE genes (genes)

We identified promoter cCREs for 90% of protein coding GENCODE genes (**Supplementary Figure 2a**). Over coverage of lncRNA genes, which tend to be much more cell type specific (Gini coefficient of 0.56 compared to 0.37 for protein coding genes, $p < 2.2 \times 10^{-16}$ ), was less at only 57%. Within specific cell types—specifically deeply profiled cell lines—an average of

82% of expressed genes (TPM > 1, as measured by short read RNA-seq) have an annotated promoter cCRE in that cell type (**Supplementary Table 10a**). This percentage increased to 90% and 92% when we analyzed genes with higher expression levels of 5 and 10 TPM, respectively. These results demonstrate that the Registry of cCREs covers the majority of promoters for genes expressed in the included cell types.

We then compared the expression of four classes of genes: (1) genes with an active promoter cCRE (high DNase and high H3K4me3) with high H3K27ac signal, (2) genes with an active promoter cCRE that had low H3K27ac signal, (3) genes with an inactive promoter cCRE (low DNase and low H3K4me3), and (4) genes without a promoter cCRE (**Supplementary Table 10b**). Across all 13 tested cell types, genes with active promoter cCREs with high H3K27ac signal had the highest expression with an average median TPM of 10.3. Genes with promoter cCREs that had low H3K27ac signal but high chromatin accessibility and H3K4me3 signals in the tested cell type had lower levels of expression with average median TPMs of 2.0, suggesting these promoter cCREs are likely poised with basal levels of transcription. Genes in the last two groups were generally not expressed with average median TPMs < 0.1.

#### 5.4 Novel transcripts (genes)

Our previous work demonstrated that while the GENCODE annotations are fairly comprehensive, transcription assays that capture 5’ ends of transcripts such as RAMPAGE and long read RNA-seq identify novel TSSs, particularly those for highly tissue specific transcripts[96].

##### RAMPAGE

We previously curated a set of 50 thousand transcription sites by integrating RAMPAGE data across 115 cell and tissue types. We classified these transcription sites, which we referred to as RAMPAGE representative peaks (rPeaks), based on their overlap with current GENCODE annotations and the genomic locations of the 3' ends of paired end reads. TSSs of annotated GENCODE genes, whether verified or novel, had high overlap with cCRE (96 and 92%, respectively, **Supplementary Table 10c**). TSSs of novel transcripts had slightly lower overlap (86%) and rPeaks that we deemed not be true TSSs and rather the result of local transcription had the lowest overlap with cCREs at 71%.

**Long read RNA-seq**

The Mortazavi lab generated 98 long read RNA-seq experiments from human biosamples and 65 from mouse biosamples[97]. Most samples (N = 137) were sequenced on the Pacific Biosciences (PacBio) Sequel II platform with a smaller number sequenced on the original PacBio Sequel platform (N = 23) and limited number sequenced on the Oxford Nanopore MinION platform (N = 3). These datasets were complemented by 8 long read datasets generated by the LRGASP project which assayed three human biosamples (6 experiments) and one mouse biosample (2 experiments) using different library construction methods and sequencing platforms.

For each long read experiment, we generated TSS clusters by merging 5' ends of all reads within 25 bp of one another. We then selected all clusters with at least 5 reads per million (RPM). This resulted in an average of ~14,000 TSS clusters per human experiment and ~13,000 TSS clusters per mouse experiment (**Supplementary Table 10d, e**). On average, 87% of human TSS clusters were within 200 bp of a human cCRE center and 81% of mouse TSS clusters were within 200 bp of a mouse cCRE center. We observed differences in human cCRE overlap depending on the sequencing platform with an overlap of 87% of TSS clusters from PacBio Sequel II data, 90% of TSS clusters from PacBio Sequel data and 80% of TSS clusters from Oxford Nanopore MinION data. These results support previous findings that the PacBio platform is better for identifying 5' ends of transcripts[98]. We also hypothesize that TSS clusters that are not within 200 bp of a cCRE center are likely false positives as they are less likely to also overlap PRO-cap peaks (**Supplementary Table 10d**). On average 68% of TSS clusters that overlap a cCRE also overlap a PRO-cap peak in that same cell type compared to only 2% of TSS clusters that do not overlap cCREs (Fisher's Exact Test, $p < 2.2 \times 10^{-16}$).

#### 5.5 Sites of nascent transcription (elements)

The Lis and Yu labs conducted 24 PRO-cap experiments to measure transcriptional initiation using cap-selection-based nascent RNA sequencing (**Supplementary Table 10f**). To evaluate the role of cap selection, they also performed two experiments in K562 cells without the cap selection step. In experiments with cap selection, an average of 98% of bidirectional PRO-cap peaks overlapped cCREs, compared to only 74% in experiments without cap selection. When restricting the analysis to cell type-specific cCREs, 90% of peaks from cap-selected experiments overlapped active cCREs in the corresponding cell type, compared to just 46% from experiments without cap selection. These findings highlight the critical importance of cap selection in the PRO-cap protocol and confirm that cCREs capture the vast majority of transcriptionally active sites in a given cell type.

#### 5.6 3D chromatin interactions (interactions)

##### ChIA-PET

The JAX lab generated 67 replicated ChIA-PET experiments—30 targeting RNAPII and 39 targeting CTCF—across a diverse set of cell and tissue types (**Supplementary Table 11a,b**). Prior to intersecting with cCREs, we selected all interactions that were reproducible between biological replicates with anchors greater than 8 kb apart, merging interactions with overlapping anchors. This resulted in an average of 23 thousand interactions for RNAPII experiments and 71 thousand interactions for CTCF experiments (**Supplementary Table 11a,b**).

Nearly all ChIA-PET interactions (>99%) had at least one anchor overlapping a cCRE, with most interactions (96%) having both anchors overlapping a cCRE (**Supplementary Table 11a,b**). However, when considering cell type-specific cCRE annotations, the percentages of both anchors overlapping a cCRE dropped to an average of 68% for RNAPII interactions and 50% for CTCF interactions (**Supplementary Table 11c**). This trend was particularly pronounced in H1 embryonic stem cells: over 80% of RNAPII and CTCF interactions had at least one anchor overlapping an active cCRE in H1 cells, but fewer than 35% of interactions had both anchors overlapping. These results suggest that some interactions involve regions with regulatory potential that lack active chromatin signatures in the surveyed cell type.

### Hi-C

The Aiden lab generated 136 intact Hi-C experiments, identifying an average of ~33,000 loops per experiment (**Supplementary Table 11d**). Consistent with ChIA-PET results, over 99% of Hi-C loops had at least one anchor overlapping a cCRE, and 97% had both anchors overlapping a cCRE. However, in specific cell types, only 43% of loops had both anchors overlapping cCREs active in those cell types (**Supplementary Table 11c**). These findings further support the idea that loops often connect active regulatory elements to regions that may be poised for activity.

##### 5.7 Functional interactions (interactions)

As previously mentioned (**Supplementary Note 2.1**), CRISPRi-FlowFISH is an experimental technique that perturbs CREs to assess their impact on gene expression, enabling systematic mapping and prediction of CRE-gene interactions. We integrated these functional interactions with the Registry of cCREs resulting in 233 cCRE-gene pairs corresponding to 197 cCREs and 15 genes, which can be found on SCREEN.

##### 5.8 Enhancer-gene interaction predictions (interactions)

The ENCODE Distal Regulation working group curated a resource of over 13 million enhancer-gene regulatory interactions across 352 cell types and tissues, integrating models, chromatin state measurements, 3D contacts, and genetic perturbation data[99]. We integrated the Registry of cCREs with this resource, making the results available on SCREEN. For biosamples in the *Core Collection*, we intersected cell type-specific cCREs with predicted interactions from cell-type-matched rE2G models and evaluated the percentage of cCRE with predicted genes. Though we performed the analysis for all classes of cCREs (**Supplementary Table 11e**), distal enhancer cCREs were of greatest interest. On average, 56% of distal enhancer cCREs were annotated with at least one target gene, with each enhancer linked to 1.9 genes on average. In 75% of cases, the predicted target corresponded to the nearest protein coding gene.

#### Supplementary Note 6 - SCREEN

We previously developed a web portal, SCREEN (Search Candidate cis-Regulatory Elements by ENCODE) to allow other researchers to query and visualize data from the Registry and the ENCODE Encyclopedia[1]. As part of this work, we redesigned and refactored the entire site to improve usability and performance while also adding new features and datasets (**Supplementary Figure 9**).

SCREEN allows users to search the Registry using a variety of methods, including (1) genomic region, (2) individual cCRE accessions, (3) SNP rsID, (4) gene name, or (5) cell/tissue type. SCREEN also supports users who are interested in exploring multiple regions through BED file upload and intersection. After searching, users are presented with an interactive table summarizing the cCREs that meet their search criteria. This table can be further filtered based on specific biosample activity, biochemical signal scores, cCRE conservation, or linked genes.

Clicking on individual elements in the results table allows users to see detailed information about each cCRE. Along with basic information such as coordinates and classification (e.g., promoter, proximal enhancer, CA-H3K4me3, etc), each cCRE details page also includes information about nearby genes and SNPs, gene expression, and overlap with transcription factor binding sites. New features include (1) ChromHMM states, which are presented via an interactive genome browser and table, (2) functional characterization data, including quantifications from our new CAPRA quantification method, and (3) expanded cCRE-gene links, which include data from 3D chromatin interactions, CRISPRi-FlowFISH experiments, and computational predictions. Alongside the search functionality, SCREEN also hosts files for users to download and use in their own analysis including the entire set of V4 cCREs, subsets of cCREs based on genomic region, classification, biosample, and biochemical signals, and large matrices containing normalized signals for each of the core biochemical marks across all biosamples and cCREs.

We developed the frontend of SCREEN using Next.js, an open-source React framework that provides robust support for server-side rendering. We have also heavily utilized components from Material UI, an open-source React component library, which implements Google's Material Design. The data for this project are stored in a Google Cloud storage bucket and Google Cloud PostgreSQL database. It is provided to our frontend through a GraphQL application program interface (API). All data on SCREEN can also be downloaded or programmatically accessed through our GraphQL API for downstream analysis.

#### Supplementary Note 7 - Dissecting the *RTBDN-MAST1* locus

##### 7.1 CRISPRi-FlowFISH testing of EH38E3291318

The Engreitz lab tested CA-CTCF cCRE EH38E3291318 in seven CRISPRi-FlowFISH experiments, measuring the readout of *JUNB*, *KLF1*, *LYL1*, *PRDX2*, *RAD23A*, *RNASEH2A*, and *WDR83OS*, two of which, *KLF1* and *PRDX2* had significant results. As previously mentioned (**Supplementary Note 4.1**, targeting EH38E3291318 resulted in a significant increase in expression of *PRDX2* (effect size = –0.03, p = $4.1 \times 10^{-4}$) where we hypothesized that inhibiting EH38E3291318 disrupted 3D chromatin interactions between the silencer cCRE EH38E4193243 and *PRDX2*. On the other hand, targeting EH38E3291318 resulted in the decreased expression of *KLF1* (effect size = 0.04, p = $6.7 \times 10^{-6}$). This duality in effect sizes aligns with the notion that disrupting EH38E3291318 interferes with the 3D chromatin architecture, rewiring enhancer-promoter interactions, thereby leading to contrasting regulatory outcomes for genes within the affected genomic neighborhood.

##### 7.2 Prioritizing additional variants for future testing

In addition to rs2290688, we identified two other potentially functional variants, rs2280742 and rs2072597 at the *RTBDN-MAST1* that may warrant future validation. Rs2280742 overlaps distal enhancer cCRE EH38E3291349, which is classified as an enhancer in myeloid-derived biosamples including K562, CD14+ monocytes, the promyeloblast cell line HL-60, and a very small subset of tissue biosamples, including heart, lung, and muscle (**Supplementary Figure 10a, Supplementary Table 12g**). It is adjacent to binding sites for transcription factors (**Supplementary Figure 10b**) and previous work identified allelic imbalance of transcription factor ChIP-seq reads at this locus[100], suggesting this variant may impact binding.

Rs2072597 is a missense variant in the second exon of KLF1, resulting in a substitution of Serine (TCC) with Proline (CCC) at the 102nd amino acid. Despite the substitution of two biochemically distinct amino acids, we hypothesize that this mutation does not significantly impact the function or stability of KLF1 for several reasons. First, the amino acid change occurs within a disordered region of the protein, rather than in a functional domain, such as one of the annotated zinc finger domains. Second, analysis of the Zoonomia 240-way alignment across mammals shows that this position is not conserved; proline is the most common residue at this site (137 species, including chimpanzee and gorilla), followed by serine (65 species), and threonine (25 species; **Supplementary Figure 10c**). This suggests that the substitution of Serine with Proline is tolerated and may not disrupt protein function across species. Finally, the allele frequency of this variant is relatively high, varying significantly between populations: 0.68 in Europeans compared to 0.39 in East Asians (**Supplementary Table 12h**). While this evidence suggests that the variant is likely benign, this amino acid change could still exert subtle effects on KLF1 stability or function. Therefore, additional validation studies are warranted to fully understand its impact.

We recognize that multiple variants at the *RTBDN-MAST1* locus may collectively influence KLF1 expression and function. Rs2290688, rs2280742, and rs2072597 are in high linkage disequilibrium with one another ($R^2 > 0.82$), suggesting that all three variants could exert small, compounding effects on the regulatory landscape. These effects may collectively contribute to the observed red blood cell traits. For instance, rs2290688 is associated with 3D chromatin interactions, rs2280742 overlaps a myeloid enhancer with evidence of transcription factor binding alterations, and rs2072597 represents a missense variant that could subtly impact protein stability or function. The interplay between these variants underscores the complexity of this locus and highlights the need for integrative analyses and functional validation to fully elucidate its roles in regulating *KLF1*.

#### Supplementary Note 8 - Additional Acknowledgements

##### 8.1 Members of the ENCODE4 cCRE working group:

Greg Andrews, Alex Barrera, Mike Beer, Brad Bernstein, Beatrice Borsari, Stephanie Calluori, You Chen, Alan Du, Shaimae Elhajjajy, Jesse Engreitz, Chuck Epstein, Kaili Fan, Weixiang Fang, Adam Frankish, Adam Frankish, Yu Fu, Idan Gabdank, Mingshi Gao, Mark Gerstein, Daniel Gilchrist, Will Greenleaf, Gamze Gursoy, Ben Hitz, Hongkai Ji, Graham Johnson, Brendan Kearney, Anshul Kundaje, Lazaros Lataniotis, Christina Leslie, John Lis, Mats Ljungman, Brian Magnuson, Georgi Marinov, Ariel McShane, Eyal Metzl Raz, Wouter Meuleman, Jill Moore, Stephanie Morris, Ali Mortazavi, Kristy Mualim, Abdullah Ozer, Mike Pazin, Henry Pratt, Soumya

Raychaudhuri, Milad Razavi-Mohseni, Thomas Reimonn, Xingjie Ren, Narges Rezaie, Marina Ruiz-Romero, Richard Sandstrom, Yin Shen, Mike Snyder, John Stam, Yi Wang, Eve Wattenberg, Annika Weimer, Zhiping Weng, Kevin White, Shannon White, Jin Woo Oh, Wang Xi, Jinrui Xu, Li Yao, Ingrid Youngworth, Haiyuan Yu, Michelle Yu, Ken Zaret, Junke Zhang, Lina Zheng

**8.2 Members of the ENCODE4 TSS annotation working group:**

Greg Andrews, Gabby Balderrama, Silvia Carbonell, Kelly Cochran, Daniel Kim, Anshul Kundaje, Julien Lagarde, Ishita Mangla, Jill Moore, Ali Mortazavi, Vivek Ramalingam, Milad Razavi-Mohseni , Liz Rebboah, Fairlie Reese, Jacob Schreiber, Laksshman Sundaram, Joshua Theisen, Jacob Tome, Diane Trout, Li Yao, Ingrid Youngworth

**8.3 Members of the ENCODE4 Biosamples working group:**

Ali Mortazavi, Annika Weimer, Ariel McShane, Aviva Presser Aiden, Barbara Wold, Beatrice Borsari, Ben Hitz, Bill Noble, Brad Bernstein, Briana Nuñez, Brian St. Hilaire, Brian Williams, Carrie Davis, Chad Nusbaum, Chuck Epstein, Cricket Sloan, Dan Gilchrist, Eileen Cahill, Elise Feingold, Erez Lieberman Aiden, Eric Mendenhall, Eva Gega, Fabio Navarro, Gabriela Balderrama Gutierrez, Haiyuan Yu, Huiya Gu, Idan Gabdank, Ishwarya Venkata, Ivan Bochkov, Jason Hilton, Jennifer Jou, Jenn Wineski, Jess Halow, Jill Moore, Jing Zhang, Joel Rozowsky, John Stamatoyannopoulos, Karan Bedi, Kathy Rader, Konor von Kraut, Laura Reinholdt, Liz Gaskell, Mark Gerstein, Mark Mackiewicz, Mats Ljungman, Meenakshi, Mike Pazin, Mike Snyder, Minji Kim, Neva Durand, Nina Farrell, Patrick Gallagher, Ping Wang, Raj Kaul, Rick Myers, Rob Spitale, Sam Moore, Saurabh Agarwal, Shin Lin, Stephanie Morris, Tim Reddy, Ulrike Litzenburger, Weiwei Zhong, Will Greenleaf, Yiing Lin, Yijun Ruan, Yunhai Luo, Zack Myers, Zhiping Weng

#### Supplementary Methods

##### Proximity of experimentally derived transcription start sites with rDHSs

We downloaded FANTOM CAGE[101] peaks from the Fantom Database and calculated their distance to the nearest rDHS using BEDTools closest[74]. We then calculated the percentage of CAGE peaks within 250 bp of an rDHS (**Supplementary Table 2a**). We performed the same analysis for RAMPAGE rPeaks[96] that were associated with GENCODE genes.

**Relevant scripts**: Supplementary-Table-2a.CAGE-RAMPAGE-Overlap.sh

##### Overlap of histone mark peaks with rDHSs

We downloaded peaks from the ENCODE portal for all experiments targeting H3K4me3, H3K27ac, and H3K4me1 (**Supplementary Table 2b-d**). We intersected these peaks with rDHSs using BEDTools intersect[74].

**Relevant scripts**: Supplementary-Table-2bcd.Histone-Peak-Overlap.sh

##### Overlapping transcription factor peaks with rDHSs

We downloaded peaks for all transcription factor experiments from the ENCODE portal that met the following criteria (**Supplementary Table 2e**):

- Transcription factor was sequence specific, i.e., has an annotated motif in HOCOMOCOv11[72]
- FRiP score (fraction of reads within peaks) > 0.3
- Peak files were labeled as "Preferred default peaks"

We intersected each of these 535 peak sets with rDHSs using BEDtools[74] and calculated the percentage of total overlapping peaks. We selected all peak sets with less than 90 percent overlap for additional analysis (N = 50).

**Relevant scripts:**

Supplementary-Figure-1a.Supplementary-Table-2e.TF-Overlap-Anchors-Justification.sh

#### Identifying and characterizing transcription factors with low overlap with rDHSs

For the 50 experiments, we selected all peaks that did not overlap rDHSs. Then, for each pair of experiment, we calculated the overlap coefficient between the non-rDHS peaks:

$$\text{overlap}(A, B) = \frac{|A \cap B|}{\min(|A|, |B|)}$$

We then performed hierarchical clustering using the default parameters of hclust to identify common sets of transcription factors.

For groups with at least three thousand sites, which included the MAFK/MAFF, ATF/CREB1 and CEBPB clusters, we calculated the average PhastCons 100-way vertebrate conservation scores across each region. We compared scores to all rDHSs, TF anchors, and non-cCRE regions. We calculated statistical significance using a Wilcox test.

**Relevant scripts**: Supplementary-Figure-1b.TF-Overlap-Matrix.sh
Supplementary-Figure-1c.Conservation.sh

#### Calling Multi-mapping cCREs

We called DHSs with the Hotspot2 peak caller[102] using unfiltered BAM alignment files from the ENCODE portal (**Supplementary Table 3c**). Instead of calling DHSs genome wide, we called DHSs on a subset of the genome centered on high confidence ATAC-seq peaks that did not overlap rDHSs (**Supplementary Table 3a,b**). Using our rDHS pipeline, clustered the multi-mapping DHSs and identified representative elements which we called multi-mapping cCREs.

**Relevant scripts**: https://github.com/weng-lab/dnaseseq-workflow

#### Characterizing the expansion of the Registry

We stratified ENCODE4 cCREs into three categories:

1. Replicated from ENCODE3. These are cCREs that are present in both versions of the Registry. These cCREs can either have the same exact position, in which case their accession remains the same, or their boundaries can slightly shift.
2. Called in ENCODE3 but filtered out due to incomplete data. We previously filtered out cCREs deriving from biosamples lacking certain combinations of data (see Supplementary Note 1 from [1]). However, with our expansion to include new classes of cCREs, we now include these elements in our Registry.
3. New in ENCODE4 are cCREs never previously called.

We characterized these three groups by intersecting them with whole genome STARR-seq peaks from K562 (ENCFF908UFR) and calculating the average per-base conservation across each cCRE using 240-way Mammalian phyloP scores from the Zoonomia consortium

To identify which biosamples contributed most to the increased Registry, we calculated chromatin accessibility enrichment for groups 2 & 3 (called in ENCODE3 and new in ENCODE4). In each biosample, we determined the fraction of new V4 cCREs with high signal and compared it to the fraction of all cCREs with high signal.

**Relevant scripts:** Supplementary-Figure-3ab.Previous-Version-Comparisons.sh
Supplementary-Figure-3c.Conservation-Previous-Versions.sh
Supplementary-Table-4a.DNase-Enrichment-V4.sh

#### Overlap with BICCN cCREs

We downloaded BICCN cCRE annotations from Li *et al.*[32] via catlas.org, which contained a consensus set of elements and elements called for individual cell types (N = 107). Using BEDTools, we intersected each peak set with three versions of cCREs: ENCODE3 cCREs (V2, 2020), a preliminary version of ENCODE4 cCREs (V3, 2022), and our updated ENCODE4 cCREs (V4, 2024). We then calculated the percentage of peaks that overlapped cCREs for each version.

To identify which cCREs contributed to the increased overlap, we calculated chromatin accessibility enrichment for newly added V4 cCREs that overlapped BICCN cCREs. For each biosample, we determined the fraction of overlapping V4 cCREs with high signal and compared it to the fraction of all cCREs with high signal. Biosamples were ranked based on enrichment and stratified by their tissue or organ of origin.

**Relevant scripts:** Supplementary-Table-4b.BICCN-Overlap.sh
Supplementary-Table-4c.DNase-Enrichment-BICCN.sh
Supplementary-Figure-3def.Expanded-Collection-Enrichment.sh

#### Overlap with Loupe *et al.* union peaks

We downloaded Supplementary Table 4 from Loupe *et al.*,[33] which contains the union of peaks called for individual brain regions and cell types. For each brain region and cell type combination (N = 12), we generated BED files of the peak locations. Using BEDTools, we intersected each peak set with three versions of cCREs: ENCODE3 cCREs (V2, 2020), a preliminary version of ENCODE4 cCREs (V3, 2022), and our updated ENCODE4 cCREs (V4, 2024). We then calculated the percentage of peaks that overlapped with cCREs for each version.

To identify which cCREs contributed to the increased overlap, we focused on the union peaks from Dorsolateral Prefrontal Cortex (DLPFC) NeuN+ cells. Specifically, we calculated chromatin accessibility enrichment for newly added V4 cCREs that overlapped these union peaks. For each biosample, we determined the fraction of overlapping V4 cCREs with high signal and compared it to the fraction of all cCREs with high signal. Biosamples were ranked based on enrichment and stratified by their tissue or organ of origin.

**Relevant scripts:** Supplementary-Table-4d.Loupe-Myers-Overlap.sh
Supplementary-Table-4e.DNase-Enrichment-Loupe-Myers.sh
Supplementary-Figure-3def.Expanded-Collection-Enrichment.sh

#### Classification of STARR+ distal enhancers

We classified STARR+ distal enhancers specific to K562 and HepG2 cells based on their biochemical signatures as defined in the cCRE workflow. Each STARR+ region was assigned to one of the following cCRE classes: distal enhancer (dELS), CA-H3K4me3, CA-CTCF, CA-TF, CA, or inactive. For each cell type, we calculated the fraction of STARR+ cCREs in each class.

To assess differences in cCRE classifications between K562 and HepG2, we used Fisher's exact test. This analysis was performed for all K562 and HepG2 STARR+ distal enhancers and for subsets of these enhancers that overlapped motifs for transcription factors GATA1, HNF4A, p53, and GFI1B.

**Relevant scripts:** Supplementary-Figure-6.cCRE-Classification.sh

#### Gene expression near candidate silencers

For each group of cCREs, we identified the closest protein coding gene, as defined by linear distance to the transcription start site, using BEDTools. We then calculated expression of these genes in transcripts per million using K562 total RNA-seq data from the ENCODE portal (ENCFF421TJX). We performed Wilcox test to evaluate significance.

**Relevant scripts:** Supplementary-Figure-8c.Silencer-Gene-Expression.sh

#### Enrichment for transcription factor binding site at STARR-silencers

To evaluate transcription factor enrichment at STARR-silencers, we calculated the enrichment of transcription factor motifs (**Supplementary Table 8b**) and ChIP-seq peak summits (**Supplementary Table 8c)** across four groups of cCREs:

- Stringent STARR-silencers, negative log2 fold change and $p < 0.01$
- Robust STARR-silencers, negative log2 fold change and $p < 0.05$
- Active cCREs, positive log2 fold change and $p < 0.01$

- Neutral cCREs, -0.1 < log2 fold change < 0.1

For motif analysis, we quantified the number of cCREs in each group overlapping transcription factor motifs identified by FIMO (see **Methods**). We assessed statistical significance using Fisher's Exact Test with FDR correction. For ChIP-seq data, we intersected peak summits with cCREs using BEDTools, calculating the number of cCREs in each group overlapping each TF summit, and determined significance using Fisher's Exact Test with FDR correction.

For transcription factors significantly enriched in STARR-silencers (FDR < 0.05 for all comparisons), we identified overlapping cCREs and computed pairwise Overlap Coefficients (see **Supplementary Methods**). These coefficients were used for hierarchical clustering (via hclust) to identify transcription factors with common binding sites.

**Relevant scripts:** Supplementary-Table-8b.STARR-Silencer-Motif-Enrichment.sh
Supplementary-Table-8c.STARR-Silencer-TF-Enrichment.sh
Supplementary-Figure-8d.STARR-Silencer-TF-Enrichment.sh

##### Phylogenetic analysis of *KLF1* missense mutation

We downloaded the 241-way mammalian multiple sequence alignment and phylogenetic tree from the Zoonomia consortium[103–105]. We next used hal2maf from the Comparative Genomics Toolkit[106] to extract the alignment at the *KLF1* missense mutation. We then parsed the resulting MAF file and converted each codon to its corresponding amino acid. Finally, we plotted the phylogenetic tree colored by amino acid in R using the ggtree library[107].

**Relevant scripts:** klf_hal2maf.sh
parse_klf_maf.py
plot_klf_tree.R

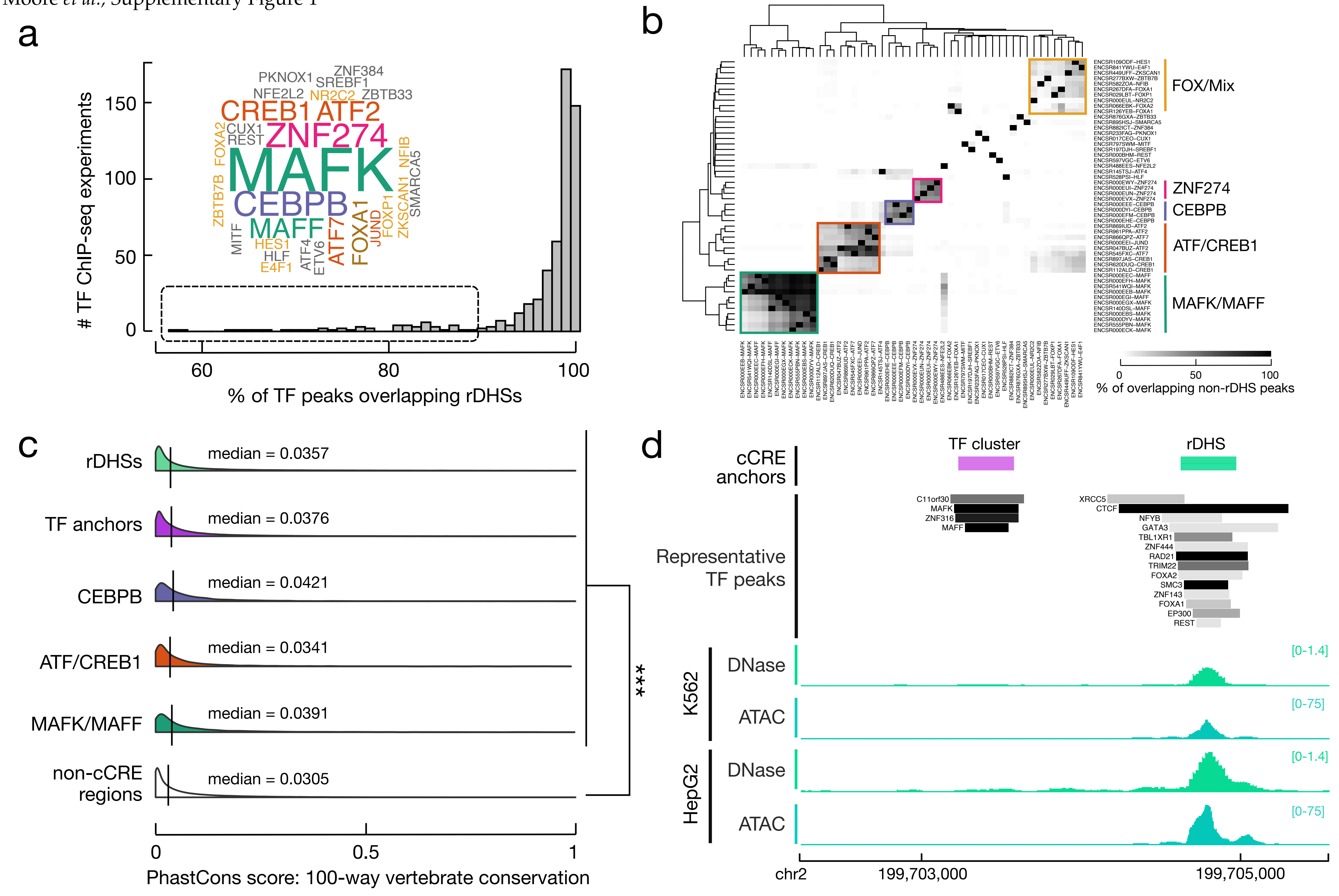
a
### TF ChIP-seq experiments
% of TF peaks overlapping rDHSs
MAFK
CEBPB
MAFF
ZNF274
CREB1 ATF2
ATF7
FOXA1
b
FOX/Mix
ZNF274
CEBPB
ATF/CREB1
MAFK/MAFF
% of overlapping non-rDHS peaks
c
rDHSs
median = 0.0357
TF anchors
median = 0.0376
CEBPB
median = 0.0421
ATF/CREB1
median = 0.0341
MAFK/MAFF
median = 0.0391
non-cCRE regions
median = 0.0305
***
PhastCons score: 100-way vertebrate conservation
d
TF cluster
rDHS
cCRE anchors
Representative TF peaks
K562
DNase
ATAC
HepG2
DNase
ATAC
[0-1.4]
[0-75]
chr2
199,703,000
199,705,000

**Supplementary Figure 1 | Expanding cCRE anchors to include transcription factor binding sites**

**a,** Histogram depicting the percentage of transcription factor (TF) peaks overlapping rDHSs. TFs with less than 90% overlap constituted a distinct set of TFs (depicted in the word cloud) such as MAFK, CEBPB, and ZNF274. Size of TF name is proportional to the number of TF experiments with less than 90% overlap. Full results can be found in **Supplementary Table 2e**. **b,** Heatmap depicting the overlap coefficient between TF peaks, which do not overlap rDHSs, between experiments with low rDHS overlap as defined in the dashed box in **a**. Five primary TF experiment clusters were defined using hierarchical clustering. **c**, Density plots depicting the average 100-way vertebrate alignment phastCons conservation score for rDHSs (green) and proposed TF anchors (purple), which are TF binding sites that do not overlap rDHSs. Three categories of TF anchors, as defined in **b**, are shown including CEBPB sites (blue), ATF/CREB1 sites (orange), and MAFK/MAFF sites (teal). Non-cCRE regions are shown in white. All groups were significantly more conserved than non-cCRE regions, supporting their inclusion in the Registry of cCREs. **d,** Example of rDHS (green) and TF anchor (purple). The rDHS has high chromatin accessibility in K563 and Hepg2 as measured by DNase-seq and ATAC-seq data. The TF cluster has low chromatin accessibility. Both sites have TF binding with the TF cluster having binding for MAFK, MAFF, ZNF316, and C11orf30.

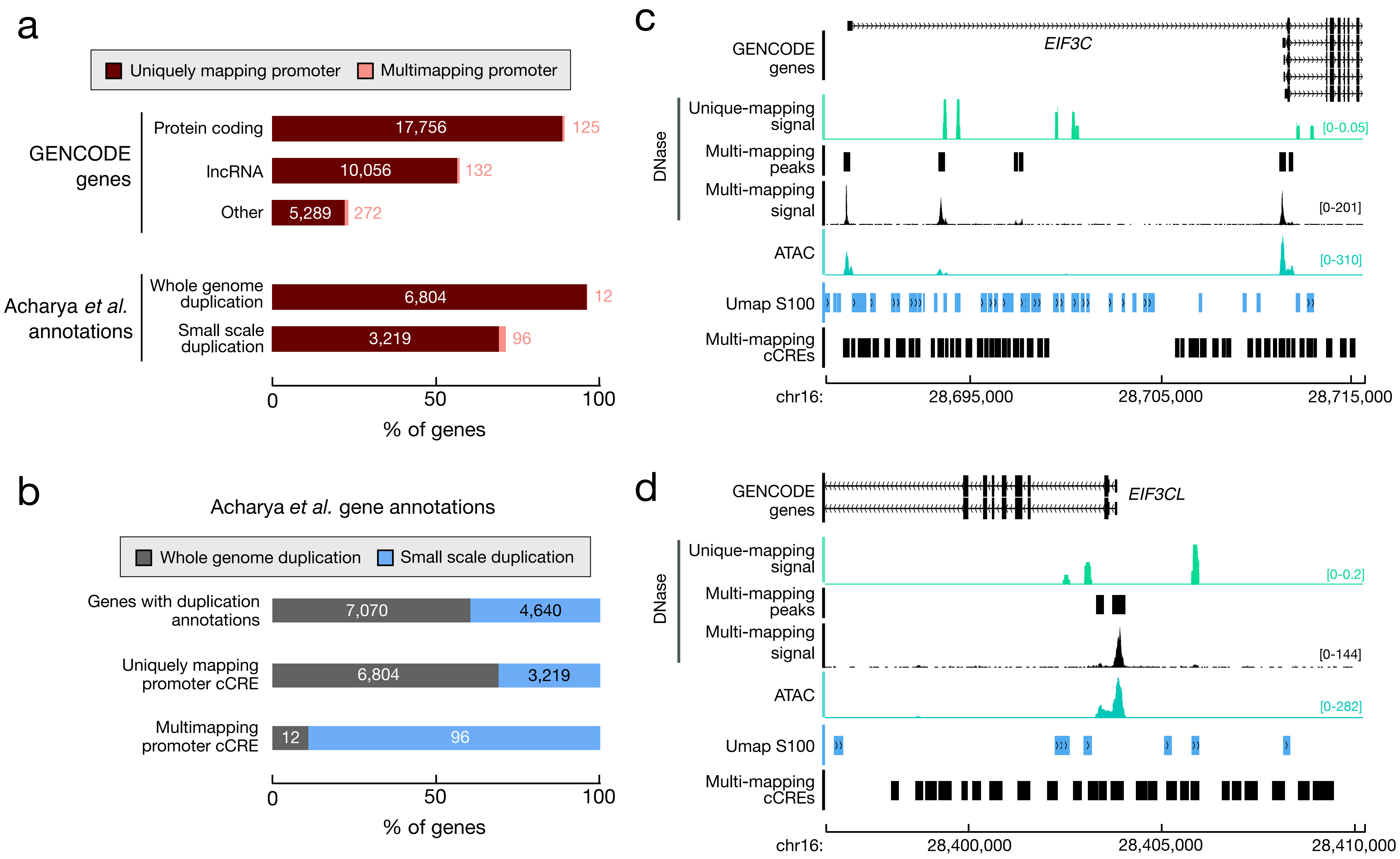
a
Uniquely mapping promoter
Multimapping promoter
GENCODE genes
Protein coding
17,756
125
lncRNA
10,056
132
Other
5,289
272
Acharya et al. annotations
Whole genome duplication
6,804
12
Small scale duplication
3,219
96
0
50
100
% of genes
b
Acharya et al. gene annotations
Whole genome duplication
Small scale duplication
Genes with duplication annotations
7,070
4,640
Uniquely mapping promoter cCRE
6,804
3,219
Multimapping promoter cCRE
12
96
0
50
100
% of genes
c
GENCODE genes
EIF3C
DNase
Unique-mapping signal
[0-0.05]
Multi-mapping peaks
Multi-mapping signal
[0-201]
ATAC
[0-310]
Umap S100
Multi-mapping cCREs
chr16:
28,695,000
28,705,000
28,715,000
d
GENCODE genes
EIF3CL
DNase
Unique-mapping signal
[0-0.2]
Multi-mapping peaks
Multi-mapping signal
[0-144]
ATAC
[0-282]
Umap S100
Multi-mapping cCREs
chr16:
28,400,000
28,405,000
28,410,000

**Supplementary Figure 2 | Including cCREs at multi-mapping loci identifies promoters for recently duplicated genes**

**a,** Barplots depicting the percentage of genes with annotated promoter cCREs. Genes are stratified based on GENCODE annotations (top) or classification by Archarya *et al*.[78]. Promoter cCREs are stratified by whether they arose from uniquely mapping cCREs (dark red) or multi-mapping cCREs. **b**, Barplots showing the proportion of genes originating from from whole genome duplication (gray) or more recent small scale duplication (blue) as annotated by Archarya *et al*. All annotated genes are shown (top) along with genes with uniquely mapping promoters (middle) and genes with multi-mapping promoters (bottom). **c**, Genome browser view of the *EIF3C* locus with the default unique-mapping DNase (green) and ATAC (teal) tracks. Multi-mapping DNase-seq signal is shown in black along with DHSs called from this signal. Umap S100 mapability tracks are shown in blue and multi-mapping cCREs are shown in black. **d**, Genome browser shot of the *EIF3CL* locus with the same tracks as defined in **c**.

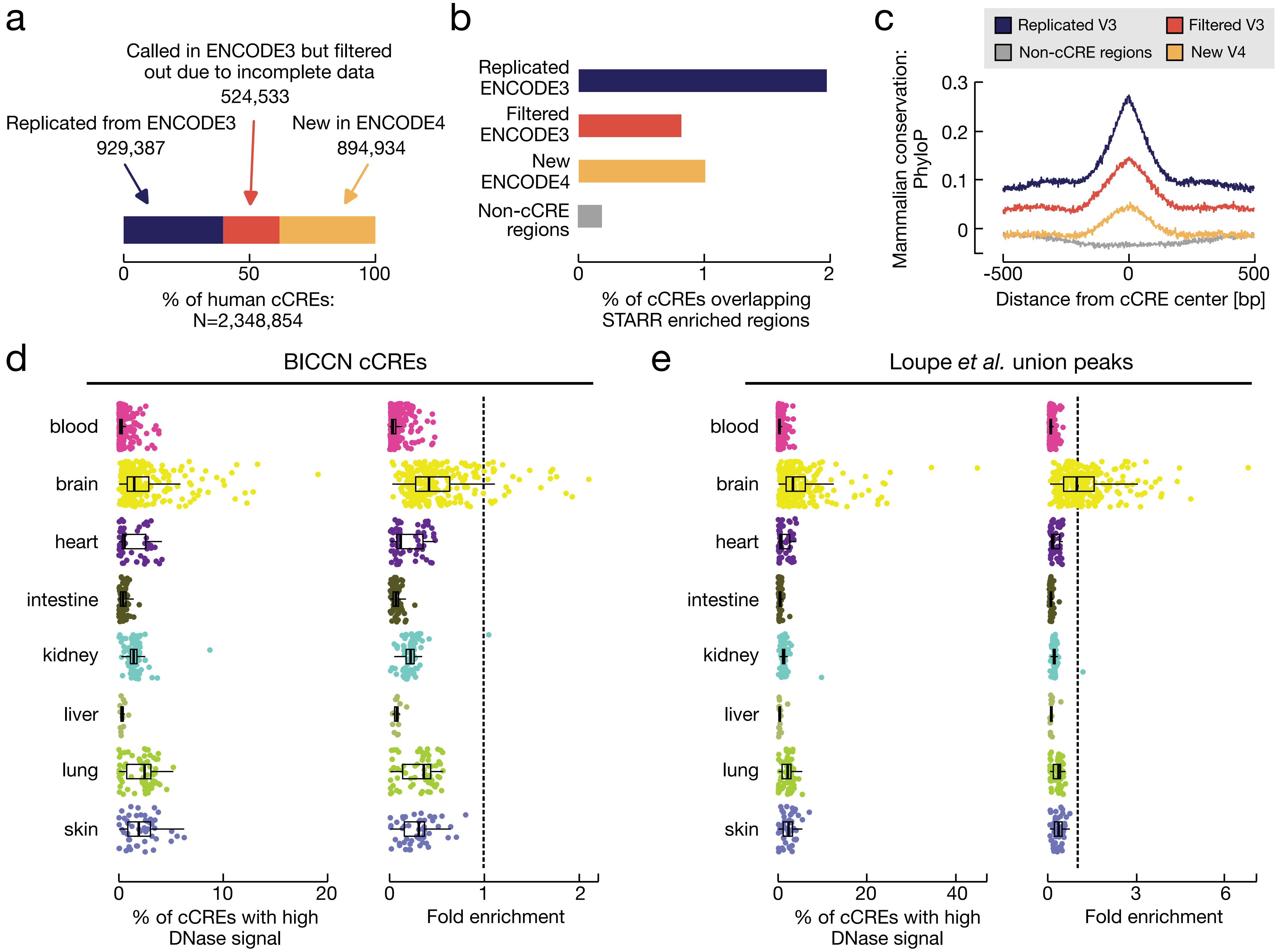
a
Called in ENCODE3 but filtered out due to incomplete data
524,533
Replicated from ENCODE3
929,387
New in ENCODE4
894,934
0
50
100
% of human cCREs:
N=2,348,854
b
Replicated ENCODE3
Filtered ENCODE3
New ENCODE4
Non-cCRE regions
0
1
2
% of cCREs overlapping STARR enriched regions
c
Replicated V3
Filtered V3
Non-cCRE regions
New V4
Mammalian conservation: PhyloP
0.3
0.2
0.1
0
-500
0
500
Distance from cCRE center [bp]
d
BICCN cCREs
e
Loupe et al. union peaks
blood
brain
heart
intestine
kidney
liver
lung
skin
0
10
20
% of cCREs with high DNase signal
0
1
2
Fold enrichment
0
20
40
% of cCREs with high DNase signal
0
3
6
Fold enrichment

#### Supplementary Figure 3 | Validation of the expansion of the Registry of cCREs

**a**, Barplot showing the percentage of human cCREs in the ENCODE4 Registry that were: previously in the ENCODE3 registry (blue), initially called in ENCODE3 but filtered out (orange), new to ENCODE4 (yellow). **b**, Barplots showing the percentage of cCREs (as defined in a) and size-matched non-cCRE regions (gray) that overlap K562 STARR-seq peaks. **c**, Line plots depicting average phyloP conservation scores calculated from 240-way Mammalian alignment for +/- 500 bp around each cCRE center stratifying by cCRE group as defined in **b**. **d**, Boxplots showing the percentage of ENCODE4-specific cCREs that overlap BICCN cCREs with high DNase signals in different biosamples, stratified by tissue/organ of origin. Each point represents an individual biosample. **e**, Boxplots showing the enrichment of ENCODE4-specific cCREs that overlap BICCN cCREs for DNase signal—compared to all cCREs—in different biosamples, stratified by tissue/organ of origin. **f**, Boxplots showing the percentage of ENCODE4-specific cCREs that overlap union peaks from Loupe *et al.*[33] with high DNase signals in different biosamples, stratified by tissue/organ of origin. Each point represents an individual biosample. **g**, Boxplots showing the enrichment of ENCODE4-specific cCREs that overlap union peaks from Loupe *et al*.[33] for DNase signal—compared to all cCREs—in different biosamples, stratified by tissue/organ of origin.

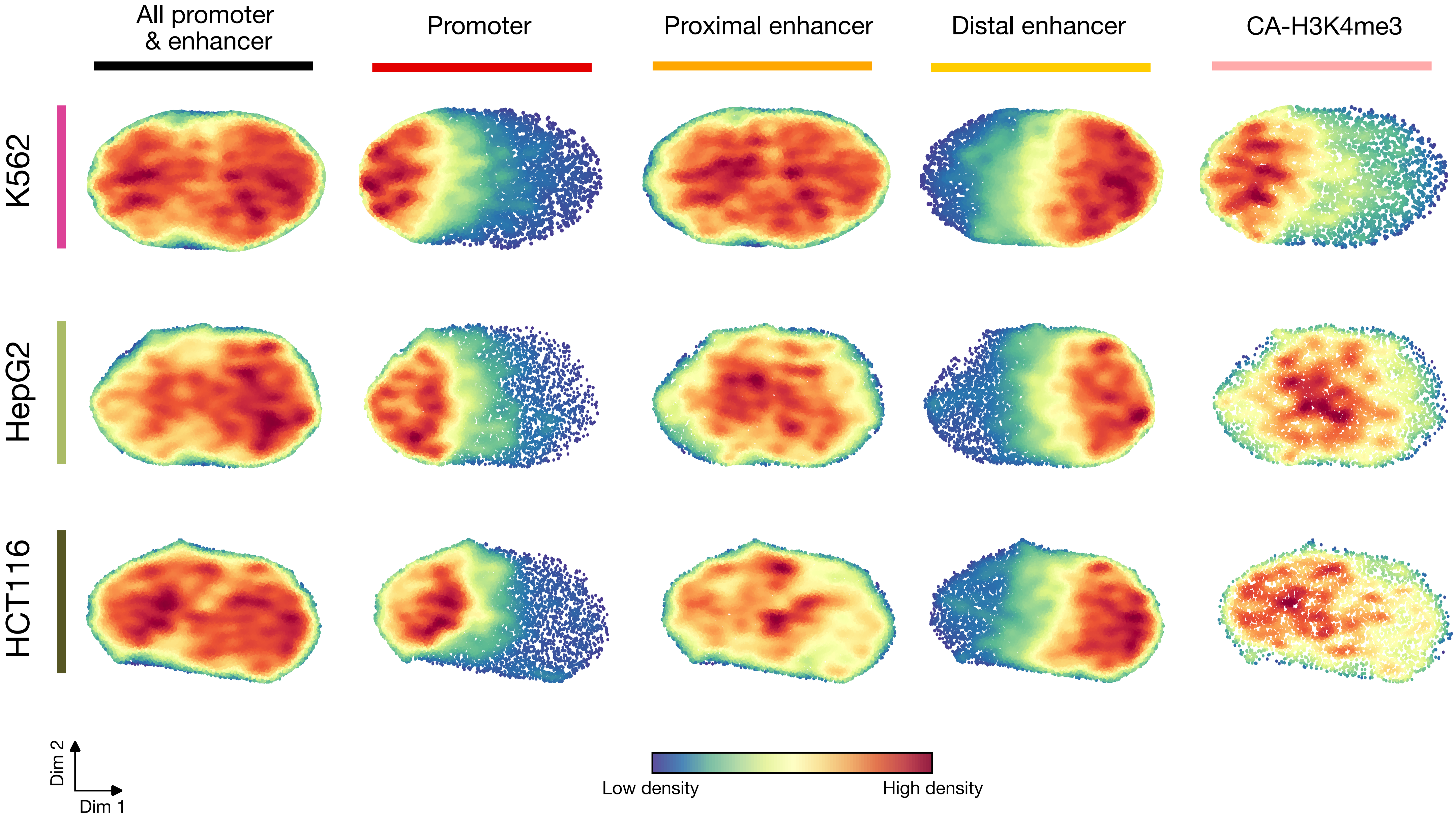

All promoter & enhancer
Promoter
Proximal enhancer
Distal enhancer
CA-H3K4me3
K562
HepG2
HCT116
Dim 2
Dim 1
Low density
High density

**Supplementary Figure 4 | Distinct sequence features of promoter and enhancer cCREs**

Density scatterplots showing the latent dimensions of cCRE sequence projected onto two dimensions by UMAP from variational autoencoders trained on three cell lines, K562 (top pink), HepG2 (middle green), and HCT116 (bottom olive). All promoter and enhancer cCREs are shown in column one followed by promoter (column 2) proximal enhancer (column 3), distal enhancer (column 4) and CA-H3K4me3 cCREs (column 5), in subsequent columns.

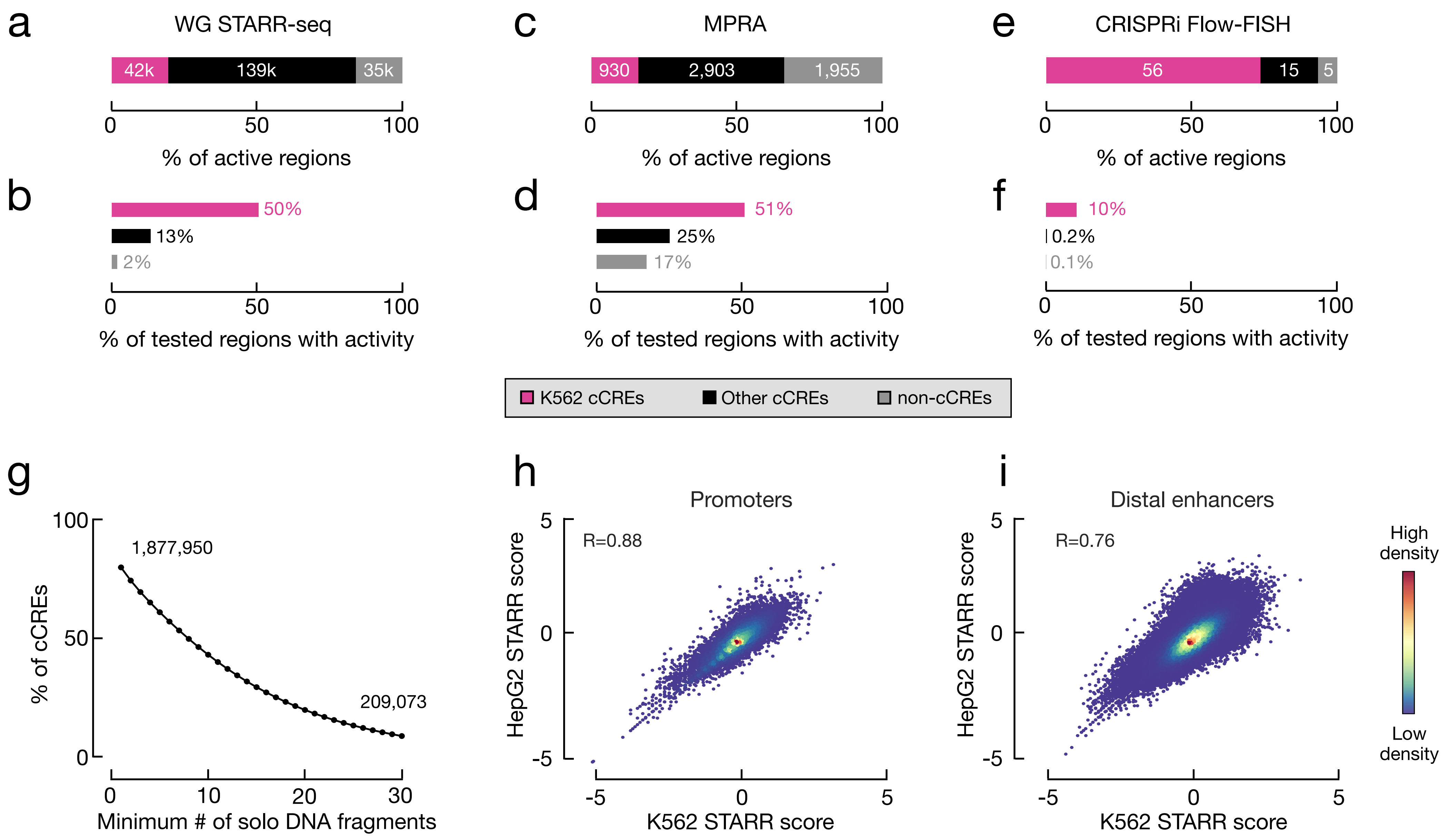
a
WG STARR-seq
42k
139k
35k
0
50
100
% of active regions
b
50%
13%
2%
0
50
100
% of tested regions with activity
c
MPRA
930
2,903
1,955
0
50
100
% of active regions
d
51%
25%
17%
0
50
100
% of tested regions with activity
e
CRISPRi Flow-FISH
56
15
5
0
50
100
% of active regions
f
10%
0.2%
0.1%
0
50
100
% of tested regions with activity
K562 cCREs
Other cCREs
non-cCREs
g
100
50
0
% of cCREs
1,877,950
209,073
0
10
20
30
Minimum # of solo DNA fragments
h
Promoters
R=0.88
5
0
-5
HepG2 STARR score
-5
0
5
K562 STARR score
i
Distal enhancers
R=0.76
5
0
-5
HepG2 STARR score
-5
0
5
K562 STARR score
High density
Low density

**Supplementary Figure 5 | cCREs are enriched for cell type-specific functional activity**

**a,** Barplot showing the percentage of active whole genome STARR-seq region (called peaks) that overlap K562 cCREs (pink), other cell type cCREs (black), and non-cCRE regions (gray). Numbers designate the number of regions in each category. **b,** Barplot depicting the percentage of cCRE or non-cCRE regions (as defined in **a**) that were tested in the assay that overlapped active whole genome STARR-seq regions. **c,** Barplot showing the percentage of active MPRA regions ($log_2$ fold change greater than one) that overlap K562 cCREs (pink), other cell type cCREs (black), and non-cCRE regions (gray). Numbers designate the number of regions in each category. **d,** Barplot depicting the percentage of cCRE or non-cCRE regions (as defined in **c**) that were tested in the assay that overlapped active MPRA regions. **e,** Barplot showing the percentage of active CRISPRi Flow-FISH regions (FDR < 0.05 ) that overlap K562 cCREs (pink), other cell type cCREs (black), and non-cCRE regions (gray). Numbers designate the number of regions in each category. **f,** Barplot depicting the percentage of cCRE or non-cCRE regions (as defined in **e**) that were tested in the assay that overlapped active CRISPRi Flow-FISH regions. **g**, Line plot depicting the percentage of cCREs with STARR scores as calculated by our CAPRA method for a single whole genome STARR-seq assay, as the minimum number of input DNA reads increases. **h**, Density scatterplot depicting the correlation between STARR scores calculated by CAPRA for promoter cCREs in K562 (x-axis) and HepG2 (y-axis). **i**, Density scatterplot depicting the correlation between STARR scores calculated by CAPRA for distal enhancer cCREs in K562 (x-axis) and HepG2 (y-axis).

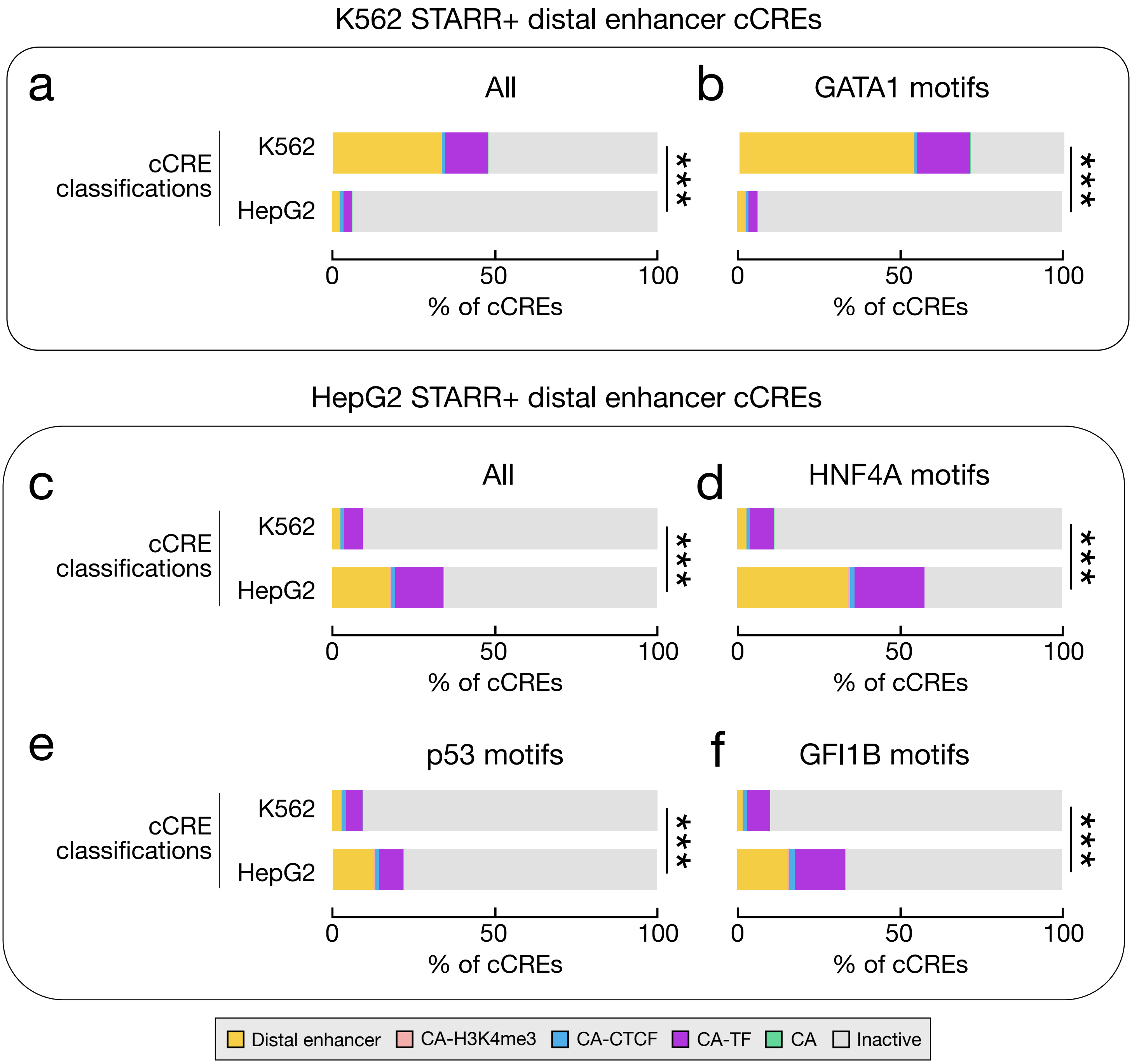
K562 STARR+ distal enhancer cCREs
a
All
b
GATA1 motifs
cCRE classifications
K562
HepG2
***
0
50
100
% of cCREs
HepG2 STARR+ distal enhancer cCREs
c
All
d
HNF4A motifs
e
p53 motifs
f
GFI1B motifs
Distal enhancer
CA-H3K4me3
CA-CTCF
CA-TF
CA
Inactive

**Supplementary Figure 6 | Cell type-specific STARR activity is concordant with cell type-specific chromatin signatures**

**a,** Barplots showing the percentage of distal enhancer cCREs with K562-specific STARR activity (K562 STARR+) belonging to each cCRE class called in K562 cells (using K562 chromatin signatures) and in HepG2 cells (using HepG2 chromatin signatures). **b,** Barplots showing the percentage of K562 STARR+ distal enhancer cCREs that overlap GATA1 motifs belonging to each cCRE class called in K562 cells and in HepG2 cells. **c,** Barplots showing the percentage of distal enhancer cCREs with HepG2-specific STARR activity (HepG2 STARR+) belonging to each cCRE class called in K562 cells (using K562 chromatin signatures) and in HepG2 cells (using HepG2 chromatin signatures). **d,** Barplots showing the percentage of HepG2 STARR+ distal enhancer cCREs that overlap HNF4A motifs belonging to each cCRE class called in K562 cells and in HepG2 cells. **e,** Barplots showing the percentage of HepG2 STARR+ distal enhancer cCREs that overlap p53 motifs belonging to each cCRE class called in K562 cells and in HepG2 cells. **f,** Barplots showing the percentage of HepG2 STARR+ distal enhancer cCREs that overlap GFI1B motifs belonging to each cCRE class called in K562 cells and in HepG2 cells. In all plots, the *** denotes $p$ less than 0.001 as calculated by a chi-square test.

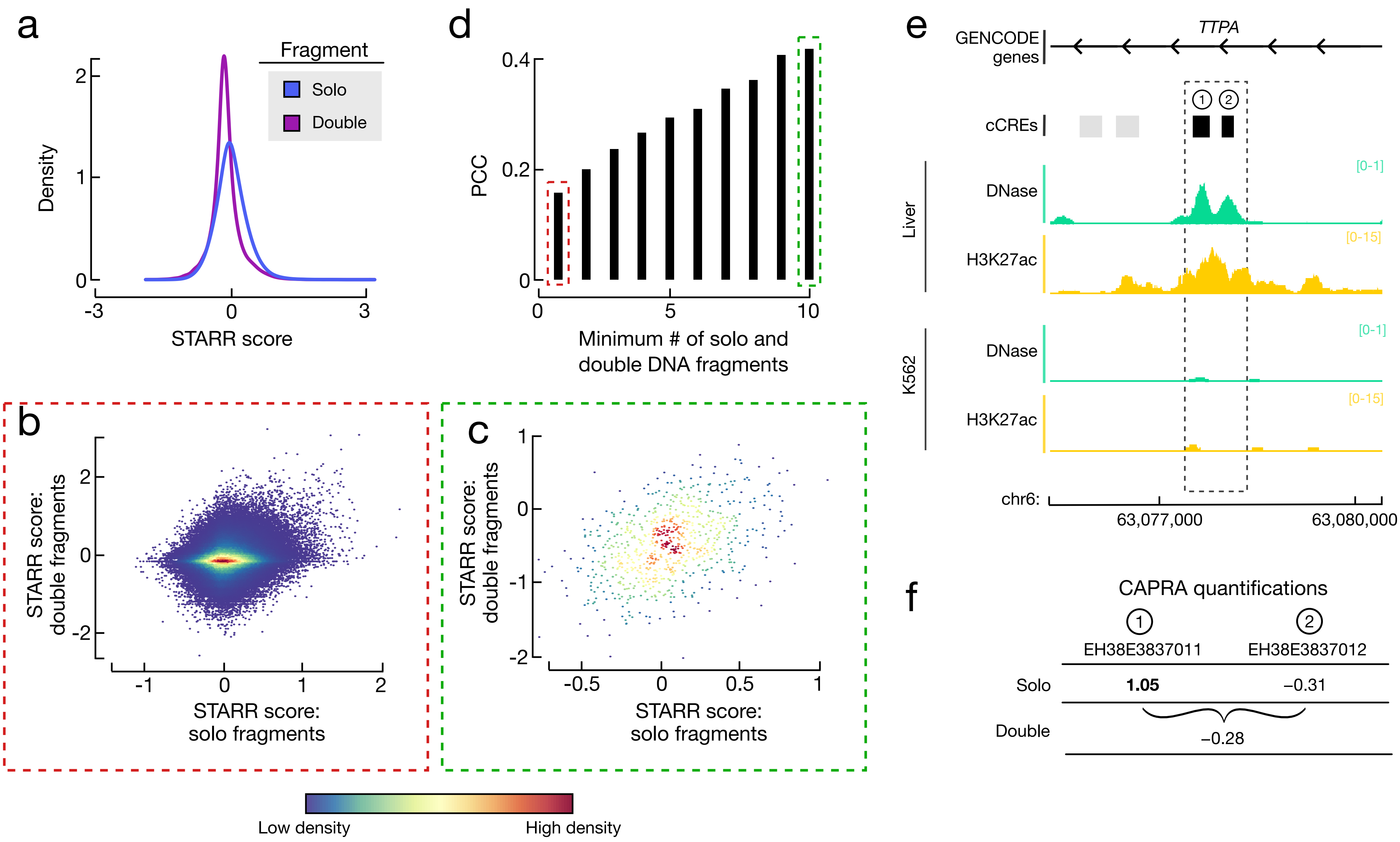

a
Fragment
Solo
Double
Density
STARR score
d
PCC
Minimum # of solo and double DNA fragments
e
TTPA
GENCODE genes
cCREs
Liver
DNase
[0-1]
H3K27ac
[0-15]
K562
DNase
[0-1]
H3K27ac
[0-15]
chr6:
63,077,000
63,080,000
b
STARR score: double fragments
STARR score: solo fragments
c
STARR score: double fragments
STARR score: solo fragments
Low density
High density
f
CAPRA quantifications
EH38E3837011
EH38E3837012
Solo
1.05
−0.31
Double
−0.28

#### Supplementary Figure 7 | Identifying combinatorial effects of cCRE activity

**a,** Density plots showing the distribution of STARR scores calculated by CAPRA from solo (blue) and double (purple) fragments. **b,** Barplots showing the pearson correlation coefficient (PCC) for cCRE pairs from the average STARR scores calculated from testing individual cCREs (solo DNA fragments) and the STARR scores calculated from testing multiple elements (double DNA fragments) as the threshold for the minimum number of input DNA fragments increases. Representative scatterplots for a cutoff of one (red box) and ten (green box) are shown in **c** and **d**. **c,** Density scatterplot of STARR scores for pairs of cCREs with a minimum of one DNA input fragment calculated by averaging quantifications from solo fragments or double fragments. **d,** Density scatterplot of STARR scores for pairs of cCREs with a minimum of ten DNA input fragments calculated by averaging quantifications from solo fragments or double fragments. **e,** Example of two cCREs (1: EH38E3837011 and 2: EH38E3837012) within the TTPA gene with lower than expected combinatorial effects. These cCRE have high DNase and H3K27ac signals in liver but low signals in K562 cells. **f,** STARR scores for EH38E3837011 and EH38E3837012 deriving from solo fragments (top) and double fragments (bottom). Scores with $p < 0.05$ are in bold.

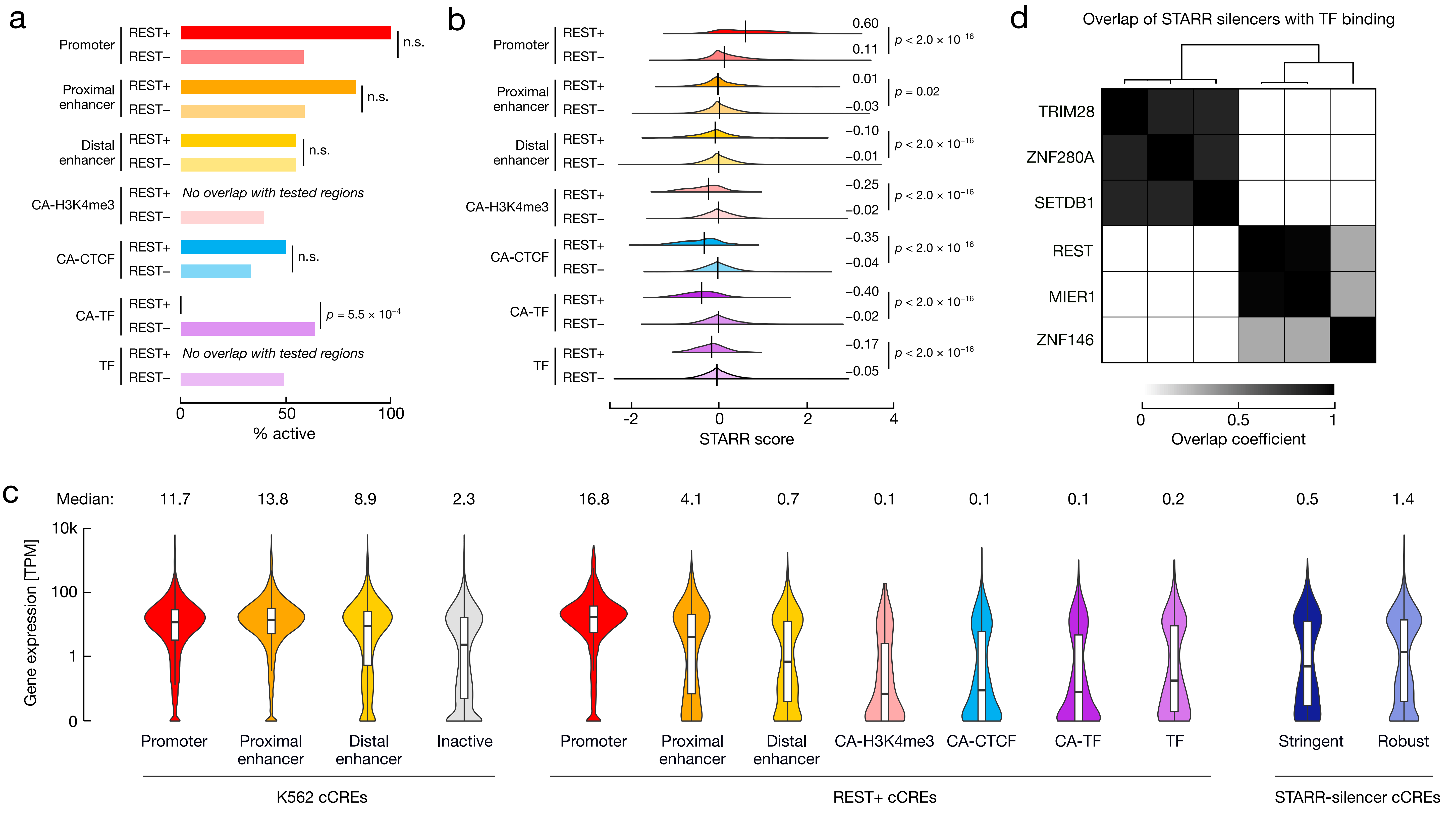
a
Promoter
Proximal enhancer
Distal enhancer
CA-H3K4me3
CA-CTCF
CA-TF
TF
REST+
REST−
n.s.
No overlap with tested regions
p = 5.5 × 10−4
0
50
100
% active
b
0.60
0.11
0.01
−0.03
−0.10
−0.01
−0.25
−0.02
−0.35
−0.04
−0.40
−0.02
−0.17
−0.05
p < 2.0 × 10−16
p = 0.02
-2
0
2
4
STARR score
d
Overlap of STARR silencers with TF binding
TRIM28
ZNF280A
SETDB1
REST
MIER1
ZNF146
0
0.5
1
Overlap coefficient
c
Median:
11.7
13.8
8.9
2.3
16.8
4.1
0.7
0.1
0.1
0.1
0.2
0.5
1.4
Gene expression [TPM]
10k
100
1
0
Promoter
Proximal enhancer
Distal enhancer
Inactive
CA-H3K4me3
CA-CTCF
CA-TF
TF
Stringent
Robust
K562 cCREs
REST+ cCREs
STARR-silencer cCREs

**Supplementary Figure 8 | Characteristics of silencer cCREs**

**a,** Barplots depicting the percentage of VISTA regions with enhancer activity in a transgenic mouse enhancer assay. Regions are stratified by cCRE class and overlap with REST ChIP-seq peaks (REST+). P-values are shown for Fisher's Exact Test. **b,** Density plots showing the distribution of K562 STARR scores for each class of cCRE stratified by REST binding (REST+). Vertical lines denote medians which are also displayed to the right of each distribution. P-values correspond to Wilcoxon tests. **c,** Nested violin-boxplots showing gene expression in K562 cells for the nearest gene of different classes and categories of cCREs. On the left are cCREs annotated in K562 based on K562-specific chromatin signatures including promoter, proximal enhancer, distal enhancer and inactive cCREs. In the middle are REST+ cCREs stratified by cell type agnostic classification. On the right are STARR-silencer cCREs stratified by threshold categories. Medians are shown above each distribution. **d,** Heatmap depicting the overlap of STARR silencers with binding sites for TRIM28, ZNF280A, SETDB1, REST, MIER1 and ZNF146 (DNA binding proteins enriched in STARR silencers, **Supplementary Table 8c**). Hierarchical clustering reveals two distinct clusters of STARR silencers based on protein binding.

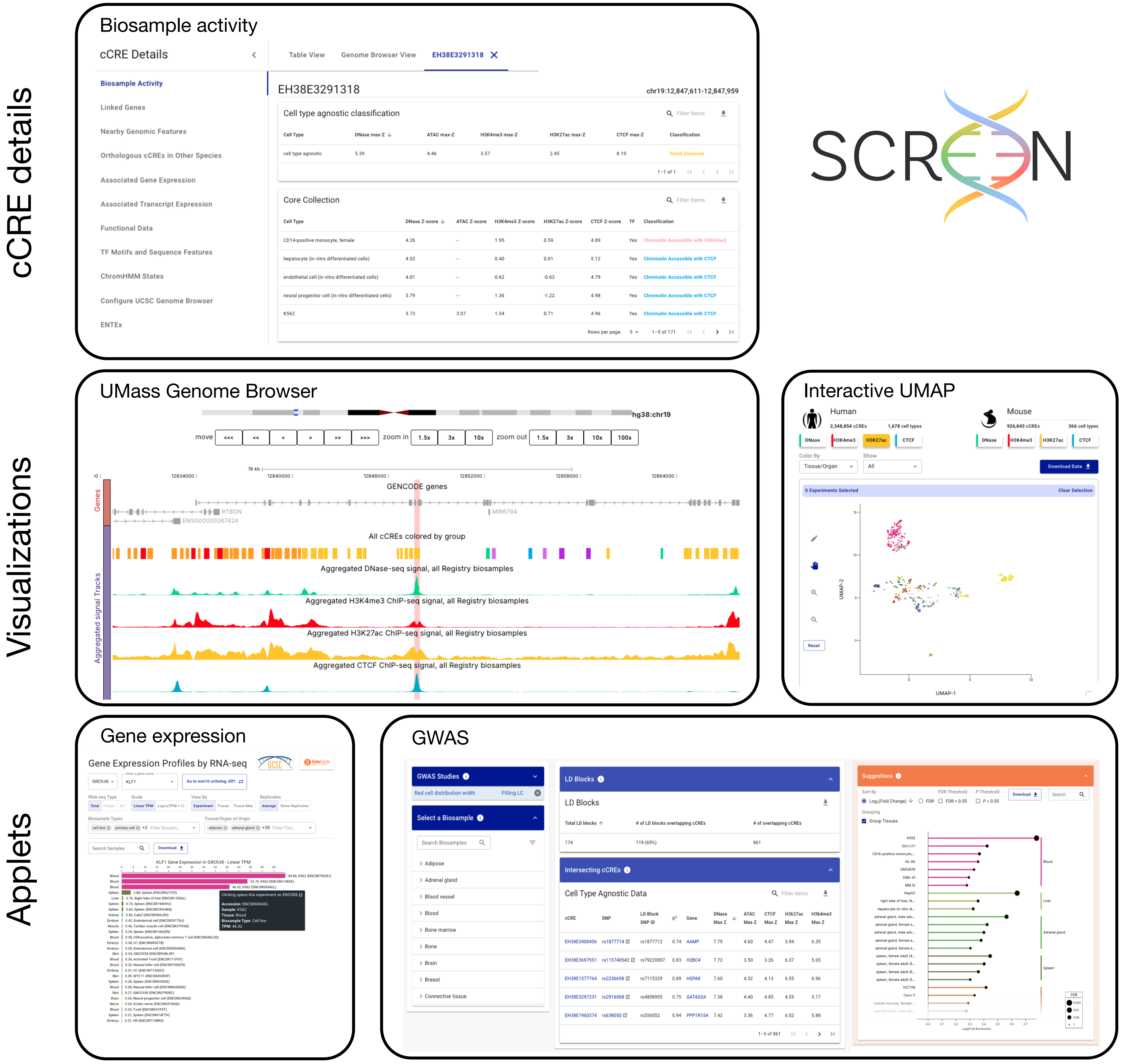
cCRE details
Visualizations
Applets
Biosample activity
cCRE Details
Table View
Genome Browser View
EH38E3291318
Biosample Activity
Linked Genes
Nearby Genomic Features
Orthologous cCREs in Other Species
Associated Gene Expression
Associated Transcript Expression
Functional Data
TF Motifs and Sequence Features
ChromHMM States
Configure UCSC Genome Browser
ENTEx
chr19:12,847,611-12,847,959
Cell type agnostic classification
Core Collection
SCREEN
UMass Genome Browser
hg38:chr19
GENCODE genes
All cCREs colored by group
Aggregated DNase-seq signal, all Registry biosamples
Aggregated H3K4me3 ChIP-seq signal, all Registry biosamples
Aggregated H3K27ac ChIP-seq signal, all Registry biosamples
Aggregated CTCF ChIP-seq signal, all Registry biosamples
Interactive UMAP
Human
Mouse
UMAP-1
UMAP-2
Gene expression
Gene Expression Profiles by RNA-seq
KLF1 Gene Expression in GRCh38 - Linear TPM
GWAS
GWAS Studies
Select a Biosample
LD Blocks
Intersecting cCREs
Cell Type Agnostic Data
Suggestions

#### Supplementary Figure 9 | SCREEN

SCREEN (Search candidate cis-Regulatory Elements by ENCODE) is an online resource for exploring and visualizing cCREs. Each cCRE has an element specific details page (top) that contains information about biosample activity and overlapping Encyclopedia annotations such as gene links, nearby gene expression, transcription factor binding sites, and conservation. SCREEN presents various visualizations (middle) including an embedded genome browser to enable seamless exploration along the genome (left) and interactive UMAPs to further explore relationships between biosamples and select subsets of samples. Finally, SCREEN hosts two applets gene expression (left) and GWAS (right).

Moore *et al.*, Supplementary Figure 10

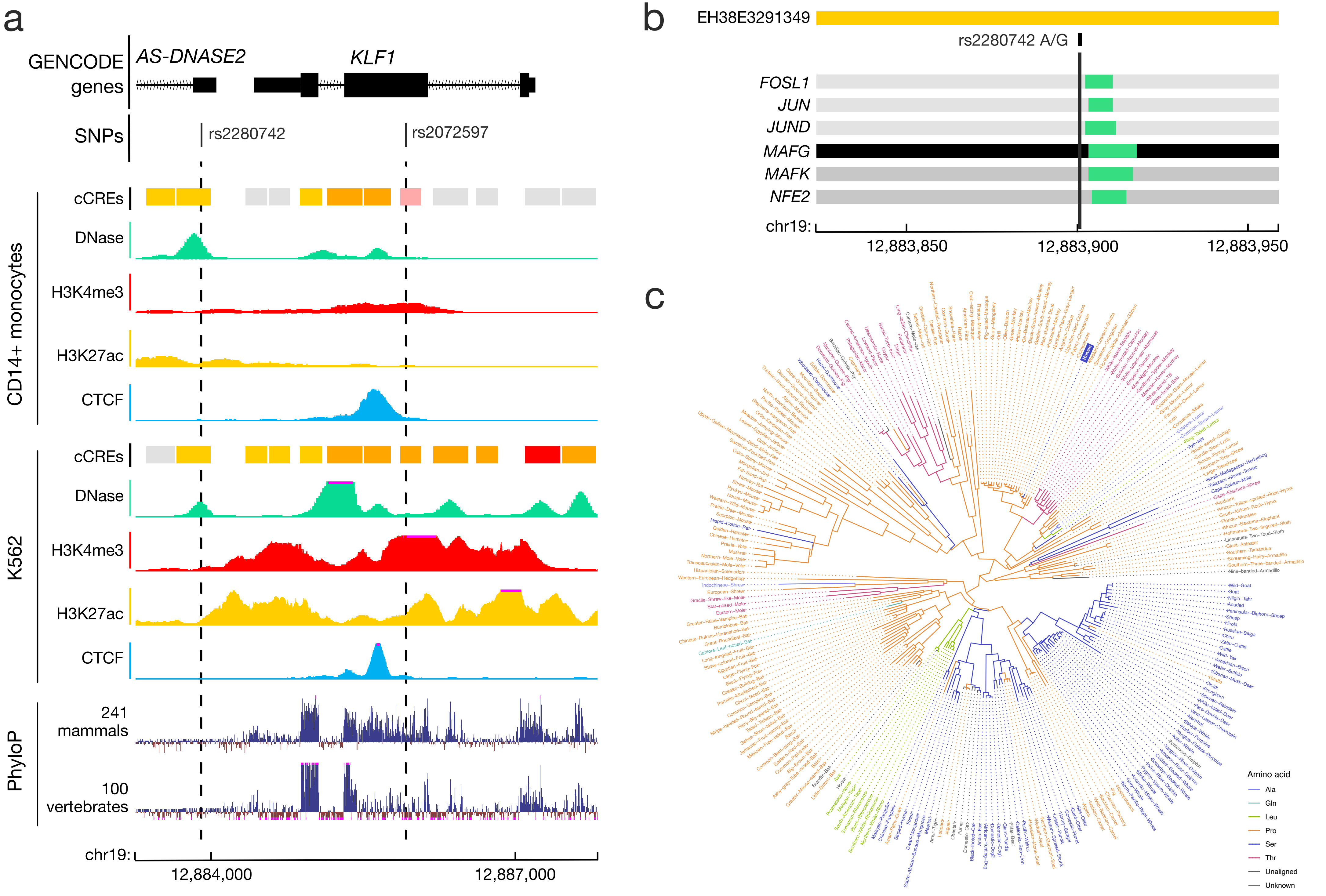

**Supplementary Figure 10 | Additional variants of interest at *RTBDN-MAST1* locus**

**a,** Genome browser view of two variants rs228072 and rs2072597 in high linkage disequilibrium with variants associated with Red Blood Cell traits. Data for two cell types, CD14+ monocytes and K562 cells are shown including cell type specific cCREs and signals for DNase (green), H3K4me3 (red), H3K27ac (yellow) and CTCF (blue). On the bottom are PhyloP conservation scores from mammalian and vertebrate alignments. **b,** Genome browser view of rs2280742 and overlapping cCRE EH38E3291349 (yellow) and transcription factor peaks with corresponding position of motif (green). **c,** Circular dendrogram displaying phylogeny for 240 placental mammals from the Zoonomia project showing amino acid usage at position 102 of *KLF1* based on each species reference genome.
